# Genetic incorporation of diverse non-canonical amino acids for histidine substitution

**DOI:** 10.1101/2025.11.04.686677

**Authors:** Anton Natter Perdiguero, Sandro Fischer, Alrika Ruth Lischke, Benjamin P. Manser, Alexandria Deliz Liang

**Author notes:** Correspondence should be addressed to A.D.L. These authors contributed equally (Anton Natter Perdiguero, Sandro Fischer).

## Abstract

Using genetic code expansion, canonical amino acid residues can be site-specifically substituted by non-canonical amino acids (ncAAs) with modified chemical properties. This technique has enabled detailed enzymatic studies, the design of enzymes that catalyze novel reactions, and the engineering of enzymes with improved function. In proteins, histidine can play versatile roles in catalysis including as an acid, a base, a nucleophile, and a coordinating ligand to a catalytic metal. However, the current scope of histidine-like ncAAs that can be incorporated is limited. Herein, we develop a toolkit consisting of nine new aminoacyl-tRNA synthetase/tRNA pairs for the site-specific genetic encoding of an expanded set of 12 new histidine-like ncAAs. The 12 ncAAs feature broadly tuned nitrogen p*K*_a_H, alternative heterocycles, and varying substitution patterns. We profile the substrate specificity of the developed aaRS/tRNA pairs and uncover many mutually orthogonal substrate specificities, which we validate for six combinations of dual encoded histidine-like ncAAs. We expect that the tools presented herein will be broadly applicable to study histidine residues in catalysis and to tune the properties of histidine residues for enzyme engineering and design.

## Introduction

Among canonical amino acids, histidine has the highest catalytic propensity and is often found in the active site of enzymes^1^. Histidine can play versatile roles in catalysis including as an acid, a base, a nucleophile, and a coordinating ligand to a catalytic metal. The imidazole side chain of histidine contains two nitrogen atoms that can participate in such chemistry—*N*^τ^ and *N*^π^ (see **Note 1** for further clarification of the nomenclature). Because histidine is structurally and chemically distinct from other canonical amino acids, its substitution with traditional mutagenesis methods is typically not a viable strategy to study or tune its role in catalysis. Alternatively, histidine residues can be substituted by histidine-like non-canonical amino acids (ncAAs). Several methods to incorporate histidine-like ncAAs into proteins exist. Global replacement of histidine has been leveraged to incorporate a selection of histidine-like ncAAs, but the ncAAs are incorporated indiscriminately at all histidine residues^2–5^. In contrast, genetic code expansion enables the site-specific incorporation of ncAAs into proteins^6^, allowing for more atomistic control. The most widely employed method for genetic code expansion uses engineered aminoacyl-tRNA synthetase (aaRS) and tRNA pairs to selectively incorporate ncAAs in response to a reassigned codon, typically the amber stop codon (TAG). This method enables expansion of the accessible chemical diversity in proteins and has been applied to carry out detailed enzymatic studies^7–9^, to design enzymes that catalyze novel reactions^10–14^, and to engineer enzymes with improved function^15–20^.

Despite the importance of histidine in catalysis and extensive efforts to engineer systems for incorporation of diverse histidine-like ncAAs^21^, the current accessible scope of such ncAAs remains limited^7,19,22–28^, particularly compared to tyrosine or lysine derivatives^29^. The most widely used histidine-like ncAA is *N*^π^-methyl-L-histidine (**πMH**, also referred to as NMH)^22,17,18,12,25,30^. The extensive use of this system in several different domains highlights the value of such histidine-line ncAAs. Expanding the scope available for histidine substitution to include different coordination chemistry, nucleophilicity, and acid/base reactivity would enable new modalities for studying and engineering of histidine-containing proteins. Additionally, a systematic assessment of the resulting engineered aaRS/tRNA pairs could illuminate features contributing to substrate recognition and enable incorporation of previously intractable ncAAs.

To address this challenge, we explored the genetic encoding of a large panel of diverse histidine-like ncAAs in *E. coli*. Through extensive engineering of various aaRS/tRNA pairs, we identify nine aaRS/tRNA systems for the site-specific genetic incorporation of a panel of 12 histidine-like ncAAs (Figure 1a). The 12 ncAAs feature broadly tuned nitrogen p*K*_a_H, alternative heterocycles, and varying substitution patterns. Additionally, within the set of engineered aaRS/tRNA pairs is the highly sought-after and elusive system for incorporation of *N*^*τ*^-methyl-L-histidine (**τMH**). We additionally profile the substrate scope of our nine evolved aaRS/tRNA pairs and discover orthogonal substrate combinations enabling the double-encoding of several histidine-like ncAAs. We expect that the tools presented herein will be broadly applicable to study histidine residues in catalysis and to tune the properties of histidine residues for enzyme engineering applications and enzyme design.

**Figure 1.**
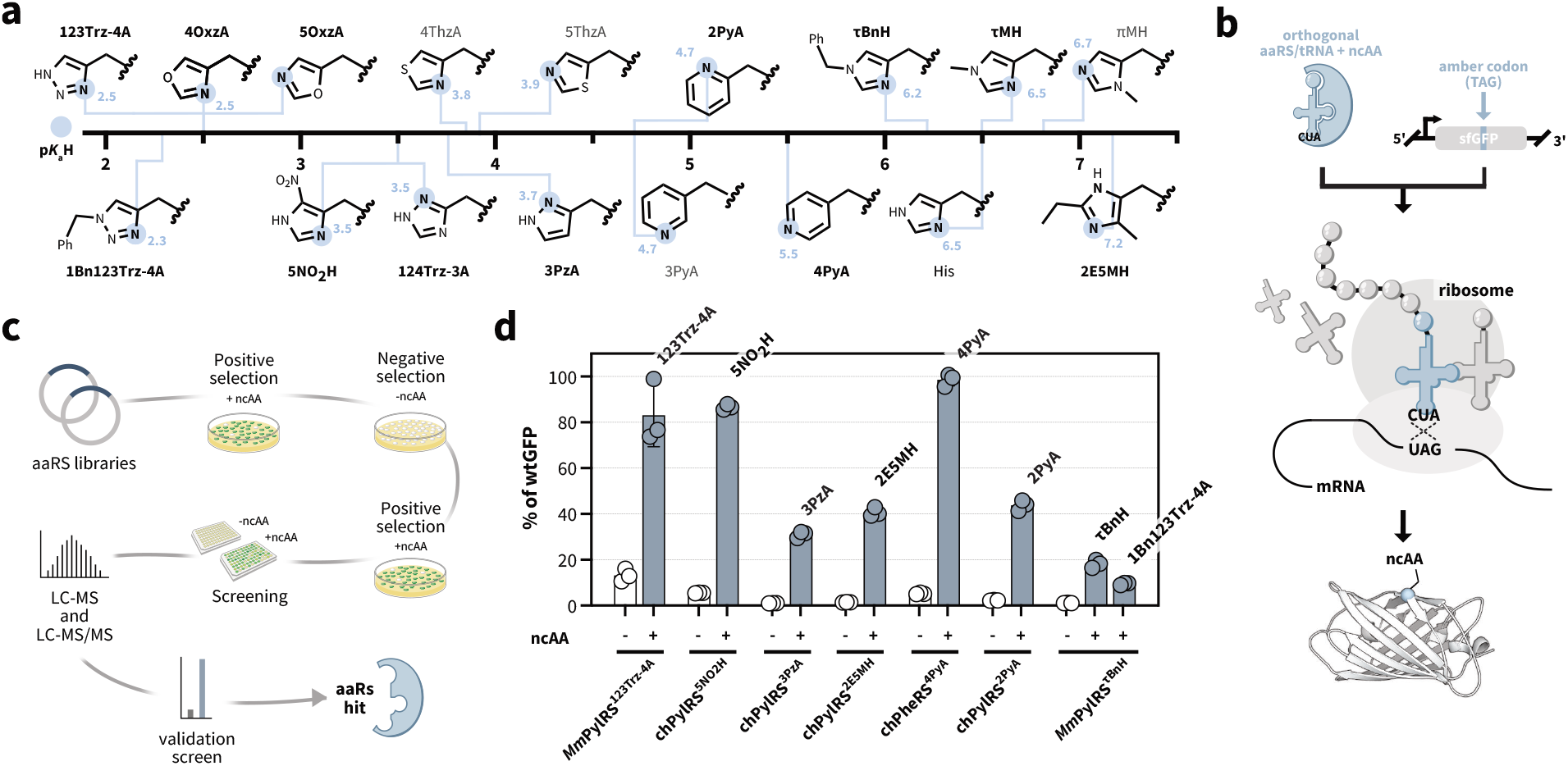
Targeting histidine-like ncAAs for site-selective incorporation by genetic code expansion. a) The p*K*_a_H scale of side chains of histidine and histidine-like ncAAs that were targets for this study. The names of ncAAs that have previously been genetically incorporated are grey, and the names of ncAAs that were newly incorporated in this work are shown in black and bold. b) Schematic of ncAA incorporation by amber suppression using genetic code expansion. c) General schematic for classical aaRS evolution, including aaRS library creation, positive selection to identify aaRSs capable of incorporating the ncAA or a canonical amino acid, negative selection to remove aaRSs that incorporate a canonical amino acid, a second positive selection to further enrich variants, a screening to quantify ncAA-dependent protein production, MS analysis to confirm the identity of the incorporated ncAA, and validation in an optimized plasmid designed for high production of the tRNA and the evolved aaRS. d) Amber suppression benchmarking of the first seven aaRS/tRNA pairs evolved in this study. Suppression efficiencies in sfGFP150_TAG_ with and without the respective ncAA. The ncAA concentrations that were used are listed in Supplementary Table 1. Data indicated as % of wt sfGFP (as measured by fluorescence, excitation at 480 nm and emission at 510 nm, and normalized to the optical density at 600 nm) in the absence or presence of different ncAAs as percentage of a wt sfGFP reference. The mean and standard deviation of 3 biological replicates is shown.

## Results

Towards expanding the scope for histidine mutagenesis, we identified several key properties that were missing from the genetic code expansion toolkit for histidine-like ncAAs: i) a diversity of heterocycles for fine-tuning reactivity, ii) histidine-like ncAAs with preserved *N*^π^ and *N*^τ^ and *increased* p*K*_a_H, iii) histidine-like ncAAs with preserved *N*^π^ and *N*^τ^ and *decreased* p*K*_a_H, and iv) *N*^τ^-substituted histidine derivatives, of which the only current ncAA contains a large photocleavable protecting group, making it excellent for photoactivation^26^ but preventing more widespread use in enzyme engineering, such as that seen for **πMH**. To address these limitations, we selected a panel of target ncAAs spanning a broad p*K*_a_H range that complement existing ncAAs and encompass a variety of desirable properties and substitutions (Figure 1a). This panel included new heterocyclic ncAAs that can enable alternative reactivity compared to imidazoles (**123Trz-4A, 4OxzA, 5OxzA, 124Trz-3A**, and **3PzA**), two stereoisomers of the useful **3PyA** to enable directional control of the reactive nitrogen (**2PyA** and **4PyA**), ncAAs containing substituted imidazole rings for tuning the p*K*_a_H (**5NO**_**2**_**H** and **2E5MH**), two new *N*^τ^-substituted histidine-like ncAAs (**1Bn123Trz-4A** and **τBnH**), and finally, the long-sought, but elusive **τMH**. From this target panel, we aimed to enable genetic code expansion (Figure 1b) through the development of orthogonal aaRS/tRNA pairs to encode each of these ncAAs site selectively (Figure 1c).

### Classical aaRS engineering approaches enable the incorporation of new ncAAs

For the target ncAAs, we carried out *de novo* selections on a set of site-saturation mutagenesis (SSM) libraries (Supplementary Table 2) generated from the PylRS from *Methanosarcina mazei* (*Mm*PylRS) and a chimeric PylRS variant that we previously evolved in an unrelated project, termed chPylRS^E7^, which derives from a fusion of the PylRSs from *Methanohalobium evestigatum* and *Methanosarcina mazei* (**Note 2**). As illustrated in Figure 1c, libraries were subjected to three alternating rounds of positive and negative selection against *Mm*tRNA^Pyl^_CUA_ in the presence or absence of the given ncAA, followed by screening of ncAA-dependent suppression in sfGFP150_TAG_ by fluorescence. The concentrations used for selections and screening can be found in Supplementary Table 1.

With 123Trz-4A, 5NO_2_H, 3PzA, and 2E5MH, distinct PylRS mutants were identified that incorporated each of the corresponding ncAAs (Supplementary Table 7). The derived synthetase for 3PzA exhibited low incorporation efficiency, but further optimization was achieved by replacing its N-terminal domain with that of *Mb*PylRS carrying the “IPYE” mutations^31^. Transfer of the engineered pairs onto our optimized plasmid for genetic code expansion application (pOS1T) afforded suppression systems that incorporate the given ncAA into sfGFP150_TAG_ with incorporation efficiencies ranging between 30% and 85% compared to wt sfGFP (Figure 1d). Incorporation of the ncAAs was confirmed by LC-MS analysis of intact sfGFP150_TAG_ and LC-MS/MS of trypsinized sfGFP150_TAG_ (Supplementary Figures 2a-d and 3a-d, respectively). Using LC-MS/MS analysis, we searched for the desired ncAA modification and also possible amino acids incorporated in response to the amber stop codon to provide an extra analysis of incorporation fidelity. For 123Trz-4A, 3PzA, and 2E5MH, no misincorporation was observed. In contrast, for 5NO_2_H, we observed very weak signals corresponding to incorporation of phenylalanine and glutamine, which were not detected by intact mass analysis. Based on comparison of the sfGFP production in the presence and absence of the ncAA, the intact mass spectra, and the low intensity of the misincorporation signals, our results strongly suggest that the misincorporation is very low, with a purity greater than 97% (Supplementary Tables 9 and 10). Although LC-MS/MS searches for misincorporation are not commonly reported in genetic code expansion efforts, these results highlight the importance of robust MS/MS analysis in rigorously characterizing incorporation fidelity. Based on this finding, we conducted LC-MS/MS misincorporation analysis for the previously reported ncAA-aaRS/tRNA systems explored herein and our newly evolved ncAA-aaRS/tRNA systems to understand and approximate the extent of low-level misincorporation in these systems (Supplementary Table 10).

The final aaRS/tRNA pair found through the classical approach was for 4PyA. We envisioned that this ncAA might be incorporated through engineering of a recently described chimeric pair derived from a fusion between a pyrrolysyl and phenylalanine system (chPheRS/3C11-chPheT_CUA_)^32,33^. Through screening a set of SSM libraries (Supplementary Table 5), we identified a variant carrying the mutations Q365N and A507S (chPheRS^4PyA-v0^, Supplementary Table 7). This mutant was further improved by three rounds of error-prone PCR followed by positive selection and screening. The final variant (chPheRS^4PyA^, Supplementary Table 7) contains 12 additional coding mutations throughout both the N- and C-terminal domains of the chPheRS and has an approximately 21-fold activity improvement compared to chPheRS^4PyA-v0^ (Supplementary Figure 5). Transfer of the chPheRS^4PyA^/3C11-chPheT_CUA_ pair into a pOS1T plasmid afforded a suppression system that incorporates 4PyA with low background activity and wt-like incorporation efficiency (Figure 1d). Incorporation of 4PyA was confirmed by LC-MS analysis of intact sfGFP150_TAG_ and LC-MS/MS analysis of trypsinized sfGFP150_TAG_ (Supplementary Figure 2g and 3g). The improvements with 4PyA highlight the value of random mutagenesis for improving activity from a weakly active starting point. We note that while our work was in progress, a report was published describing incorporation of 4PyA using polyspecific *Methanocaldococcus jannaschii* tyrosyl tRNA synthetase and *Methanogenic archaeon ISO4-G1* PylRS (*G1*PylRS) mutants^34^, albeit at lower efficiencies (<20% of wt sfGFP).

### Panel screening can be leveraged to identify aaRS/tRNA pairs for new ncAAs

For the remaining ncAAs within the target set, we screened an in-house panel of aaRS/tRNA pairs, which consists of previously reported aaRS/tRNA pairs and some unreported pairs that have been derived for various ongoing projects. From this panel, we evaluated sfGFP150_TAG_ production in the presence and absence of the target ncAAs, and we were able to identify aaRS/tRNA pairs for three additional ncAAs: 2PyA, τBnH, and 1Bn123Trz-4A .

In the case of 2PyA, two reported synthetases for the incorporation of ortho-substituted phenylalanine derivatives^35,36^ displayed promiscuous activity with 2PyA. However, the activity was low, and the background in the absence of 2PyA was high (Supplementary Figure 4). Thus, additional classical engineering was required to achieve selective incorporation. Mutations of two residues located near the ncAA side chain—N346 and C348—often impart important changes in substrate recognition but are insufficient to enable robust incorporation. Thus, to optimize the incorporation of 2PyA further, we constructed libraries of chPylRS^E7^ in which N346 and C348 were fixed to mimic these mutations in the weakly active aaRS/tRNA pairs and an additional five positions within the ncAA binding site were randomized by SSM (Supplementary Table 7). Through screening and selection, we identified a more active and selective mutant, chPylRS^2PyA^ (Supplementary Table 3). Transfer of the chPylRS^2PyA^/*Mm*tRNA^Pyl^_CUA_ pair into a pOS1T plasmid afforded a suppression system that incorporates 2PyA with low background activity and approximately 50% yield compared to wt sfGFP (Figure 1d). Incorporation of 2PyA was confirmed by LC-MS analysis of intact sfGFP150_TAG_ and LC-MS/MS analysis of trypsinized sfGFP150_TAG_ (Supplementary Figure 2f and 3f).

With τBnH and 1Bn123Trz-4A, we found that both of these ncAAs were incorporated by an unreported *Mm*PylRS mutant from the panel (hereafter *Mm*PylRS^τBnH^, Supplementary Table 7). Expression from sfGFP150_TAG_ yielded modest incorporation efficiencies of approximately 20% and 10%, respectively. For both ncAAs, there was very low background suppression in the absence of ncAA (Figure 1d). Additionally, incorporation of the desired ncAAs was confirmed by LC-MS analysis of intact sfGFP150_TAG_. LC-MS/MS of trypsinized sfGFP150_TAG_ exhibited low misincorporation levels and indicated that the fidelity was higher for τBnH than 1Bn123Trz-4A (Supplementary Figures 2h and 3h and Supplementary Tables 9 and 10). Based on the low background, further engineering was not conducted.

### Substrate profiling reveals surprising substrate specificities

From our classical aaRS/tRNA engineering method, we were able to incorporate several new ncAAs, but several high-value targets remained challenging. Thus, with a panel of aaRS/tRNA pairs that direct the incorporation of diverse histidine analogs in hand, we opted to profile the substrate specificity of different ncAA-aaRS combinations. We expected that such a panel might reveal some incorporation trends and identify aaRS/tRNA pairs with polyspecificity for incorporation of our remaining ncAAs. We included several previously reported aaRS/tRNA pairs that have been described for the incorporation of histidine analogs: *Mm*PylRS^IFGFF^/*Mm*tRNA^Pyl^, which carries mutations grafted from *Mb*PylRS^IFGFF 22^, and *G1*PylRS^MIFAF^/*G1*tRNA^ΔNPyl 28^, which carries mutations from *Ma*PylRS^MIFAF 23^, both of which were developed for πMH incorporation. Additionally, we included the *Mm*PylRS^FLF^/*Mm*tRNA^Pyl^ and *Mm*PylRS^8_2^/*Mm*tRNA^Pyl^ pairs described for 3PyA incorporation^21,27^, the *Mb*PylRS^4Thz^/*Mm*tRNA^Pyl^ described for 4ThzA incorporation^25^ and the *Mm*PylRS^QF^/*Mm*tRNA^Pyl^ and *Mm*PylRS^7_1^/*Mm*tRNA^Pyl^ pair described for 3-thienyl-L-alanine (3ThA) incorporation^21,25^. We tested each suppressor system with the ncAAs in Figure 1a as well as 2-thienyl-L-alanine (2ThA) and 3ThA. As a reporter, we used sfGFP150_TAG_, and we quantified the sfGFP production in the presence or absence of the given ncAA using the concentrations reported in Supplementary Table 1 and a standardized plasmid system (Figure 2a, and Supplementary Figure 6).

**Figure 2.**
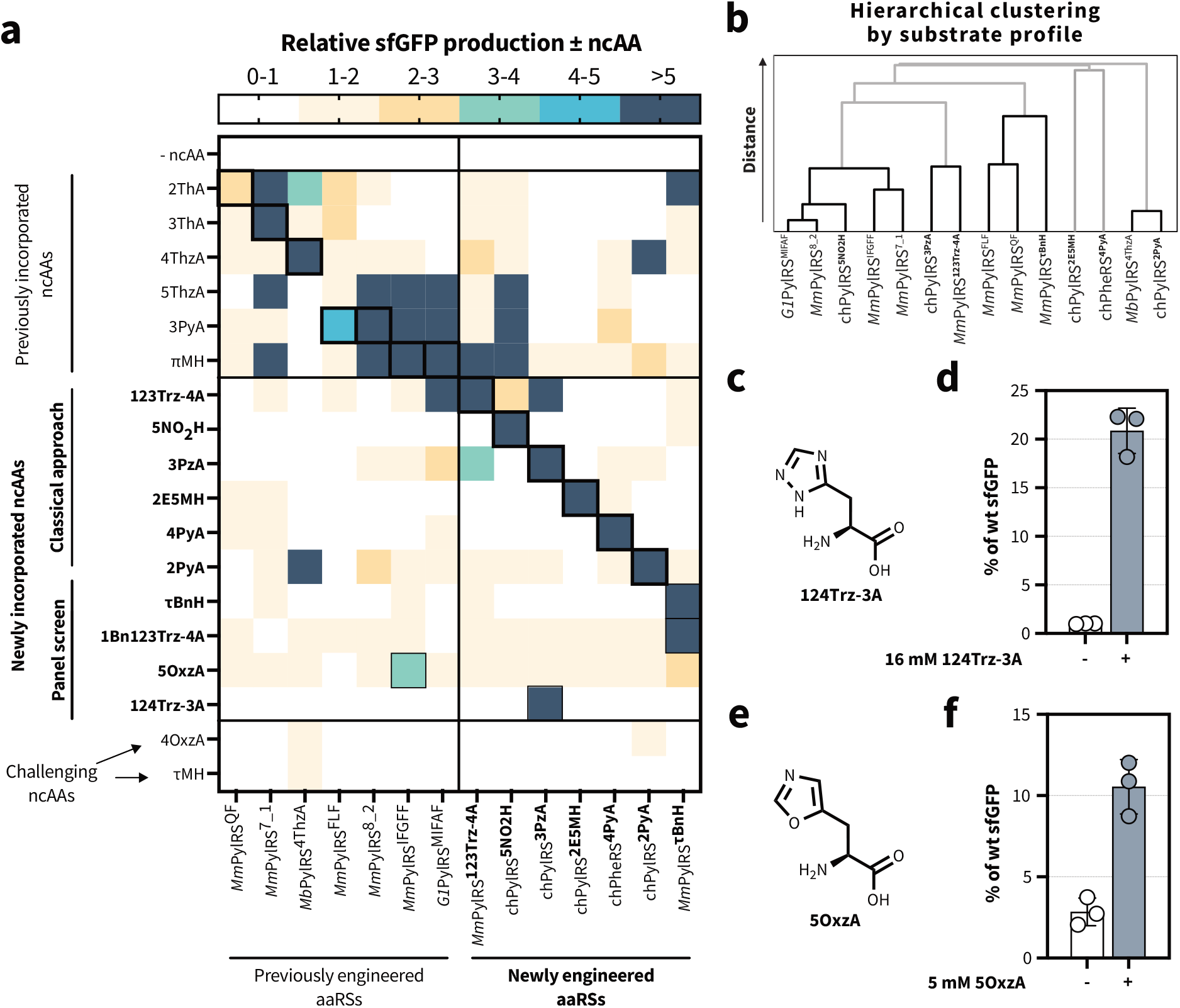
aaRS-ncAA substrate profiling and incorporation of additional histidine-like ncAAs by polyspecificity. a) Substrate specificity profiling for different aaRS-ncAA combinations. Production of sfGFP150_TAG_ was carried out for different aaRS/tRNA pairs in the absence or presence of the indicated histidine-like ncAA. The relative sfGFP production in the presence of the ncAA vs absence was determined. Each row shows a distinct ncAA, and each column a distinct aaRS. The entry for the “cognate” ncAA of the aaRS/tRNA pair is outlined with a thick black border. The ncAA concentrations used are reported in Supplementary Table 1. The mean of 2-3 biological replicates are shown. b) Hierarchical clustering of aaRSs by substrate profile indicating the similarity of aaRSs based on their substrate preferences. In the dendrogram, the distance at which the connections are made indicates the level of similarity. Connections made at small distances are more similar, whereas connections made at larger distances are more dissimilar. c) Chemical structure of 124Trz-3A. d) Incorporation of 124Trz-3A into sfGP150_TAG_ using the chPylRS^3PzA^ system. e) Chemical structure of 5OxzA. f) Incorporation of 5OxzA into sfGP150_TAG_ using the *Mm*PylRS^IFGFF^ system. Data indicated as sfGFP production (+ncAA/-ncAA) shows normalized sfGFP production with and without a given ncAA. Data indicated as % of wt sfGFP (as measured by fluorescence, excitation at 480 nm and emission at 510 nm, and normalized to the optical density at 600 nm) in the absence or presence of different ncAAs as percentage of a wt sfGFP reference. For panels d and f, the mean and standard deviation of 3 biological replicates is shown.

Within the panel, we observed high substrate specificity for aaRS/tRNA pairs with their *cognate* ncAAs (meaning the ncAAs for which they were originally evolved) compared to canonical amino acids. Additionally, for most aaRS/tRNA pairs, we find that they can incorporate other ncAAs in addition to the cognate substrate. This observation, termed polyspecificity, is a common result for aaRSs obtained from directed evolution^37,38^. As expected, polyspecificity is often found for ncAAs that are chemically highly similar to the cognate substrate. Notably, there was significantly less polyspecificity within the newly engineered aaRS/tRNA pairs reported herein, compared to previously described aaRS/tRNA pairs. The reason for this is unclear but could reflect ease of incorporation from previously described histidine-like ncAAs, differences in application of the classical engineering approach, or lower chemical similarity of the new ncAA set, which was designed to fill gaps in the existing histidine-like ncAA set.

Despite the generally higher specificity observed, for highly similar chemical structures, polyspecificity can still be observed. For example, two distinct synthetases were evolved for 123Trz-4A and 3PzA, *Mm*PylRS^123Trz-4A^ and chPylRS^3PzA^, respectively. Based on the chemical similarity between these ncAAs, we expected both synthetases could be polyspecific for 123Trz-4A and 3PzA, which was confirmed experimentally. Notably, *Mm*PylRS^123Trz-4A^ has a broader substrate scope, also efficiently incorporating 4ThzA and πMH, substrates for which chPylRS^3PzA^ has no activity. Interestingly, the same libraries were used to derive *Mm*PylRS^123Trz-4A^ as chPylRS^3PzA^, but *Mm*PylRS^123Trz-4A^ was not found in the 3PzA screening. This result points to the stochastic nature of multi-step selection and screening for such large libraries, beyond even the randomness of large library creation itself. Moreover, we observed that πMH was incorporated by many different aaRS/tRNA pairs, and both *Mm*PylRS^123Trz-4A^ and chPylRS^5NO2H^ incorporated πMH more efficiently than their cognate substrate. Analogously, chPylRS^5NO2H^ also incorporates 3PyA at wt-like levels, more efficiently than the previously reported *G1*PylRS^MIFAF^ and chPylRS^FLF^. Collectively, these results also highlight the ability of many different evolutionary solutions to efficiently address a single enzyme engineering goal.

Overall, we observe many non-obvious substrate selectivities, with some aaRS variants capable of distinguishing substrates with highly similar structures and some that seem to act more as generalists. Towards mapping these differences more systematically, we performed hierarchical clustering analysis based on substrate profiles (Figure 2b). The hierarchical analysis revealed that the aaRSs can be separated into distinct clades based on their substrate profiles. Both chPylRS^2PyA^ and *Mb*PylRS^4Thz^ have highly similar substrate profiles that are also the most different from the other aaRSs. Additionally, both chPheRS^4PyA^ and chPylRS^2E5MH^ are outliers with unique profiles within the dataset. In the case of chPheRS^4PyA^, this observation is perhaps not so surprising given that this system is derived from a very different aaRS sequence than the others. In terms of future engineering campaigns, we expect that selecting several parent sequences from disparate substrate profile clades may improve the chances of successful aaRS evolution. Beyond describing these substrate profiles, the ability to predict the substrate tolerance of an aaRS would be of extremely high value; thus, we studied if incorporation parameters were correlated with computationally predicted binding affinities. However, we found that the predicted binding affinities—even when accounting for logP values—are a very poor indicator for aaRS substrate preference within this set of ncAAs (Supplementary Figure 7). These results highlight the complexity of aaRS activity and selectivity, particularly for relatively small, polar ncAAs with highly similar structures.

In addition to exploring substrate preferences, we were able to identify two additional pairs with robust suppression efficiencies for new ncAAs. In the case of 124Trz-3A, we discovered that this ncAA is a substrate for chPylRS^3PzA^. Indeed, production of sfGFP in the presence of elevated 124Trz-3A concentrations showed incorporation of 124Trz-3A with an approximately 20% yield of wt sfGFP and a low background in the absence of 124Trz-3A. (Figure 2e). The incorporation was confirmed by LC-MS analysis of intact sfGFP150_TAG_, and LC-MS/MS analysis of trypsinized sfGFP150_TAG_ indicated low levels of misincorporation (Supplementary Figures 2e and 3e, Supplementary Table 9 and 10). We also identified 5OxzA as a substrate for *Mm*PylRS^IFGFF^, albeit with low protein yields compared to wt sfGFP (∼12%, Figure 2g). Incorporation of 5OxzA was confirmed by LC-MS analysis of intact sfGFP150_TAG_, and LC-MS/MS analysis of trypsinized sfGFP150_TAG_ also indicated low misincorporation (Supplementary Figure 2j and 3j, Supplementary Table 9 and 10). In all, from the comprehensive substrate profiling, we identified two additional histidine-like ncAAs that are accepted by existing aaRS/tRNA pairs with modest efficiencies.

### Substrate profiling can be leveraged to conduct smarter engineering

From the aaRS-ncAA substrate profiling, we carefully examined the incorporation profiles for ncAAs for which we had failed to identify an aaRS/tRNA pair from the methods above, namely 4OxzA and τMH. We identified two aaRS/tRNA pairs with minute (< 1.2-fold) but reproducible activity above background (*Mb*PylRS^4ThzA^ and chPylRS^2PyA^, respectively). From the aaRS sequences, we sought to identify mutations that might be consistent in these aaRS/tRNA pairs to leverage them as a starting point for further engineering. We identified A314Q in chPylRS^2PyA^ (corresponds to A302Q in *Mm*PylRS and A267Q in *Mb*PylRS) and A267Q in *Mb*PylRS^4ThzA^ as a shared mutation in the systems found for 2PyA and 4ThzA, respectively. We thus generated libraries of chPylRS^E7^ in which we fixed the mutation A314Q (Supplementary Table 6). We carried out selections with these libraries on 4OxzA and τMH as described above. Although no hit was found for τMH, we identified two aaRS/tRNA pairs that incorporate 4OxzA (chPylRS^4OxzA^ and chPylRS^4OxzA-2^). Interestingly, chPylRS^4OxzA-2^ carries the “QSW” mutations found in *Mb*PylRS^4ThzA^ as well as two additional mutations. This is surprising, as the transplantation of only the “QSW” mutations into *Mm*PylRS (*Mm*PylRS^4ThzA^) yields a poorly active enzyme (Supplementary Figure 8) and the “QSW” mutations cannot be readily transplanted to other PylRS systems (Supplementary Figure 9), as has been observed previously for other sets of mutations in PylRS^39^. In the pOS1T vector, chPylRS^4OxzA^ (Figure 3c) showed a superior ncAA-dependent sfGFP production in the presence of 4OxzA, compared to chPylRS^4OxzA-2^ (Supplementary Figure 10). Incorporation of 4OxzA with chPylRS^4OxzA^ was confirmed by LC-MS analysis of intact sfGFP150_TAG_ and LC-MS/MS of trypsinized sfGFP150_TAG_ (Supplementary Figures 2k and 3k, Supplementary Table 9). These results highlight the value of substrate profiling, particularly in light of the extremely weak starting point that could only be identified reproducibly above background based on this profiling.

**Figure 3.**
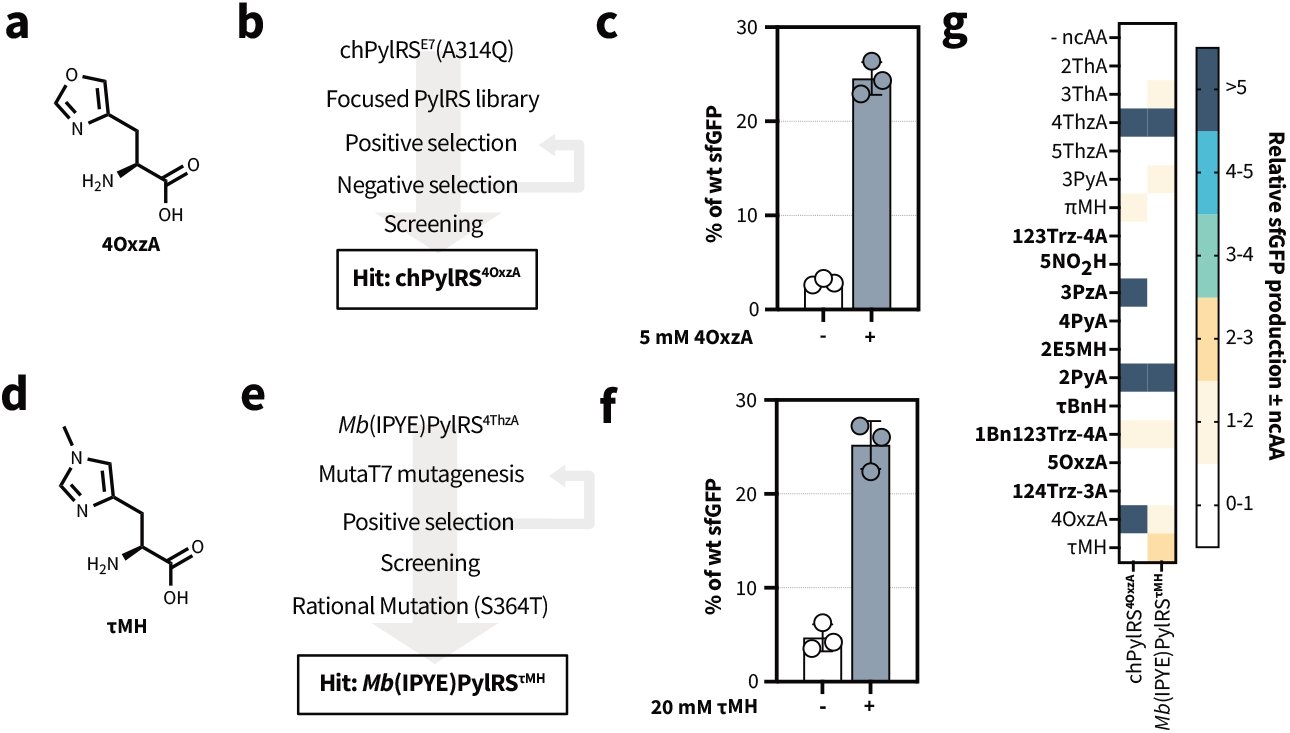
Evolution of additional aaRS/tRNA pairs for histidine-like ncAAs. a) Chemical structure of 4OxzA. b) Schematic of directed evolution workflow for identification of chPylRS^4OxzA^. c) Incorporation of 4OxzA into sfGP150_TAG_ using the chPylRS^4OxzA^ system. d) Chemical structure of τMH. e) Schematic of directed evolution workflow for identification of *Mb*(IPYE)PylRS^τMH^. f) Incorporation of τMH into sfGP150_TAG_ using the *Mb*(IPYE)PylRS^τMH^ system. g) The substrate specificity profiles of chPylRS^4OxzA^ and *Mb*(IPYE)PylRS^τMH^ showing the relative sfGFP production in the presence vs in the absence of the ncAA. The data were collected and analyzed as indicated in Figure 2a.

### *In vivo* mutagenesis succeeds where other strategies failed

From our initial ncAA targets, τMH was the ncAA that was most widely sought after, elusive, and frankly puzzling, not only to our lab but to others as well^41,22,27,42,21^. The lack of successful aaRS engineering suggested that the challenges might be related to steps ancillary to aminoacylation (such as cell uptake or metabolism) or post-aminoacylation (such as EF-Tu binding, ribosomal recognition, ribosomal incorporation, etc.)^21^. However, intracellular accumulation of τMH was supported by LC-MS analysis, suggesting that that τMH can accumulate in *E. coli* (data not shown). Additionally, although post-aminoacylation issues might be consistent with previous results^21^, such a constraint seemed unlikely given the relatively innocuous structure and canonical backbone. Thus, we evaluated our previous attempts to identify an aaRS/tRNA pair, which included classical screening using more than ten libraries from SSM and error-prone PCR, substrate walking from τBnH and *Mm*PylRS^τBnH^ with progressively smaller *N*^*τ*^ substituents, and the polyspecificity approach described for 4OxzA from *Mb*PylRS^4ThzA^. Based on recent success stories with *in vivo* evolution, we turned to a facile strategy for *in vivo* random mutagenesis, the MutaT7-transition^43^ system. Although MutaT7-transition has notable limitations compared to some newer (near-)continuous evolution systems, it was easier to establish, and the integration with our classical selection method was reliable. We selected *Mb*PylRS^4ThzA^ carrying the “IPYE” mutations^31^ as a parent sequence and performed ten rounds of passaging. These passages were coupled to positive selection in the presence of τMH, followed by selection on agar plates and screening in the presence and absence of the ncAA. A variant, termed *Mb*(IPYE)PylRS^p10^, was identified which showed an approximately 2.7-fold increase in sfGFP150^TAG^ suppression in the presence of τMH (Supplementary Figure 11). Most surprising from these engineering results was the location of the three mutations derived from MutaT7-transition, which were outside of the active site based on crystallography of homologous proteins^44,45^ and structure predictions^40,46^. Such mutations may be important for enzyme dynamics, modulating the orientation of other active-site residues, interaction with the tRNA, or directly interact with the ncAA through conformational changes induced during catalysis. From this variant, we confirmed incorporation of τMH into sfGFP150_TAG_ by LC-MS and LC-MS/MS, but phenylalanine misincorporation was significant as determined by an LC-MS/MS search (Supplementary Figure 2l and 3l). To reduce the observed misincorporation of phenylalanine, we introduced a rational mutation designed to reduce the size of the substrate binding pocket. This mutation significantly reduced the background incorporation of phenylalanine and increased the ncAA-dependent sfGFP150_TAG_ production to approximately 5.5-fold (Figure 3f). The final engineered aaRS (*Mb*(IPYE)PylRS^τMH^) enabled incorporation of τMH into sfGFP150_TAG_, as determined by LC-MS of intact sfGFP150_TAG_ and low misincorporation is suggested LC-MS/MS of trypsinized sfGFP150_TAG_ (Supplementary Figures 2m and 3m and Supplementary Table 9 and 10). This system represents a significant advance, providing the first genetic incorporation of this long elusive ncAA and highlights the difficulty in pinpointing the limiting factors when there are challenges encountered incorporating new ncAAs. We anticipate that this initial system will enable proof-of-concept studies for proteins with high expression levels, and the mutations identified here may serve as a foundation for future optimization efforts to enable incorporation in more challenging to express target proteins.

### Mutually orthogonal pairs enable dual incorporation of unique histidine-like ncAAs

Given the specificity profiles of several of the aaRSs, we sought to leverage the orthogonality between given aaRS-ncAA combinations to enable dual histidine-like ncAA encoding (Figure 4a). Several PylRS/tRNA^Pyl^ that are mutually orthogonal in their aminoacylation specificity have previously been used to incorporate two or more ncAAs into a single protein^24^. An established system is the combination of an **N**PylRS/*Ms*tRNA^Pyl^ (a PylRS with an N-terminal domain) and a **ΔN**PylRS/tRNA^ΔNPyl^ pair (a PylRS lacking an N-terminal domain)^24,27,47,48^. All aaRS pairs that we engineered in this study are **N**PylRS-type pairs, and we expected that we could combine them with a **ΔN**PylRS pair to incorporate two histidine-like ncAAs simultaneously. We selected the *Mm*PylRS/*Ms*tRNA^NPyl^_CUA_ and *G1*PylRS^MIFAF^/*Ma*tRNA^ΔNPyl^(8)_UCA_ as well as *Mm*PylRS/*Ms*tRNA^NPyl^_CUA_ and *Ma*PylRS^IFGFF^/*Ma*tRNA^ΔNPyl^ (8)_UCA_ for potential dual encoding of a subset of histidine-like ncAA combinations. We confirmed the opal suppression activity of the **ΔN**PylRS/tRNA^ΔNPyl^_UCA_ pairs (Supplementary Figure 12) and chose to proceed with *G1*PylRS^MIFAF^/*Ma*tRNA^ΔNPyl^_UCA_ based on its superior activity. Additionally, we confirmed the orthogonality relationship between the chPheRS^4PyA^/3C11-chPheT system towards other systems tested in our study (Supplementary Figure 14). We found that the chPheRS^4PyA^/3C11-chPheT and *Mm*PylRS/*Ms*tRNA^Pyl^ are not fully orthogonal, which is expected, as chPheRS^4PyA^ is a chimera of a PheRS and an **N**PylRS. Gratifyingly, the chPheRS^4PyA^/chPheT system is orthogonal towards the *G1*PylRS/*Ma*tRNA^ΔNPyl^ (8) pair.

**Figure 4.**
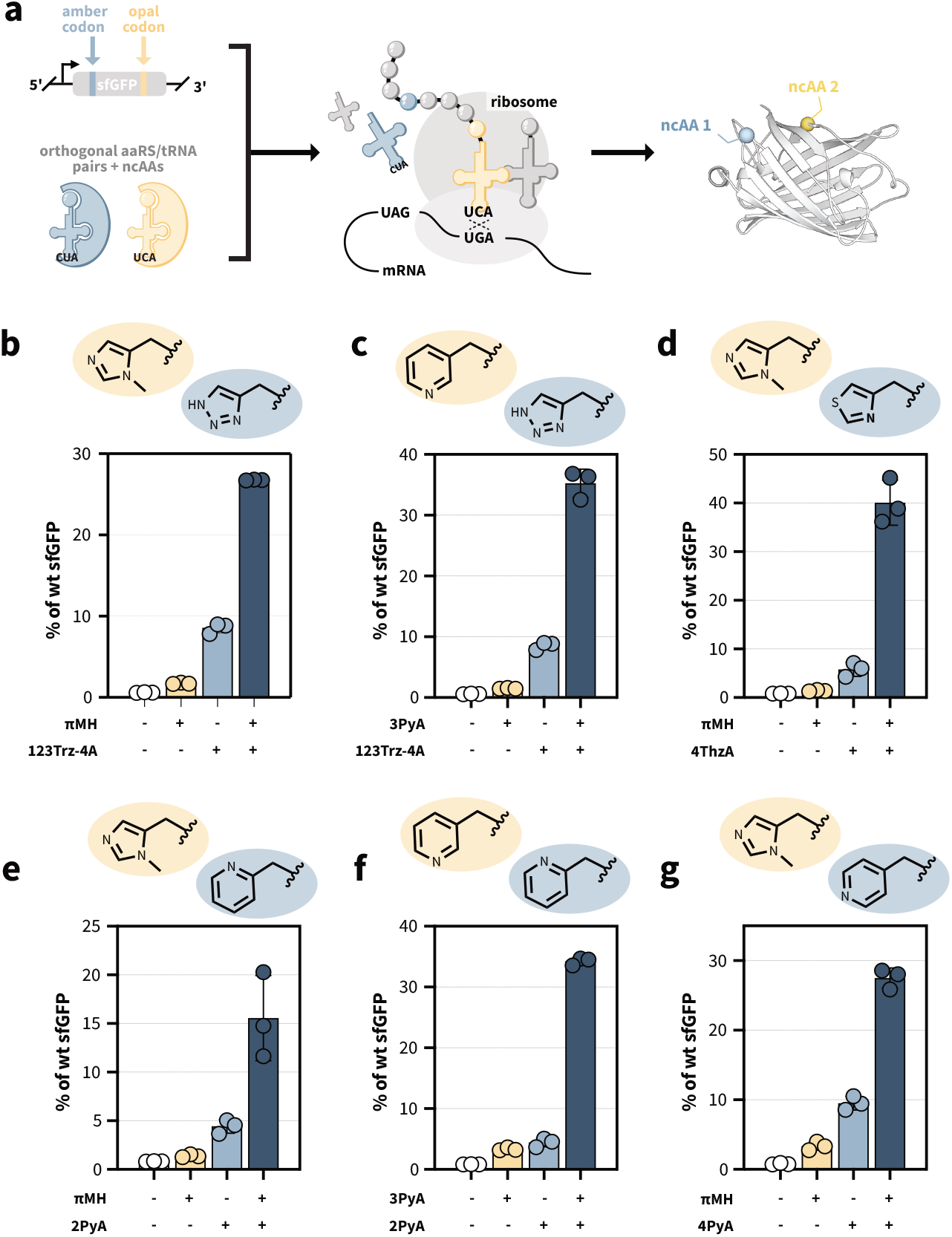
Dual histidine-like ncAA incorporation. a) Schematic representation of double suppression at amber and opal codons in sfGFP40_TAG_150_TGA_. b-g) Dual histidine-like ncAA incorporation into sfGFP40_TAG_150_TGA_. Side chains of histidine-like ncAAs incorporated at the amber codon (blue) and opal codon (yellow) are depicted. Data indicated as % of wt sfGFP ( as measured by fluorescence, excitation at 480 nm and emission at 510 nm, and normalized to the optical density at 600 nm) in the absence or presence of different ncAAs as percentage of a wt sfGFP reference. The mean and standard deviation of 3 biological replicates is shown.

Upon validating suitable pairs for dual incorporation, we defined approximately 20 possible dual combinations consisting of one of the **N**PylRS construct and *G1*PylRS^MIFAF^ from the aaRS-ncAA substrate profile. We explored dual suppression for a subset of six aaRS-ncAA combinations (Supplementary Table 11) at TAG and TGA codons in a sfGFP40_TAG_150_TGA_ reporter. For all of the six ncAA combinations (Figure 4b-g), we observed sfGFP production at efficiencies of 15-40% of wt sfGFP only in the presence of both ncAAs. We confirmed the presence of the desired ncAA at the given position using LC-MS/MS of trypsinized sfGFP40_TAG_150_TGA_ samples (Supplementary Figure 13a-l). When the ncAA for opal suppression was not supplied, ncAA-independent readthrough was more pronounced than when the ncAA for amber suppression, consistent with previous observations that opal suppression is less efficient for recoding than amber suppression. Two recently reported strategies could be envisioned to potentially circumvent this issue: quadruplet recoding^34^ or use of the Ochre recoded cell line^49^. However, because the incorporations were typically quantitative by LC-MS/MS, these strategies were not deemed necessary in this case.

Collectively, we encoded six combinations of histidine-like ncAAs using an **N**PylRS or chPheRS pair together with a **ΔN**PylRS pair and confirmed the orthogonality of the substrate selectivity of these aaRS-ncAA combinations. We anticipate that such combinations could facilitate the generation of highly tailored metal coordination sites in a manner that is so far only accessible through small molecule catalysts.

## Discussion

In this work, we have significantly expanded the diversity of histidine-like ncAAs accessible through genetic code expansion, developing nine novel aaRS/tRNA pairs for the site-specific incorporation of 12 histidine-like ncAAs with systematically varied properties. These ncAAs span a wide range of nitrogen p*K*_a_H values, including five alternative heterocycles beyond imidazole, and provide both *N*^π^- and *N*^τ^-substituted variants. This expanded toolkit addresses key gaps in the genetic code expansion landscape and provides researchers with significantly expanded chemical diversity for studying and engineering histidine-dependent reactivity.

The development of this toolkit required diverse engineering strategies. Classical directed evolution from *de novo* selections using SSM succeeded for several targets (123Trz-4A, 5NO_2_H, 3PzA, and 2E5MH), while others could be found through screening of existing variant libraries (τBnH), focused libraries with rationally fixed mutations (2PyA and 4OxzA), or *in vivo* mutagenesis (τMH). The particularly challenging case of τMH illustrates that structural similarity to successfully incorporated ncAAs does not guarantee ease of incorporation. Through extensive engineering, this work provides not only a toolkit but also highlights the versatility of engineering strategies that can—and sometimes must—be leveraged to access high-value incorporation targets.

Importantly, the systematic LC-MS/MS analysis revealed low-level misincorporation for some systems that was not detectable by intact mass analysis alone, which has also been previously reported^50^. While the misincorporation levels are low and unlikely to interfere with most applications, these findings underscore the importance of rigorous characterization. We recommend that researchers incorporating ncAAs for genetic code expansion applications conduct similar LC-MS/MS searches *with their target protein* to quantify potential misincorporation.

A striking finding from our substrate profiling is the remarkable orthogonality between many of the newly evolved aaRS variants. Unlike previously reported aaRS systems for phenylalanine, tyrosine, or lysine derivatives, which often show broad substrate promiscuity, the new variants for histidine-like ncAAs frequently discriminate between structurally similar substrates. We propose that this heightened specificity may be consequence of engineering aaRSs to accept small, polar substrates. In the case of aaRSs evolved for large ncAAs, less competition from canonical amino acids may allow for less stringent evolutionary solutions allowing greater polyspecificity. In contrast, these smaller, polar ncAAs may necessitate more customized substrate binding pockets with less tolerance for substrate variation. This hypothesis could explain why incorporation of diverse histidine-like ncAAs has lagged behind other ncAA classes and why we observe significant substrate orthogonality with these new systems. However, the surprising substrate profiles highlight blind spots in our understanding of aaRS activities, reinforce the continued importance of empirical screening, and underscore the unresolved need for medium/high-throughput computational methods that can robustly predict suitable variants for complex enzyme reactions.

While making individual ncAA incorporation more challenging, the high degree of orthogonality was advantageous for dual encoding. We anticipate that such combinations could enable the design of highly tailored metal coordination geometries and cooperative catalytic motifs that are currently inaccessible in natural proteins and difficult to achieve even with small molecule catalysts. The ability to install two distinct histidine-like ncAAs at defined positions provides experimental control over both the electronic properties and spatial arrangement of catalytic residues. Moreover, based on the discovery of multiple mutually orthogonal PylRS pairs^51^ and the ability to encode multiple distinct ncAAs simultaneously^47,48^, we expect that many more combinations of histidine-like ncAAs could be accessible.

Given the importance of histidine in enzyme systems, the toolkit presented here significantly expands the accessible chemical space for the study and engineering of proteins. The nine aaRS/tRNA pairs, 12 ncAAs, and validated dual incorporation systems provide researchers with powerful new methods for interrogating the roles of histidine residues in enzyme catalysis and for engineering proteins with novel or enhanced functions. Additionally, the starkly different substrate scopes of these aaRSs provide valuable tools for further mechanistic studies of aaRS activity. We anticipate that these tools will find broad application in mechanistic enzymology, enzyme engineering, genetic code expansion, and the design of artificial enzymes.

## Methods

### General instrumentation and materials

Absorbance and fluorescence measurements were measured on a Tecan Infinite M Nano+. ^1^H-NMR spectra were recorded in CDCl_3_, D_2_O, DMSO-d_6_ or MeOD on a Bruker AV-AV-400 (400 MHz), chemical shift d in ppm relative to solvent signals (d = 7.26 ppm for CDCl_3_, 4.79 ppm D_2_O, 2.50 for DMSO-d_6_, 3.31 for MeOD), coupling constants J are given in Hz. ^13^C-NMR spectra were recorded in D_2_O on a Bruker AV-AV-400 (400 MHz). All purchased chemicals were used without further purification. Automated flash column chromatography was performed on a Biotage Isolera One system using either Biotage Sfär Silica or Biotage Sfär C18 D columns. Preparative HPLC chromatography was performed on an Agilent 1260 Infinity II system using either a Gemini 5 μm NC-C18 110 Å or Zorbax NH2 7 μM column. All sequencing was conducted by Microsynth (Balgach, CH).

### Mass spectrometry methods

During aaRS/tRNA screening, initial high-throughput liquid chromatography-mass spectrometry (LC-MS) of sfGFP was conducted using an Agilent 1290 Infinity II LC system coupled to a single quadrupole mass spectrometer (ESI).

Verification of all proteins discussed in the text was additionally carried out by high-resolution LC-MS carried out at the Functional Genomics Center Zürich (FGCZ). Briefly, samples were resolved on an ACQUITY UPLC@BioResolve-RP-mAb (2.7 μm, 2.1 mm x 150 mm, 450 Å) column at a constant flow rate of 0.2 mL/min, with a column temperature of 60 °C. The LC gradient started at 95% buffer A (0.1% DFA) and 5% buffer B (25% acetonitrile/75% iso-propanol with 0.1% DFA). The proportion of buffer B was increased to 20% within 2 minutes, then ramped to 70% over 14 minutes, followed by a wash at 80% for 2 minutes. The gradient was then returned to 5% buffer B to re-equilibrate the column. The analysis was performed on a calibrated Waters Synapt G2-Si mass spectrometer directly coupled to the Waters H-Class UPLC. MS spectra were acquired in the positive-ion mode by scanning the m/z range from 400 to 5000 Da with a scan duration of 1 s and an interscan delay of 0.1 s. The spray voltage was set to 3 kV, the cone voltage to 50 V, and the source temperature to 100 °C. The data were recorded with the MassLynx 4.2 Software. The recorded m/z data of single peaks were deconvoluted into mass spectra by applying the maximum entropy algorithm MaxEnt1 (MassLynx 4.2) with a resolution of the output mass 0.5 Da/channel and Uniform Gaussian Damage Model at the half height of 0.7 Da.

To confirm the incorporation at the peptide level, LC-MS/MS was carried out by the FGCZ. Purified protein (approximately 1 mg/mL) was digested in solution by mixing 5 μL of sample with 40 μL digestion buffer (10 mM Tris, 2 mM CaCl_2_, pH 8.2). Protein was reduced and alkylated by 0.9 μL 100 mM Tris(2-carboxyethyl)phosphine + 1.4 μL 100mM chloroacetamide. 2 μL trypsin (100 ng/μL in 10 mM HCl) were added and microwave assisted digestion was carried out at 60 °C for 30 min. The samples were dried and dissolved in 20 μL ddH_2_O + 0.1% formic acid. Samples were analyzed on Waters M-class UPLC coupled to a calibrated Q-Exactive mass spectrometer (Thermo). Data-dependent (DDA) method was used in this analysis. Samples were loaded onto a nanoEase M/Z Symmetry C18 trap column (180 μm × 20 mm, 100 Å, 5 μm particle size) and separated on a nanoEase M/Z HSS C18 T3 column (75 μm × 250 mm, 100 Å, 1.8 μm particle size), at a constant flow rate of 300 nL/min, with a column temperature of 50 °C. The LC gradient started at 5% solvent B (100% acetonitrile with 0.1% formic acid) and was increased to 35% over 42 minutes, then ramped to 60% within 5 minutes, followed by a wash at 95% for 10 minutes. The gradient was then returned to 5% buffer B to re-equilibrate the column. For MS setting, one scan cycle comprised of a full scan MS survey spectrum, followed by HCD (higher-energy collision dissociation) fragmentation on the 12 most intense signals for cycle. Full-scan MS spectra (350–1’500 m/z) were acquired at a resolution of 70’000 at 400 m/z, while HCD MS/MS spectra were recorded in the FT-Orbitrap at a resolution of 35’000. HCD MS/MS spectra were performed with a target value of 1e5 using a normalized collision energy 25%. The samples were acquired using internal lock mass calibration on m/z 371.1010 and 445.1200. The MS raw files were searched against sfGFP sequence sequences by Byonic 5.2 (Proteinmetrics, USA) with the consideration of carbamidomethylation at cysteine residues and oxidation at methionine residues. In addition, the mutated site was set to J (default mass is 100 Daltons) in sfGFP sequence. The mass increase was set to different ncAA and normal amino acids as variable modifications. For example, +47.0789 at J indicated Phenylalanine. If misincorporation could be identified with the manual inspection of MS spectra, the extracted ion chromatograms (XIC) for each observed amino acid were integrated to provide a rough approximation of the misincorporation. We highlight two caveats that make this analysis an approximation not a direct quantitation: i) ionization efficiencies between point mutations of peptides can vary and ii) in some cases mutations can change the efficiency of trypsin digestion. Thus, we provide relative integrations only as an extremely coarse estimate.

Small-molecule LC-MS analysis and UPLC analysis were both conducted with an Agilent 1290 Infinity II LC system coupled to a single quadrupole mass spectrometer (ESI).

### p*K*_a_ calculations

For the p*K*_a_ estimations (Figure 1a) literature reported p*K*_a_s were only available for seven of the 17 structures. Three p*K*_a_ estimators were evaluated—Qupkake^52^, MolGpka^53^, and Chemicalize from ChemAxon (https://chemicalize.com/)—using the neutral amino acid, the zwitterionic form of the amino acid, and the side chain. Each single method and voting combinations of multiple methods were evaluated to determine the tool most accurate for recapitulating literature values. The most accurate method consisted of an averaging of the values from Qupkake for the zwitterionic form, Qupkake for the side chain, and Chemicalize.

### aaRS SSM Library construction

Libraries were created using modified Golden Gate cloning. A pSL plasmid encoding the corresponding synthetase was amplified using inverse PCR with primers carrying a BsaI restriction site and an additional 6 bp overhang at the 5’ end (Supplementary Tables 11-13) using Q5 Hot Start High-Fidelity DNA polymerase. PCR products were purified using an NEB Monarch PCR cleanup kit. The purified PCR products were digested with DpnI and BsaI in 1x rCutSmart. Digests were carried out at 37 °C overnight. Digested products were purified using an NEB Monarch PCR cleanup kit. DNA ligation reactions contained T4-ligase and 1x T4-ligase-buffer. Ligation was carried out at 16 °C overnight. Ligation products were purified using an NEB Monarch PCR cleanup kit. Electrocompetent NEB10β *E. coli* (100 μL) were transformed with ligation product by electroporation in a cuvette with 2 mm gap on an Eppendorf Eporator with 250 Ω and 2500 V with pulse times of approximately 5 ms. The cells were recovered in SOC media for 1 h at 37 °C with shaking at 220 rpm. The number of transformants was estimated by dilution series plating of LB-agar with 50 μg/mL kanamycin. The recovered cells were transferred to 10 mL LB media with 50 μg/mL kanamycin and grown for 5 h. The cells were harvested by centrifugation, and the DNA was isolated using an NEB Monarch Plasmid Miniprep Kit. The library quality was confirmed by Sanger sequencing of 2-3 individual clones.

### aaRS error-prone PCR Library construction

Libraries were created using modified Golden Gate cloning. The backbone of a pSL plasmid encoding the corresponding synthetase was amplified using PCR with primers carrying a BsaI restriction site and an additional 6 bp overhang at the 5’ end (Supplementary Table 15) using Q5 Hot Start High-Fidelity DNA polymerase. The synthetase region to be mutagenized was amplified using PCR with primers carrying a BsaI restriction site and an additional 6 bp overhang at the 5’ end using the JBS Error-Prone Kit (Jena Biosciences) according to the manufacturers protocol. PCR products were purified using an NEB Monarch PCR cleanup kit. The purified PCR products were pooled and digested with DpnI and BsaI in 1x rCutSmart. Digests were carried out at 37 °C overnight. Digested products were purified using an NEB Monarch PCR cleanup kit. DNA ligation reactions contained T4-ligase and 1x T4-ligase-buffer. Ligation was carried out at 16 °C overnight. Ligation products were purified using an NEB Monarch PCR cleanup kit. Electrocompetent NEB10β *E. coli* (100 μL) were transformed with ligation product by electroporation in a cuvette with 2 mm gap on an Eppendorf Eporator with 250 Ω and 2500 V with pulse times of approximately 5 ms. The cells were recovered in SOC media for 1 h at 37 °C with shaking at 220 rpm. The number of transformants was estimated by dilution series plating of LB-agar with 50 μg/mL kanamycin. The recovered cells were transferred to 10 mL LB media with 50 μg/mL kanamycin and grown for 5 h. The cells were harvested by centrifugation, and the DNA was isolated using an NEB Monarch Plasmid Miniprep Kit.

### aaRS Library selections

For positive selections, electrocompetent NEB10β cells (100 μL) carrying the corresponding pDPS2 plasmid were transformed with 250 ng of a given library by electroporation in a cuvette with 2 mm gap on an Eppendorf Eporator at 250 Ω and 2500 V with pulse times of approximately 5 ms. The cells were recovered in SOC media for 1 h at 37 °C with shaking and transferred to 10 mL LB media with 50 μg/mL kanamycin, 10 μg/mL tetracycline, and ncAA at the given concentration. The cells were grown for 1-2 h, harvested by centrifugation, resuspended in 250 μL LB media and plated on LB-agar with 0.4% arabinose, 50 μg/mL kanamycin, 10 μg/mL tetracycline, 100 μg/mL chloramphenicol, 0.4% arabinose, and ncAA at the given concentration (Supplementary Table 1). Plates were incubated for 24 h at 37 °C. If additional selection rounds were carried out, cells were collected by washing the plate with 10 mL LB media and harvesting the cells by centrifugation. Plasmid DNA was isolated using an NEB Monarch Plasmid Miniprep Kit. DNA was digested with AgeI in 1x rCutsmart buffer for 18 h at 37 °C to remove the selection plasmid and purified using an NEB Monarch PCR cleanup kit. If additional selection rounds were not carried out, we proceeded as though the step was the final positive selection round described further below.

For negative selections, electrocompetent NEB10β cells (100 μL) carrying the corresponding pBARN plasmid were transformed with 50 ng of library DNA by electroporation in a cuvette with 2 mm gap on an Eppendorf Eporator at 250 Ω and 2500 V with pulse times of approximately 5 ms. The cells were recovered in SOC media for 1 h at 37 °C with shaking and transferred to 10 mL LB media with 50 μg/mL kanamycin and 35 μg/mL chloramphenicol. The cells were grown in the presence of the antibiotics for 1-2 h. The cells were harvested by centrifugation, resuspended in 250 μL LB media and plated on LB-agar with 0.4% arabinose, 50 μg/mL kanamycin, and 35 μg/mL chloramphenicol. Plates were incubated for 24 h at 37 °C. Cells were collected by washing the plate with 10 mL LB media; cells were harvested by centrifugation; and plasmid DNA was purified using an NEB Monarch Plasmid Miniprep Kit. DNA was digested with AgeI in 1x rCutsmart buffer. Digests were carried out at 37 °C overnight to the remove selection plasmid and purified using an NEB Monarch PCR cleanup kit.

For screening after the final round of positive selection, individual colonies were used to inoculate 96-well plates with 250 μL 2xYT media with 50 μg/mL kanamycin, 10 μg/mL tetracycline. Cultures were grown for 24 h at 37 °C with shaking at 400 rpm. Subsequently, 25 μL of each culture was used to inoculate 96-well plates with 250 μL 2xYT media with 50 μg/mL kanamycin, 10 μg/mL tetracycline, 0.4% arabinose, and ncAA at the given concentration. Cultures were grown for 24 h at 37 °C with shaking at 400 rpm. 100 μL of each culture were transferred to a 96-well clear well plate. Fluorescence (excitation at 480 nm and emission at 510 nm) and absorbance at 600 nm were measured. DNA from cultures with high Fluorescence/OD_600_ ratios was isolated using an NEB Monarch Plasmid Miniprep Kit and analyzed by Sanger sequencing to identify mutations in the PylRS gene.

### Evolution of *Mb*(IPYE)PylRS^4ThzA^ for τMH incorporation

NEB10β *E. coli* were co-transformed with pSLdT7-*Mb*(IPYE)PylRS^4ThzA^, pDPS2-*Ms*tRNA^Pyl^_CUA_ and pDae079so by heat-shock, recovered in SOC for 1 h at 37 °C and plated on LB-agar with 50 μg/mL kanamycin and 10 μg/mL tetracycline and 100 μg/mL spectinomycin. An individual colony was inoculated into LB-media with 50 μg/mL kanamycin and 10 μg/mL tetracycline and 100 μg/mL spectinomycin and grown overnight. Subsequently, cells were diluted 1:100 into LB-media with 50 μg/mL kanamycin and 10 μg/mL tetracycline and 100 μg/mL spectinomycin, 0.2% arabinose, 5 mM τMH and grown for 8h. Cells were diluted and grown four additional times with addition of 35-100 μg chloramphenicol. Finally, cells were harvested by centrifugation for 10 min at 4 °C and 4200 g. Plasmid DNA was isolated using an NEB Monarch Plasmid Miniprep Kit and digested with AgeI in 1x rCutsmart buffer for 18 h at 37 °C to remove the selection and mutagenesis plasmids and purified using an NEB Monarch PCR cleanup kit. A positive selection round was carried out as described in the section “aaRS library selection”.

Lastly, NEB10β *E. coli* were co-transformed with the pSLdT7-*Mb*(IPYE)PylRS^p10^ and pDPS2-*Ms*tRNA^Pyl^_CUA_ by heat shock (42 °C, 30 s), recovered in SOC for 1 h at 37 °C and plated on LB-agar with 50 μg/mL kanamycin and 10 μg/mL tetracycline. Individual colonies were used to inoculate 5 mL autoinduction media with 100 μg/mL kanamycin, 20 μg/mL tetracycline, 0.4% arabinose and 20 mM τMH. The cultures were grown for 24 h at 37 °C with shaking at 240 rpm. The cells were harvested by centrifugation for 10 min at 4 °C and 4200 g; the supernatant was decanted; and the cell pellets were stored at -80 °C. To isolate the sfGFP, the cells were thawed at room temperature and resuspended in lysis buffer (500 μL, 20 mM Tris, 300 mM NaCl, pH 7.2 at 4 °C, 0.2% *n*-octyl *beta*-D-thioglucopyranoside, 4 mg/mL Lysozyme). The lysis was conducted at 22 °C for 4 h. The sfGFP was isolated by purification with Ni-NTA resin (HisPur from Thermo Scientific) according to the manufacturer’s instructions. The purified sfGFP was analyzed by LC-MS and LC-MS/MS as indicated in the Mass Spectrometry section.

### sfGFP amber suppression assay for aaRS/tRNA orthogonality screening

NEB10β *E. coli* were co-transformed with pDPS2 and corresponding pSL plasmid by heat shock (42 °C, 30 s), recovered in SOC for 1 h at 37 °C and plated on LB-agar with 50 μg/mL kanamycin and 10 μg/mL tetracycline. Plates were incubated for 24 h at 37 °C. Individual colonies were used to inoculate 250 μL autoinduction media with 100 μg/mL kanamycin, 20 μg/mL tetracycline, and ncAA at the indicated concentrations. Cultures were grown for 24 h at 37 °C with shaking at 400 rpm. wt sfGFP was analogously expressed from pBAD-sfGFP by omission of 20 μg/mL tetracycline addition for the culturing of cells. 100 μL of each culture was transferred to a 96-well clear well plate. Fluorescence (excitation at 480 nm and emission at 510 nm) and absorbance at 600 nm were measured. Data is indicated as normalized fluorescence excitation at 480 nm and emission at 510 nm normalized to the optical density at 600 nm).

### sfGFP amber suppression assay

NEB10β *E. coli* were co-transformed with pBAD-sfGFP150_TAG_ (modified from Addgene #85483, a gift from Ryan Mehl’s lab)^54^ and the corresponding aaRS/tRNA expression plasmid by heat shock (42 °C, 30 s), recovered in SOC for 1 h at 37 °C and plated on LB-agar with 50 μg/mL kanamycin and 10 μg/mL tetracycline. Plates were incubated for 24 h at 37 °C. Individual colonies were used to inoculate 250 μL LB media with 50 μg/mL kanamycin, 10 μg/mL tetracycline. Cultures were grown for 24 h at 37 °C with shaking at 400 rpm. Subsequently, cultures were diluted into 250 μL autoinduction media with 100 μg/mL kanamycin, 20 μg/mL tetracycline, 0.4% arabinose and ncAA at the indicated concentrations. wt sfGFP was analogously expressed from pBAD-sfGFP by omission of tetracycline addition for the culturing of cells. 100 μL of each culture were transferred to a 96-well clear well plate. Fluorescence (excitation at 480 nm and emission at 510 nm) and absorbance at 600 nm were measured. Data indicated as % of wt sfGFP shows normalized fluorescence (excitation at 480 nm and emission at 510 nm normalized to the optical density at 600 nm) in the absence or presence of different ncAAs as percentage of a wt sfGFP reference. Data indicated as relative sfGFP production ± ncAA are calculated from the normalized fluorescence in the presence of the ncAA divided by the normalized fluorescence in the absence of the ncAA.

### sfGFP dual suppression assay

NEB10β *E. coli* were co-transformed with pBAD-sfGFP40_TAG_150_TGA_ (modified from Addgene #85483)^54^ and the corresponding aaRS/tRNA expression plasmids by heat shock (42 °C, 30 s), recovered in SOC for 1 h at 37 °C, and plated on LB-agar with 50 μg/mL kanamycin and 10 μg/mL tetracycline and 35 μg/mL chloramphenicol. Plates were incubated for 24 h at 37 °C. Individual colonies were used to inoculate 250 μL LB media with 50 μg/mL kanamycin, 10 μg/mL tetracycline, 35 μg/mL chloramphenicol. Cultures were grown for 24 h at 37 °C with shaking at 400 rpm. Subsequently, cultures were diluted into 250 μL autoinduction media with 100 μg/mL kanamycin, 20 μg/mL tetracycline, 70 μg/mL chloramphenicol, 0.4% arabinose and ncAA at the indicated concentrations. Cultures were grown for 24 h at 37 °C with shaking at 400 rpm. wt sfGFP was analogously expressed from pBAD-sfGFP by omission of tetracycline and chloramphenicol addition for the culturing of cells. 100 μL of each culture were transferred to a 96-well clear well plate. Fluorescence (excitation at 480 nm and emission at 510 nm) and absorbance at 600 nm were measured. Data indicated as % of wt sfGFP show normalized fluorescence (excitation at 480 nm and emission at 510 nm normalized to the optical density at 600 nm) in the absence or presence of different ncAAs as percentage of a wt sfGFP reference.

### Analysis of ncAA incorporation in sfGFP150_TAG_

NEB10β *E. coli* were co-transformed with pBAD-sfGFP150_TAG_ and the corresponding aaRS/tRNA expression plasmid by heat shock (42 °C, 30 s), recovered in SOC for 1 h at 37 °C and plated on LB-agar with 50 μg/mL kanamycin and 10 μg/mL tetracycline. Plates were incubated for 24 h at 37 °C. Individual colonies were used to inoculate 5 mL autoinduction media with 100 μg/mL kanamycin, 20 μg/mL tetracycline, 0.4% arabinose and ncAA at the indicated concentration. The cultures were grown for 24 h at 37 °C with shaking at 240 rpm. The cells were harvested by centrifugation for 10 min at 4 °C and 4200 g; the supernatant was decanted; and the cell pellets were stored at -80 °C. To isolate the sfGFP, the cells were thawed at room temperature and resuspended in lysis buffer (500 μL, 20 mM Tris, 300 mM NaCl, pH 7.2 at 4 °C, 0.2% *n*-octyl *beta*-D-thioglucopyranoside, 4 mg/mL Lysozyme). The lysis was conducted at 22 °C for 4 h. The sfGFP was isolated by purification with Ni-NTA resin (HisPur from Thermo Scientific) according to the manufacturer’s instructions. The purified sfGFP was analyzed by LC-MS and LC-MS/MS as indicated in the Mass Spectrometry section.

### Analysis of dual ncAA incorporation in sfGFP

NEB10β *E. coli* were co-transformed with pBAD-sfGFP40_TAG_150_TGA_ (modified from Addgene #85483)^54^ and the corresponding aaRS/tRNA expression plasmids by heat shock (42 °C, 30 s), recovered in SOC for 1 h at 37 °C, and plated on LB-agar with 50 μg/mL kanamycin and 10 μg/mL tetracycline and 35 μg/mL chloramphenicol. Plates were incubated for 24 h at 37 °C. Individual colonies were used to inoculate 5-8 mL autoinduction media with 100 μg/mL kanamycin, 20 μg/mL tetracycline, 70 μg/mL chloramphenicol, 0.4% arabinose and ncAAs at the indicated concentrations. The cultures were grown for 24 h at 37 °C with shaking at 240 rpm. The cells were harvested by centrifugation for 10 min at 4 °C and 4200 g; the supernatant was decanted; and the cell pellets were stored at -80 °C. To isolate the sfGFP, the cells were thawed at room temperature and resuspended in lysis buffer (500 μL, 20 mM Tris, 300 mM NaCl, pH 7.2 at 4 °C, 0.2% *n*-octyl *beta*-D-thioglucopyranoside, 4 mg/mL Lysozyme). The lysis was conducted at 22 °C for 4 h. The sfGFP was isolated by purification with Ni-NTA resin (HisPur from Thermo Scientific) according to the manufacturer’s instructions. The purified sfGFP was analyzed by LC-MS and LC-MS/MS as indicated in the Mass Spectrometry section.

### Intracellular τMH concentration determination

A glycerol stock of NEB10β *E. coli* was used to inoculate 5 mL 2xYT media. Cultures were grown for 16 h at 37 ºC. Cultures were diluted 1:100 into 5 mL 2xYT media with or without 5 mM τMH. Cultures were grown for 16h at 37ºC. The OD_600_ of each culture was measured. The cells were harvested by centrifugation for 5 min at 4 °C and 4200 g; the supernatant was decanted; and the cells were washed four times with phosphate buffered saline (pH 7.4). The cells were harvested by centrifugation for 5 min at 4 °C and 4200 g and resuspended in 400 μL H_2_O/MeOH (2:3). Approximately 300 mg of 0.1 mm Glass Beads (Scientific Industries, Inc) were added, and cells were vortexed for 5 minutes. Lysates were clarified by centrifugation for 30 min at 4 ºC and 20000 g. Supernatants were filtered through a Millipore Amicon Ultra Centrifugal Filter with 30kDa MW cutoff by centrifugation for 15 min at 4ºC and 14000 g. Lysates were analyzed on an ACQUITY UPLC® BEH C18 1.7 μm column using an Agilent 1290 Infinity II LC system coupled to a single quadrupole mass spectrometer (ESI) in single-ion-monitoring mode (m/z = 170). The LC gradient started at with a flow of 0.4 mL/min and 100% buffer A (H_2_O) for 1 min, followed by a gradient to 95% buffer B (MeCN) for 2 min, followed by a gradient to 100% buffer A for 30 s, followed by 100% buffer A for 1 min.

### ncAA-aaRS affinity prediction with Boltz2

Structure and affinity predictions were initially conducted with Boltz2^46^ were carried out using a Google Colab implementation^55^. It was observed that the stereochemistry of the ncAA (but not ATP) was sometimes inverted during the prediction. Thus, the Boltz2 predicted protein structures with ATP bound were used to perform affinity predictions with Gnina, a fork of autodock vina^56^. We also docked the AA-AMP adducts to the predicted protein structure in the absence of ATP, but we observed a similar lack of correlation between predicted binding affinities and incorporation efficiencies. LogP values were calculated in python using the RDKit (https://www.rdkit.org).

### Notes

Note 1, Nomenclature: Herein, we use the IUPAC nitrogen labels. For completeness, *N*^τ^ (or τ-N) is also sometimes referred to as ε-N, 1-N, or 3-N, and *N*^π^ (or π-N) is also sometimes referred to as δ-N, 1-N, or 3-N.

Note 2, chPylRS^E7^ evolution: We reasoned that a more efficient PylRS construct might beneficial for the incorporation of “difficult-to-incorporate” ncAAs^57^. We focused on the optimization of a chimeric PylRS (chPylRS) construct. A small selection of chPylRS constructs were prepared by fusing the N-terminal domain of PylRS from thermophilic archaea and bacteria^58^ to the C-terminal domain of PylRS from *Methanosarcina mazei* (83-454), a mesophilic archaeon (Supplementary Table 3). The constructs were expressed under control of a glnS promoter (plasmid: pSL) and tested for the incorporation of the near-native substrate *N*^ε^-(tert-butoxycarbonyl)-L-lysine (BocK) at low concentrations (0.2 mM). Among the tested constructs, chPylRS^*Me-Mm*^ was the only active construct but gratifyingly displayed slightly increased suppression efficiencies with *Mm*tRNA^Pyl^_CUA_ compared to *Mm*PylRS and chPylRS^mb(IPYE)-mm^, a previously reported chPylRS system^59^. chPylRS^*Me-Mm*^ consists of the N-terminal domain of the PylRS from *Methanohalobium evestigatum* (1-94) and the C-terminal domain of PylRS from *Methanosarcina mazei* (83-454). To further improve the activity of chPylRS^*Me-Mm*^, we pursued a strategy analogous to a previous PylRS engineering campaign^60^. In short, residues 1-94 of chPylRS^*Me-Mm*^ were subjected to high error rate error-prone PCR (3-7 mutations/kb), positive selection and screening of fluorescence intensity at low BocK concentrations (0.2 mM). The best performing clones were pooled and subjected to a second round of error-prone PCR, positive selection, and screening of fluorescence intensity at low BocK concentrations (0.2 mM). After two rounds, a clone carrying the mutations S18C/K45E/P68Q/K70I/V74A/N80S/K93I (chPylRS^E7^) emerged. From our selection plasmid, chPylRS^E7^ displayed approximately 4.3-fold improved suppression efficiencies at low BocK concentrations (0.2 mM) compared to chPylRS^*Me-Mm*^ and approximately 6.9-fold improved suppression efficiencies compared to *Mm*PylRS (Supplementary Figure 1). Notably, upon transference of chPylRS^E7^ to our optimized plasmid for robust production of aaRS/tRNA pairs (pOS1T), the advantage over *Mm*PylRS was lost, suggesting that these mutations are beneficial for expression from the selection/screening plasmid (pSL) but not in the final GCE plasmid (pOS1T). Interestingly, in our hands, this outcome also appears to be the case for several other reported engineered aaRS/tRNA pairs, highlighting the role of the plasmid—likely through attenuation of protein or tRNA production—in observed engineering outcomes. Nonetheless, the improvements in the selection plasmid were significant, and thus we hypothesized that this variant might be better suited for selection and screening with new aaRSs.

## Acknowledgments

For funding, the authors thank the University of Zurich, the Swiss National Science Foundation (grant no. 10002490), and the UZH CanDoc Award (ANP: grant no. FK-24-093, SF: grant no. FK-22-086). The authors thank the Functional Genomics Center Zurich (FGCZ) for assistance with high-resolution LC-MS and LC-MS/MS data collection and analysis.

## Author contributions

ADL conceived of and supervised the project. ADL, ANP, and SF contributed to design of experiments. ARL carried out chPylRS^E7^ evolution under the supervision of ANP. ANP, SF, and BPM synthesized ncAAs. ANP and SF engineered and validated the aaRS/tRNA pairs for incorporation of ncAAs into sfGFP. All authors contributed to writing and editing the final paper.

