## Supplementary_information for "Genetic incorporation of diverse non-canonical amino acids for histidine substitution"

[†]These authors contributed equally.

Table of Contents


| | | |
|---|---|---|
| I. | Supplementary data figures | 2 |
| II. | Synthesis | 16 |
| III. | Supplementary tables | 54 |
| IV. | DNA and Protein Sequences | 59 |
| V. | Plasmid Construction | 68 |

I. **Supplementary data figures**

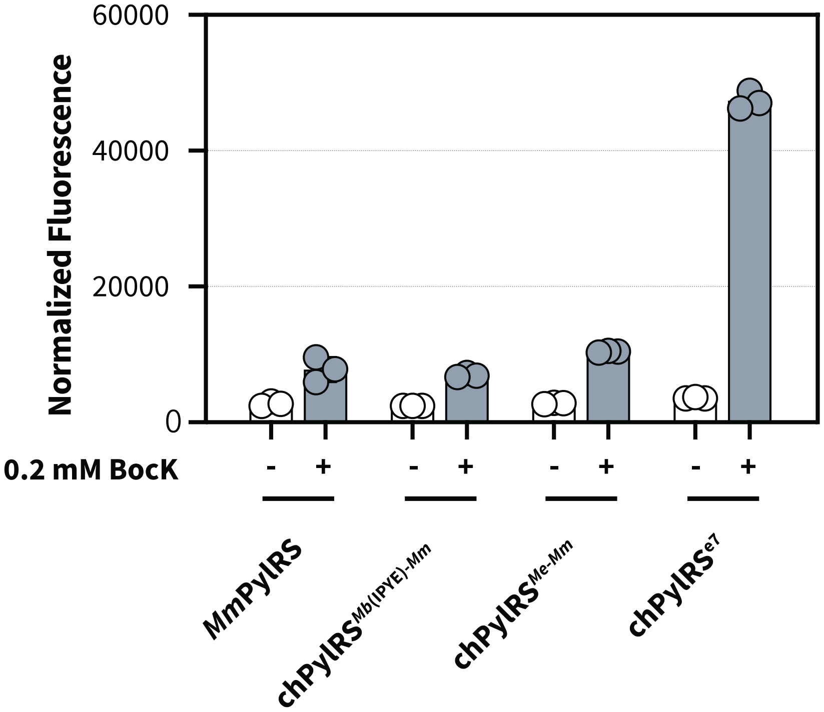


**Supplementary Figure 1 | Validation of chPylRS$^{E7}$ variants for incorporation of BocK.** Suppression of sfGFP150$_{TAG}$ in NEB10β in the presence or absence of different PylRS variants with 0.2 mM BocK. Data show normalized fluorescence (excitation at 480 nm and emission at 510 nm normalized to the optical density at 600 nm) and represents the mean and standard deviation of 3 biological replicates.

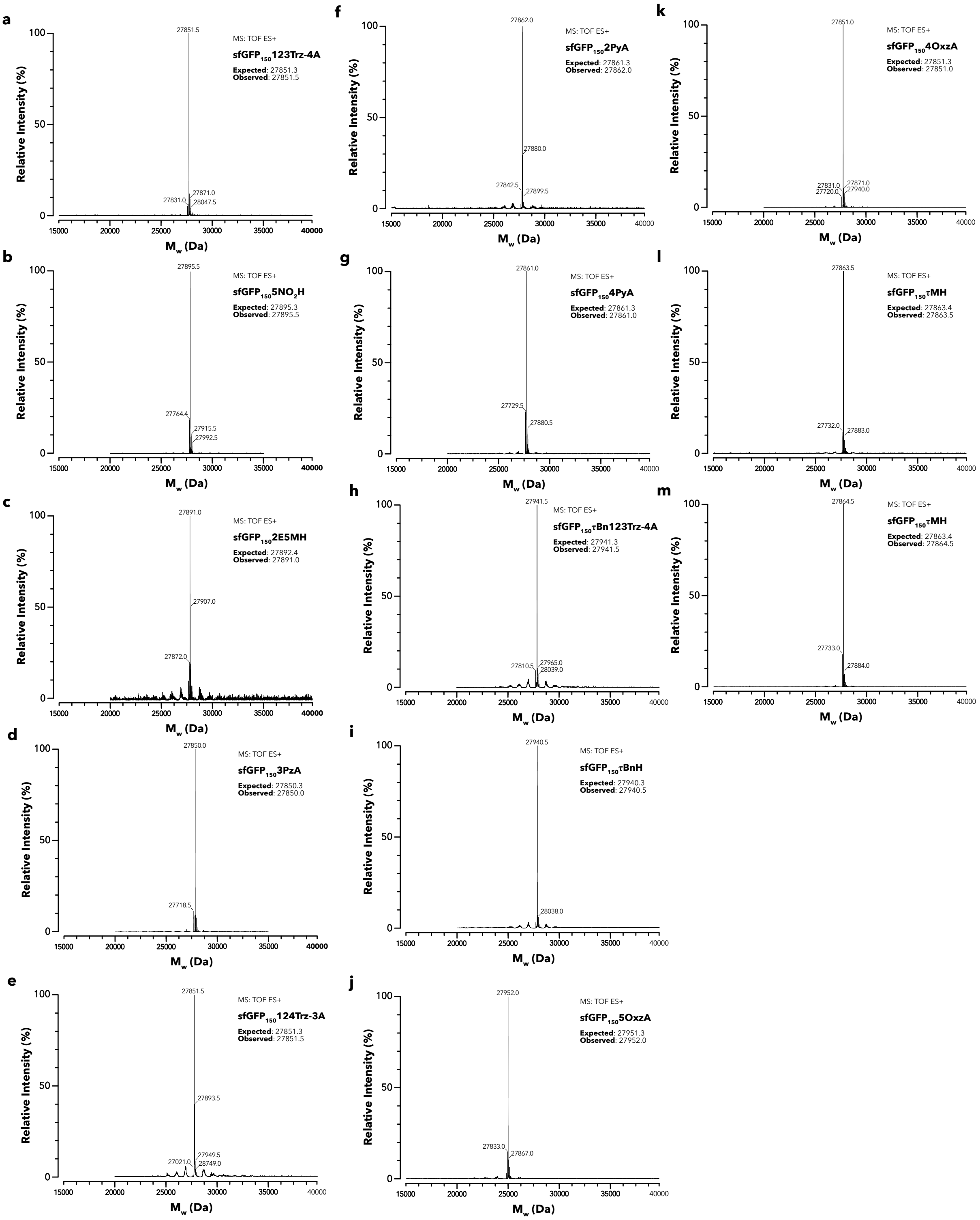


**Supplementary Figure 2 | LC-MS spectra of sfGfP150$_{TAG}$ containing a single histidine-like ncAA.** Intact LC-MS analysis (positive electrospray time of flight) of sfGFP150$_{TAG}$ production in the presence of different ncAAs (**a-m**). The expected and observed masses confirming incorporation of the desired ncAA are indicated. Additional peaks, consistent with typically observed [$H_2O$] elimination (-18 Da), $[Na]^+$ addition (+23 Da) or loss of N-terminal [Met] (-131 Da) were observed.

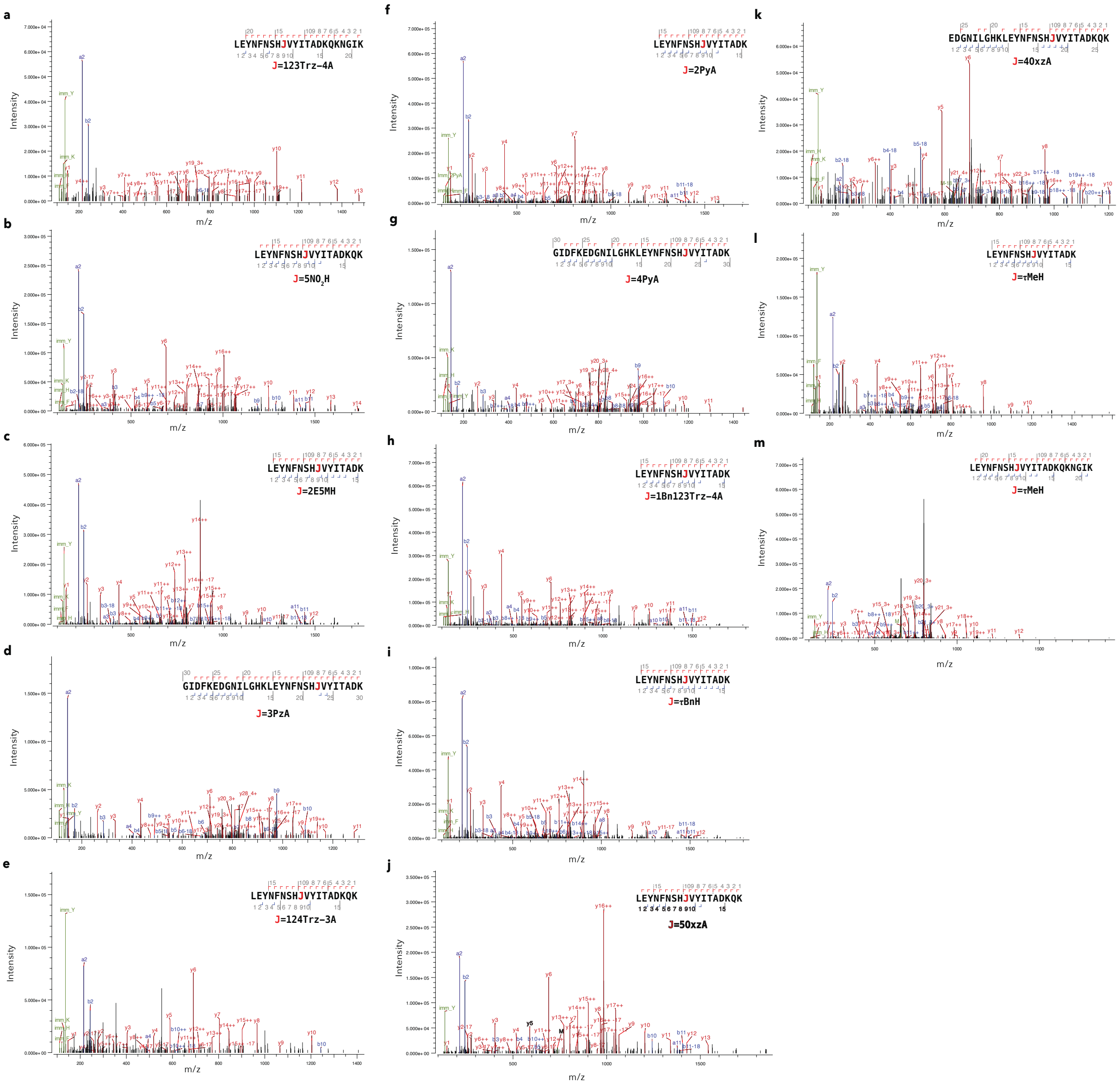


**Supplementary Figure 3 | LC-MS/MS spectra of sfGFP150$_{TAG}$ containing a single histidine-like ncAA.** A representative LC-MS/MS spectrum from a tryptic digest of sfGFP150$_{TAG}$ expressed with different ncAAs and their corresponding aaRS/tRNA pairs. Typically, multiple peptides containing the desired ncAA were observed and no peptides for canonical amino acid incorporation were observed. For samples of sfGFP-5$NO_2$H, sfGFP-τBnH, sfGFP-τBn123Trz-4A and sfGFP-124Trz-1A, some peptides containing canonical amino acids were observed (see **Supplementary Table 9 and 10**).

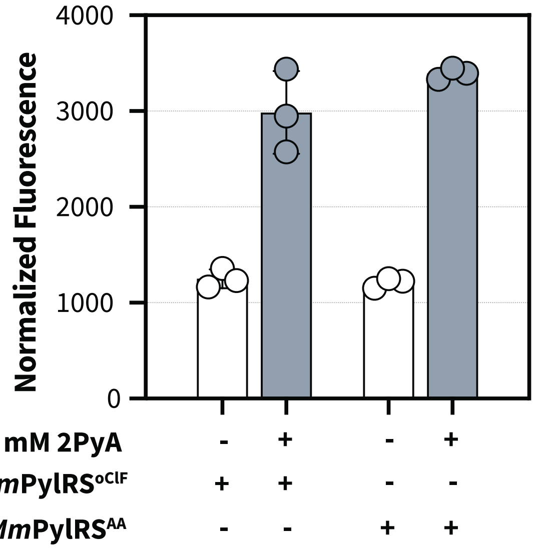


**Supplementary Figure 4 | Testing of reported variants for incorporation of 2PyA.** Suppression of sfGFP150$_{TAG}$ in NEB10β in the presence or absence of with 6 mM 2PyA with different PylRS variants (*Mm*PylRS$^{oClF}$ [36] and *Mm*PylRS$^{AA}$ [61]). Data show normalized fluorescence (excitation at 480 nm and emission at 510 nm normalized to the optical density at 600 nm) and represents the mean and standard deviation of 3 biological replicates.

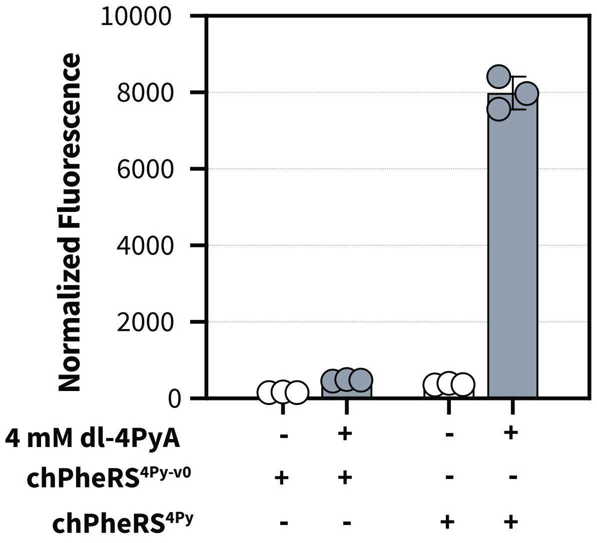


**Supplementary Figure 5 | Validation of chPheRS$^{4PyA}$ variants for incorporation of 4PyA.** Suppression of sfGFP150$_{TAG}$ in NEB10β in the presence or absence of 4 mM 4PyA with different **chPheRS** variants. Data show normalized fluorescence (excitation at 480 nm and emission at 510 nm normalized to the optical density at 600 nm) and represents the mean and standard deviation of 3 biological replicates.

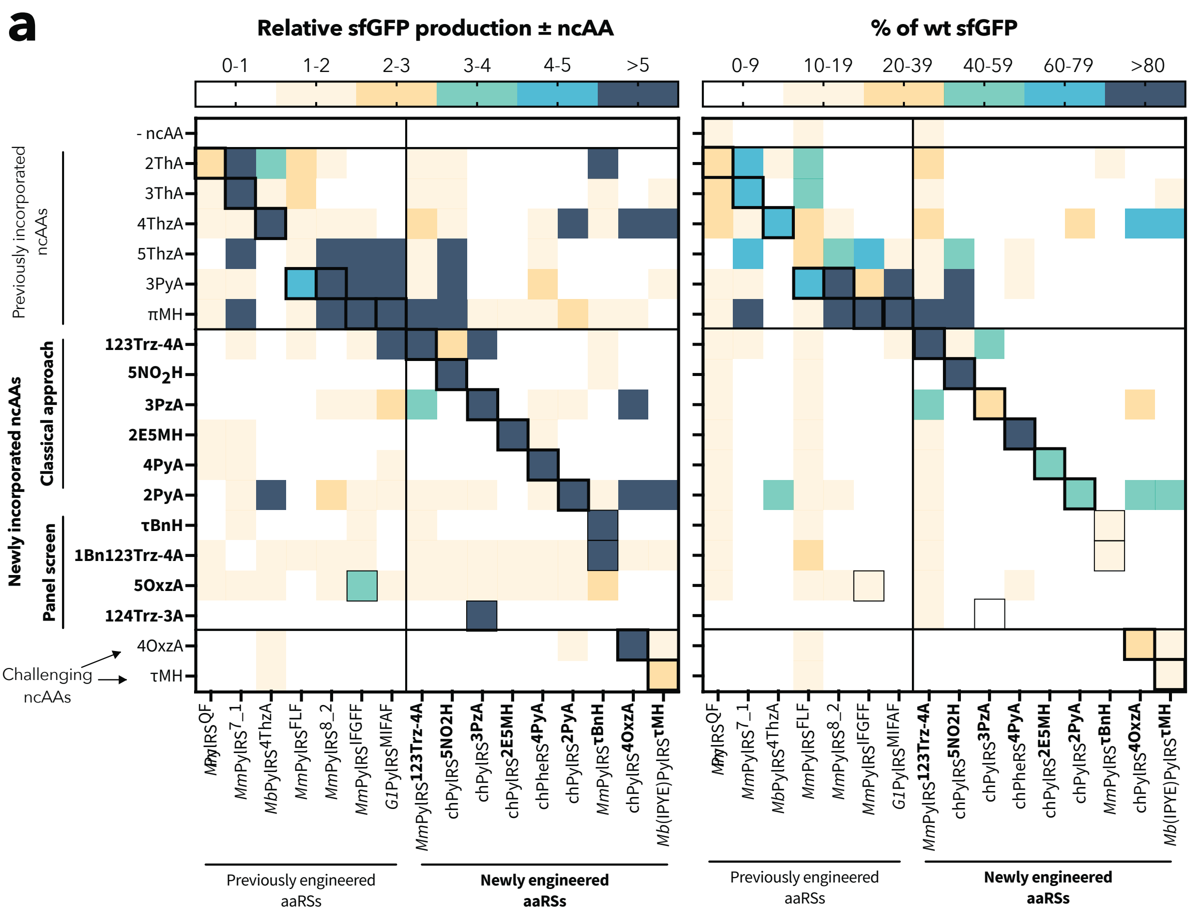


**Supplementary Figure 6 | aaRS-ncAA substrate specificity profiling.** a) substrate profiling analysis with relative sfGFP production ± ncAA (reproduction from Figure 2a for clear comparison) and % of wt sfGFP. Relative sfGFP production ± ncAA is calculated from the normalized fluorescence (excitation at 480 nm and emission at 510 nm normalized to the optical density at 600 nm) in the presence of the ncAA divided by the normalized fluorescence in the absence of the ncAA. The mean of 2-3 biological replicates is shown. Data indicated as % of wt sfGFP shows normalized fluorescence in the absence or presence of different ncAAs as percentage of a wt sfGFP reference. The mean of 2-3 biological replicates is shown.

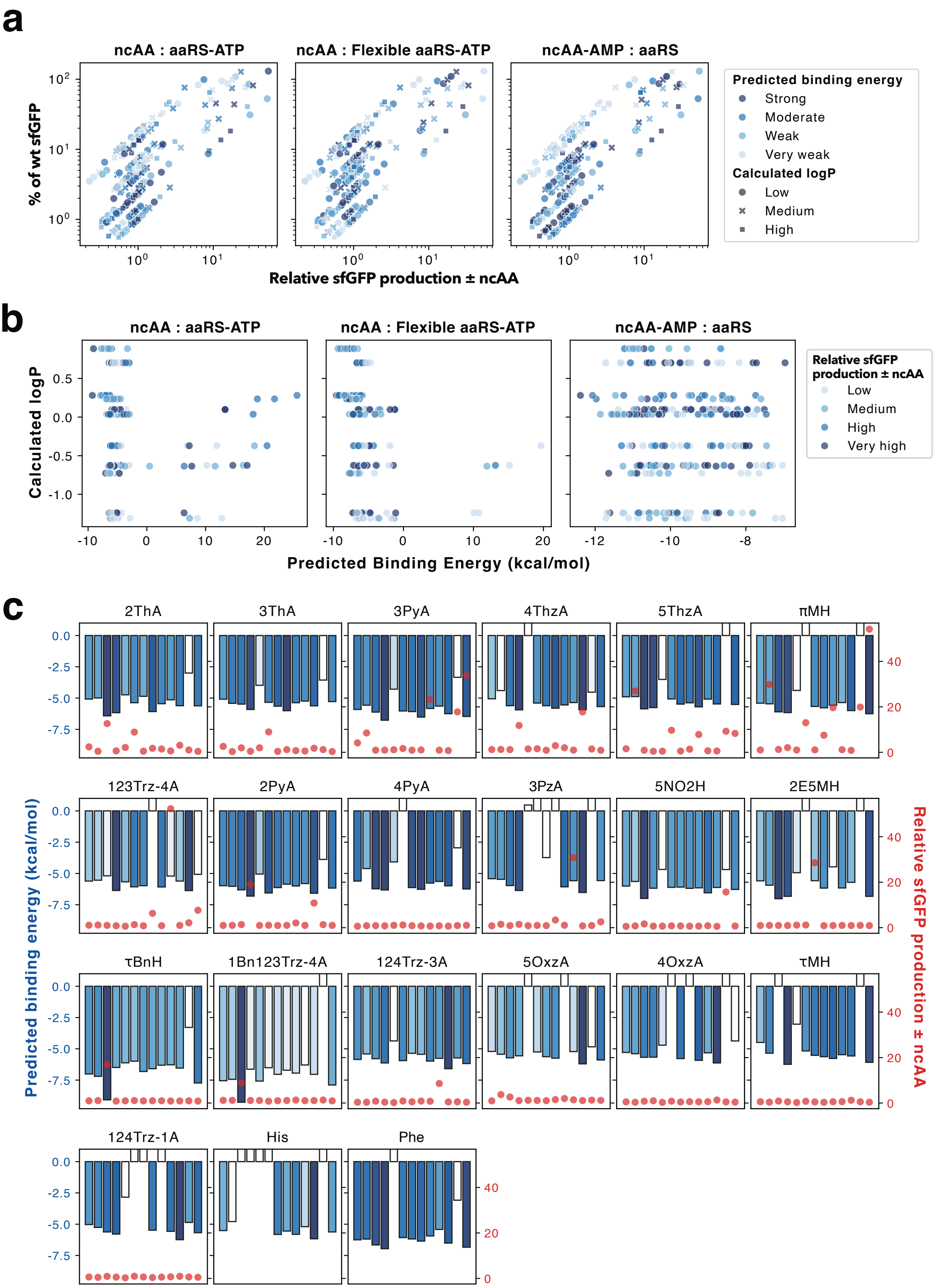


**Supplementary Figure 7 | Analysis of predicted affinities and sfGFP production metrics for different PylRS variants.** For each ncAA, the binding affinities of the ncAA to the aaRS-ATP adduct and the binding affinities of the ncAA-AMP adduct to the aaRS were predicted. The predictions were obtained from aaRS-ATP structure prediction with Boltz2[46] and docking of the ncAA with Gnina[56]. For binding of the ncAA to the aaRS-ATP adduct both a frozen protein configuration and a flexible protein configuration for residues within 3.5 Å of the substrate binding pocket were considered. The predicted binding affinities were visualized in three different methods: a) the relative sfGFP production ± ncAA vs. % of wt sfGFP with color coding indicating predicted binding energy and marker style indicating the calculated logP value, b) the calculated logP vs. the predicted binding energy with coloring indicating the relative sfGFP production ± ncAA, and c) separate subplots for each ncAA for the ncAA:aaRS-ATP binding analysis, with bars (left y-axis) and scatter circles (right y-axis) indicating the predicted binding energies and the relative GFP production ± ncAA, respectively. Because the differences in many of the binding energy predictions are not large, colors of the bars correspond to the predicted binding energies to aid in finding the most negative energies quickly. No clear correlation between predicted binding energies and sfGFP production metrics could be found, even accounting for logP.

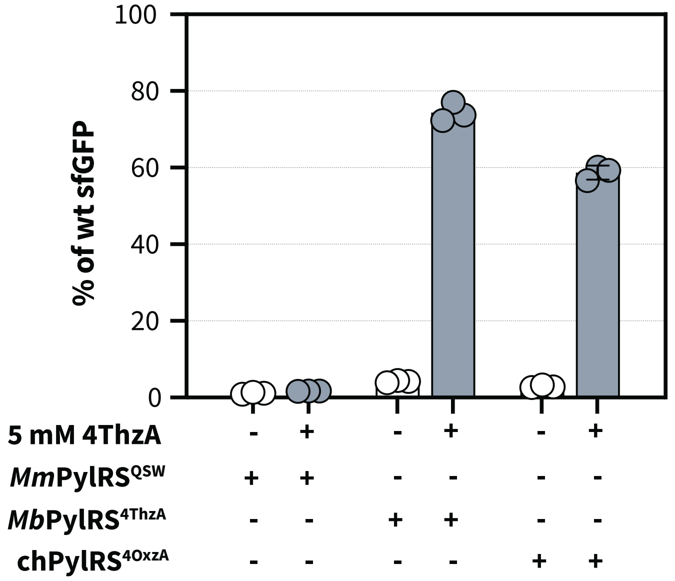


**Supplementary Figure 8 | Comparison of different PylRS variants for 4ThzA incorporation.** Suppression of sfGFP150$_{TAG}$ in NEB10β in the presence or absence of 5 mM 4ThzA with different PylRS variants. Data shows normalized fluorescence (excitation at 480 nm and emission at 510 nm) is normalized to the optical density at 600 nm) as percentage of a wt sfGFP reference. The mean and standard deviation of 3 biological replicates is shown.

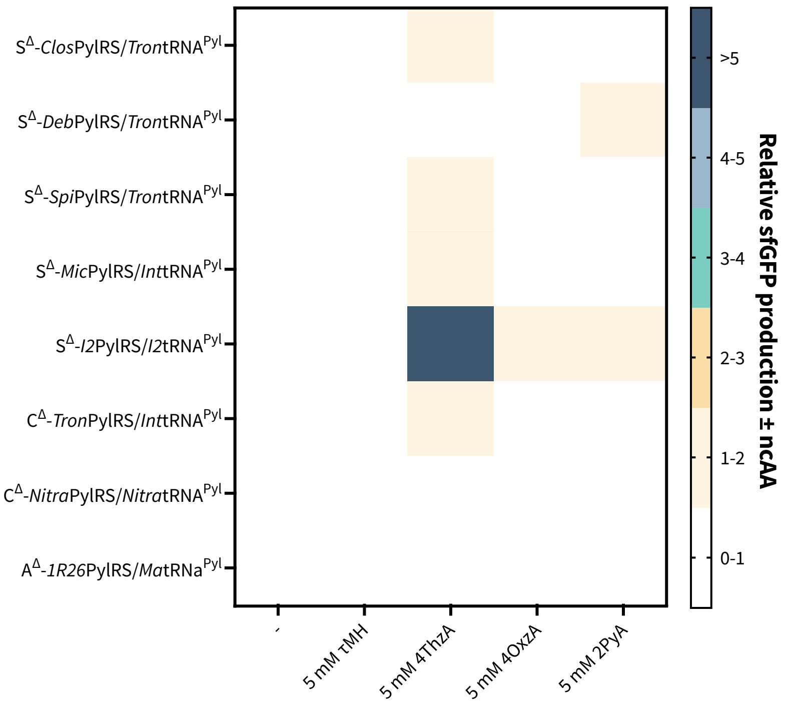


**Supplementary Figure 9| Testing of a broader range of PylRS$^{QSW}$ variants.** Suppression of sfGFP150$_{TAG}$ in NEB10β in the presence or absence of different ncAAs with different PylRS variants[51] carrying the "QSW" mutations. Data shows relative sfGFP production with and without a given ncAA. Relative sfGFP production ± ncAA is calculated from the normalized fluorescence (excitation at 480 nm and emission at 510 nm normalized to the optical density at 600 nm) in the presence of the ncAA divided by the normalized fluorescence in the absence of the ncAA. The mean of 3 biological replicates is shown.

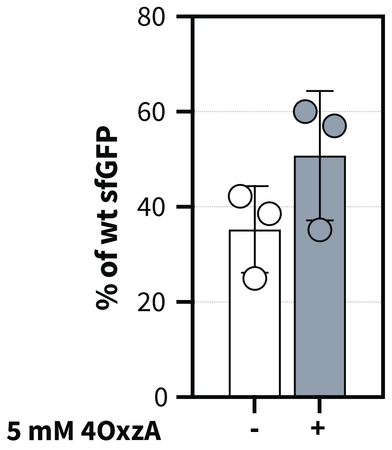


**Supplementary Figure 10 | Validation of chPylRS$^{4OxzA-2}$ for 4OxzA incorporation in sfGFP150$_{TAG}$.** a) Suppression of sfGFP150$_{TAG}$ in NEB10β in the presence or absence of 5 mM 4OxzA. Data shows normalized fluorescence (excitation at 480 nm and emission at 510 nm) is normalized to the optical density at 600 nm) as percentage of a wt sfGFP reference. The mean and standard deviation of 3 biological replicates is shown.

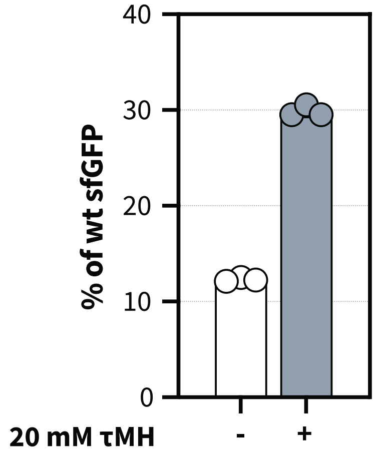


**Supplementary Figure 11 | Validation of *Mb*(IPYE)PylRS$^{P10}$ for τMH incorporation in sfGFP150$_{TAG}$.** Suppression of sfGFP150$_{TAG}$ in NEB10β in the presence or absence of 20 mM τMH. Data shows normalized fluorescence (excitation at 480 nm and emission at 510 nm) is normalized to the optical density at 600 nm) as percentage of a wt sfGFP reference. The mean and standard deviation of 3 biological replicates is shown.

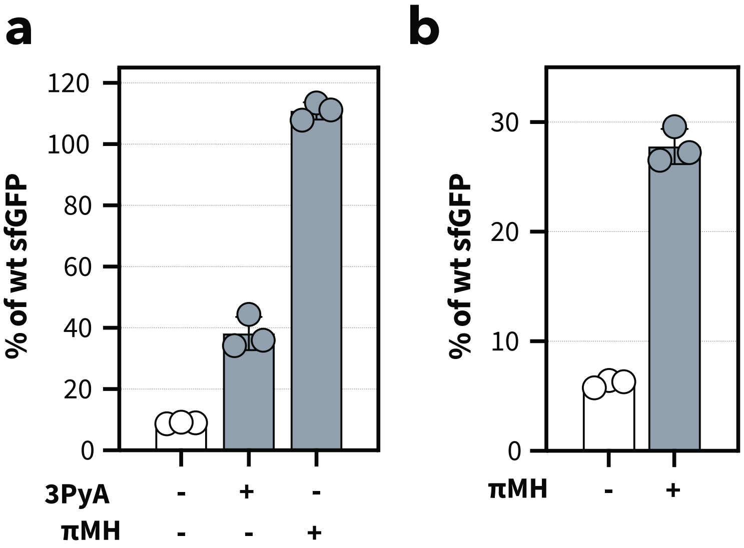


**Supplementary Figure 12 | Validation of opal suppression with PylRS pairs.** a) Suppression of sfGFP150$_{TGA}$ in NEB10β in the presence or absence of different ncAAs with *G1*PylRS$^{MIFAF}$/*Ma*tRNA$^{ΔNPyl}$(8)$_{UCA}$. b) Suppression of sfGFP150$_{TGA}$ in NEB10β in the presence or absence of πMH with *Ma*PylRS$^{IFGFF}$/*Ma*tRNA$^{ΔNPyl}$(8)$_{UCA}$. Data shows normalized fluorescence (excitation at 480 nm and emission at 510 nm) is normalized to the optical density at 600 nm) as percentage of a wt sfGFP reference. The mean and standard deviation of 3 biological replicates is shown.

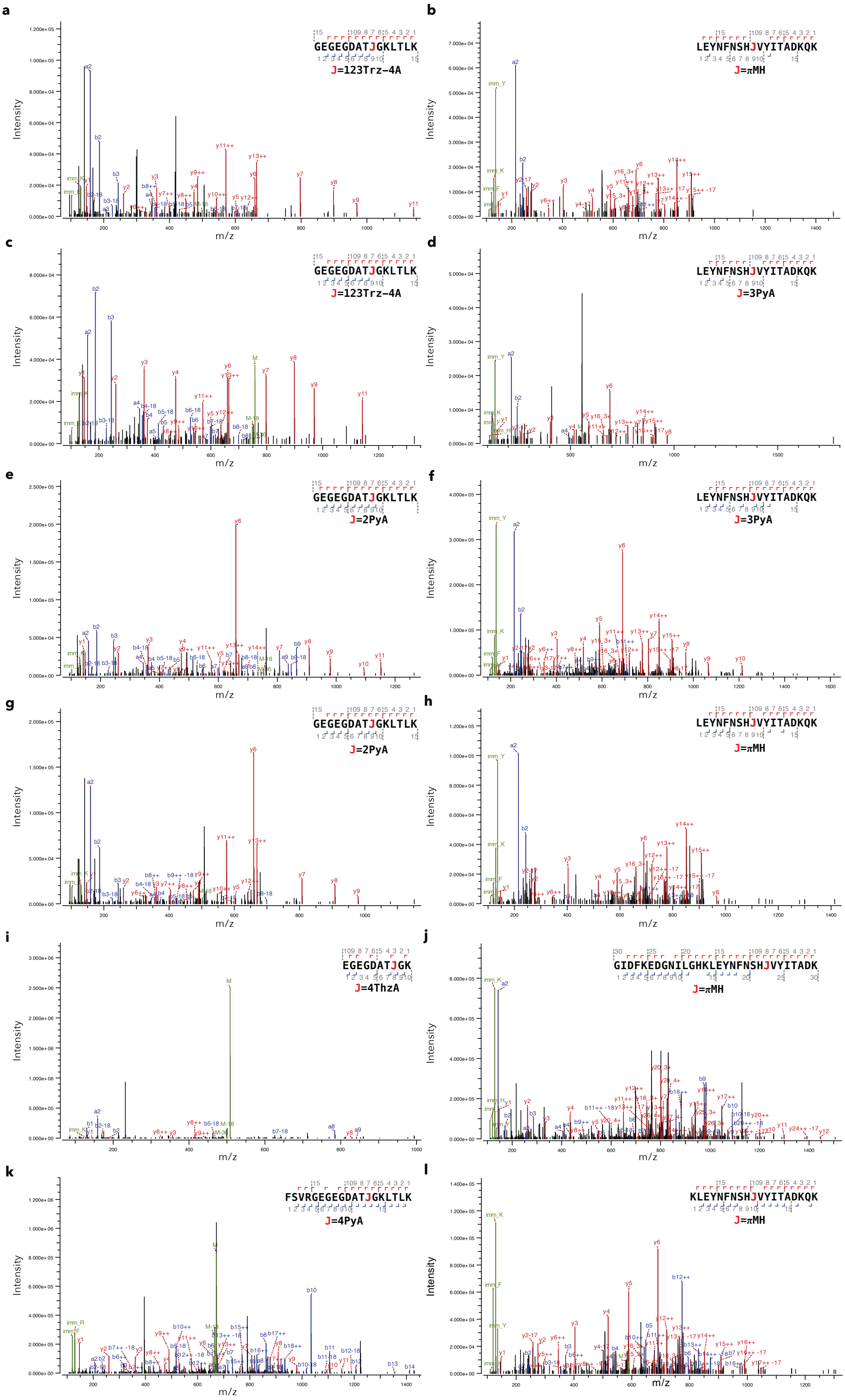


**Supplementary Figure 13 | LC-MS/MS spectra of sfGFP40$_{TAG}$150$_{TGA}$ containing two histidine-like ncAA.** Representative LC-MS/MS spectra from tryptic digests of sfGFP40$_{TAG}$150$_{TGA}$ expressed with different combinations of aaRS/tRNA pairs. In all cases, peptides containing the desired ncAA were observed and no peptides containing an undesired ncAA at the respective positions were detected. In the samples sfGFP40$_{123Trz-4A}$150$_{\pi MH}$, sfGFP40$_{2PyA}$150$_{3PyA}$ and sfGFP40$_{2PyA}$150$_{\pi MH}$, some Gln incorporation was detected. In the sfGFP40$_{4ThzA}$150$_{\pi MH}$, some amount of Phe incorporation was detected. a,b) sfGFP40$_{123Trz-4A}$150$_{\pi MH}$; c,d) sfGFP40$_{123Trz-4A}$150$_{3PyA}$; e,f) sfGFP40$_{2PyA}$150$_{3PyA}$; g,h) sfGFP40$_{2PyA}$150$_{3PyA}$; sfGFP40$_{2PyA}$150$_{\pi MH}$; i,j) sfGFP40$_{4ThzA}$150$_{\pi MH}$; k,l) sfGFP40$_{4PyA}$150$_{\pi MH}$.

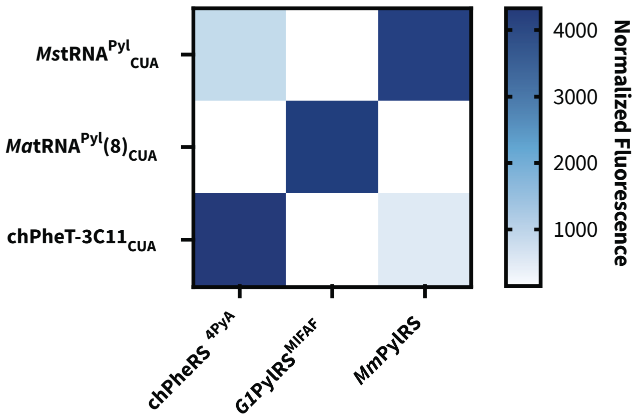


**Supplementary Figure 14 | Orthogonality testing of chPheRS$^{4PyA}$/3C11-chPheT$_{CUA}$ construct.** Suppression of sfGFP150$_{TAG}$ in NEB10β for different aaRS-tRNA combinations (chPheRS$^{4PyA}$ with 4 mM dl-4PyA, *G1*PylRS$^{MIFAF}$ with 2 mM πMH, *Mm*PylRS with 4 mM BocK). Data shows normalized fluorescence (excitation at 480 nm and emission at 510 nm is normalized to the optical density at 600 nm). The mean of 3 biological replicates is shown.

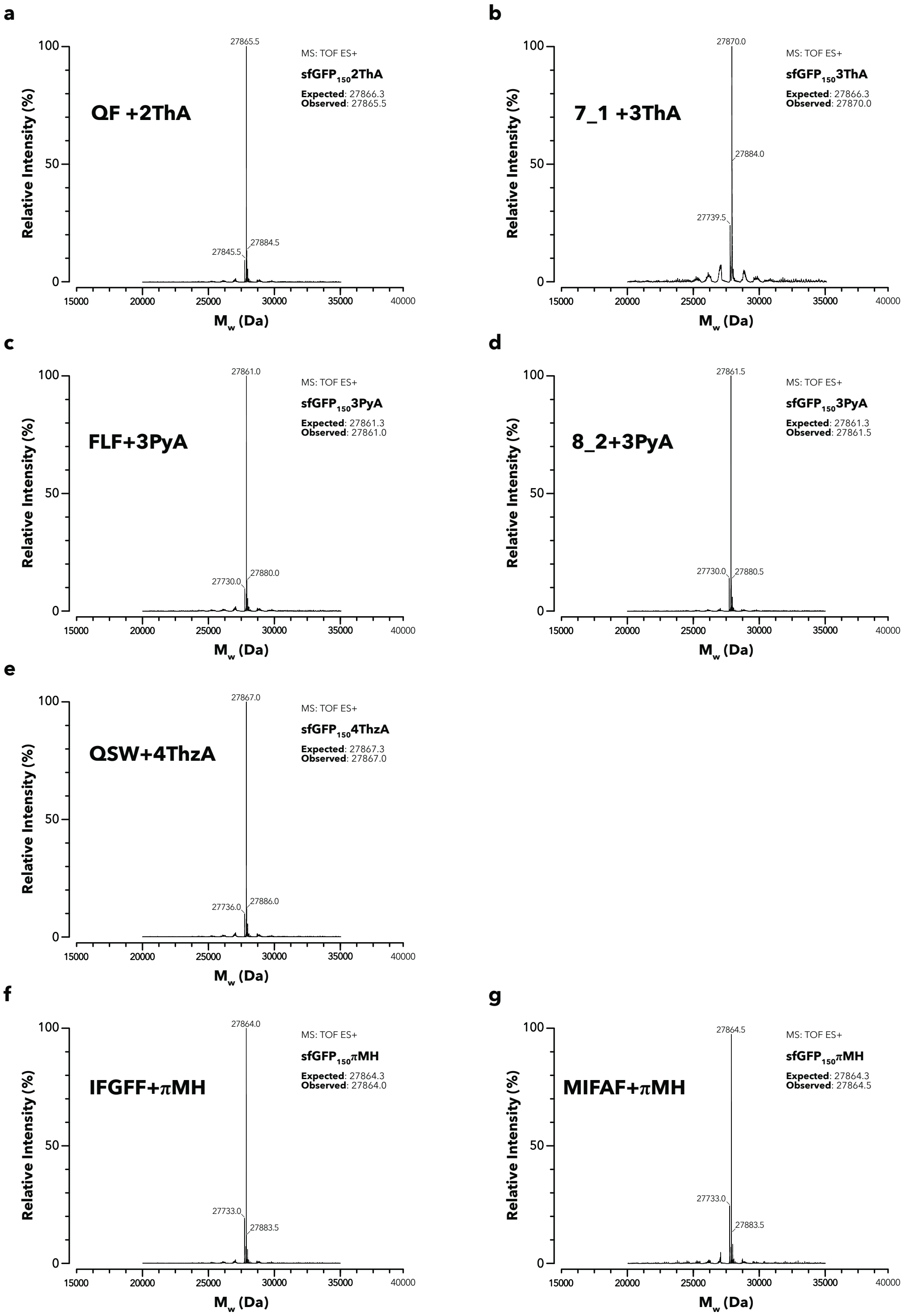


**Supplementary Figure 15 | LC-MS spectra of sfGfP150$_{TAG}$ containing a single histidine-like ncAA incorporated by previously reported aaRS/tRNA pairs.** Intact LC-MS analysis (positive electrospray time of flight) of sfGFP150$_{TAG}$ production in the presence of ncAAs with their cognate aaRS/tRNA pair. a) *Mb*PylRS$^{QF}$ with 2ThA. b) *Mm*PylRS$^{7_1}$ with 3ThA. c) *Mm*PylRS$^{FLF}$ with 3PyA. d) *Mm*PylRS$^{8_2}$ with 3PyA. e) *Mb*PylRS$^{4ThzA}$ with 4ThzA. f) *Mm*PylRS$^{IFGFF}$ with πMH. g) *G1*PylRS$^{MIFAF}$ with πMH. The expected and observed masses confirming incorporation of the desired ncAA are indicated. Additional peaks, consistent with typically observed [$H_2O$] elimination (-18 Da), $[Na]^+$ addition (+23 Da) or loss of N-terminal [Met] (-131 Da) were observed.

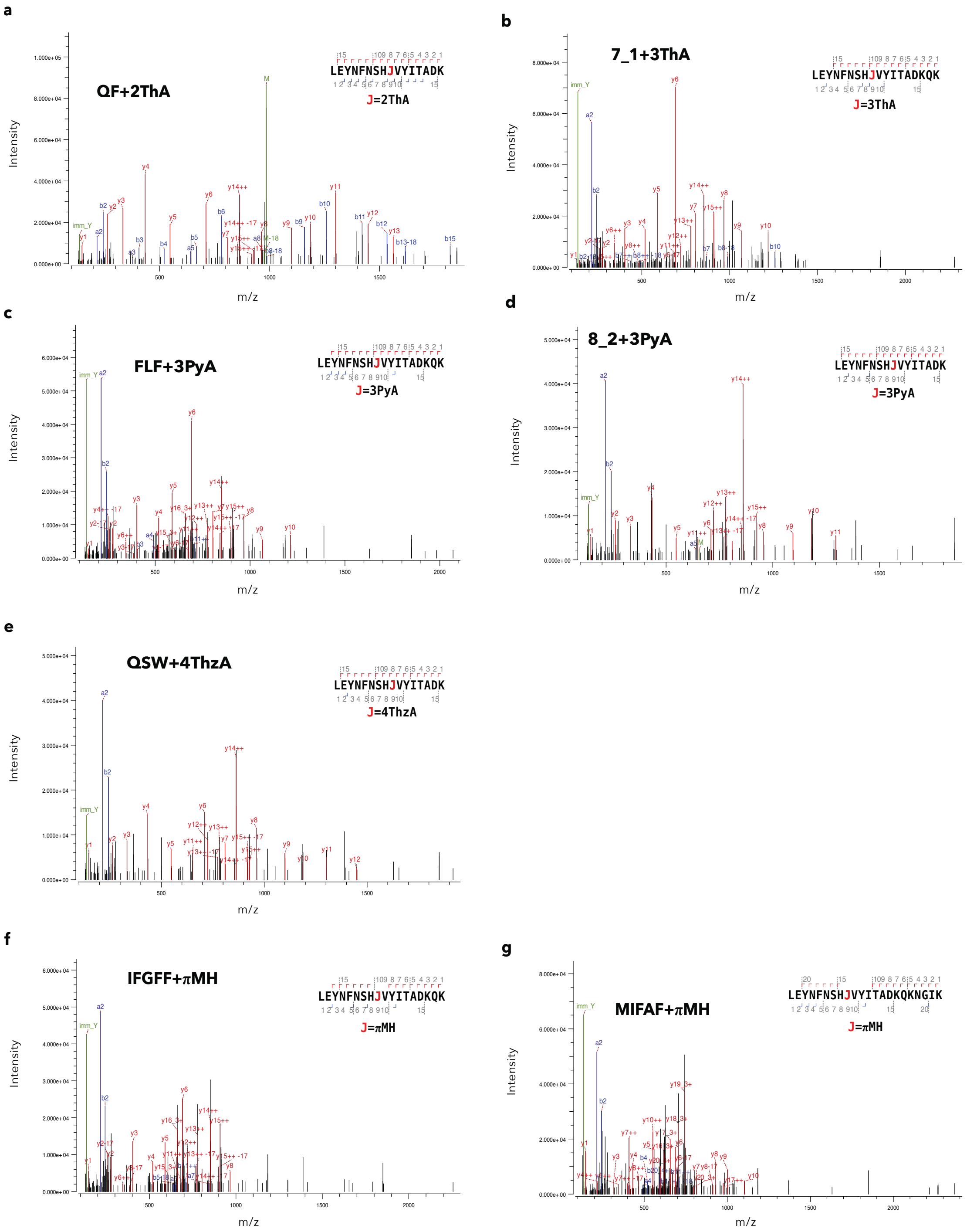


**Supplementary Figure 16 | LC-MS/MS spectra of sfGFP150$_{TAG}$ containing a single histidine-like ncAA incorporated by previously reported aaRS/tRNA pairs.** A representative LC-MS/MS spectrum from a tryptic digest of sfGFP150$_{TAG}$ expressed with different ncAAs and their corresponding aaRS/tRNA pairs. a) *Mb*PylRS$^{QF}$ with 2ThA. b) *Mm*PylRS$^{7_1}$ with 3ThA. c) *Mm*PylRS$^{FLF}$ with 3PyA. d) *Mm*PylRS$^{8_2}$ with 3PyA. e) *Mb*PylRS$^{4ThzA}$ with 4ThzA. f) *Mm*PylRS$^{IFGFF}$ with πMH. g) *G1*PylRS$^{MIFAF}$ with πMH. Typically, multiple peptides containing the desired ncAA were observed and no peptides for canonical amino acid incorporation were observed. For samples of sfGFP150-3ThA and both aaRSs of sfGFP150-3PyA, some peptides containing canonical amino acids were observed (see **Supplementary Table 9**).

### II. Synthesis

#### 1,2,4-Triazol-3-yl-alanine hydrochloride synthesis (124Trz-3A)

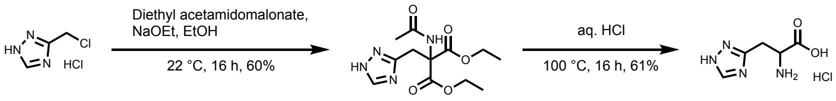


Diethyl-2-acetamido-2-(1,2,4-triazol-3-ylmethyl)malonate

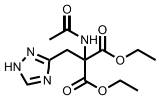

The reaction was performed under a nitrogen atmosphere. Sodium ethoxide (21% in EtOH, 3.6 mL, 9.7 mmol, 3.8 eq.) was dissolved in dry EtOH (2 mL) and cooled to 0 °C. Diethyl acetamidomalonate (1.11 g, 5.10 mmol, 2.0 eq.) was dissolved in dry EtOH (13 mL) and was added dropwise to the sodium ethoxide solution. The reaction mixture was stirred at room temperature for 2 h. 1,2,3-Triazol-3-ylmethylchloride hydrochloride (393 mg, 2.55 mmol, 1.0 eq.) was dissolved in dry EtOH (6 mL) and added to the malonate solution. The reaction mixture was stirred at room temperature for 16 h and quenched with the addition of water. The solvents were removed *in vacuo* and the crude material was dissolved in water. The solution was adjusted to pH 6 with aq. HCl (4 M) and was extracted with EtOAc (6x). The combined organic phase was dried over $Na_2SO_4$ and the solvent was removed *in vacuo* to yield diethyl-2-acetamido-2-(1,2,4-triazol-3-ylmethyl)malonate (456 mg, 1.53 mmol, 60%) as a yellow solid.

$^1$H-NMR (400 MHz, $CDCl_3$) δ 7.96 (s, 1H), 6.87 (s, 1H), 4.36 – 4.20 (m, 4H), 3.87 (s, 2H), 1.99 (s, 3H), 1.26 (t, $^3J_{HH}$ = 7.1 Hz, 6H).

1,2,4-Triazol-3-yl-alanine hydrochloride (124Trz-3A)

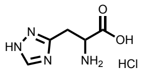

Diethyl-2-acetamido-2-(1,2,4-triazol-3-ylmethyl)malonate (2.11 g, 7.06 mmol, 1.0 eq.) was suspended in aq. HCl (4 M, 36 mL) and stirred at 100 °C for 16 h. The solvent was removed *in vacuo*, the crude product was dissolved in EtOH (14 mL) and aniline (700 µL) was added to precipitate 1,2,4-triazol-3-yl-alanine hydrochloride (836 mg, 4.28 mmol, 61%) as a beige solid with aniline as an impurity.[62]

$^1$H-NMR (400 MHz, $D_2O$) δ 8.37 (s, 1H), 4.15 (dd, $^3J_{HH}$ = 7.9, 5.0 Hz, 1H), 3.44 (dd, $^2J_{HH}$ = 16.0 Hz, $^3J_{HH}$ = 5.0 Hz, 1H), 3.33 (dd, $^2J_{HH}$ = 16.0 Hz, $^3J_{HH}$ = 7.9 Hz, 1H).

$^{13}$C NMR (101 MHz, $D_2O$) δ 172.7, 156.4, 145.9, 53.2, 28.0.

MS (ESI): calc. for $[M+H]^+$: 157.06, obs.: 157.1.

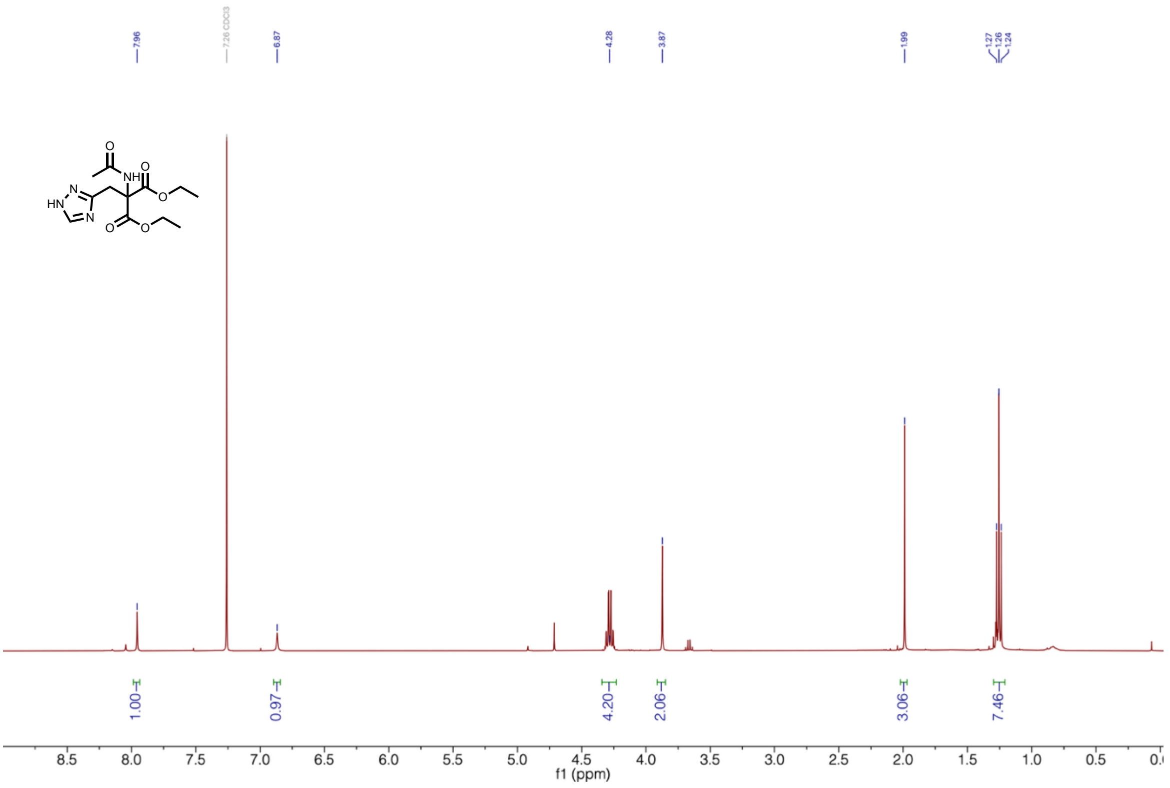

**Supplementary Figure 17 | $^{1}$H-NMR of diethyl-2-acetamido-2-(1,2,4-triazol-3-ylmethyl)malonate.**

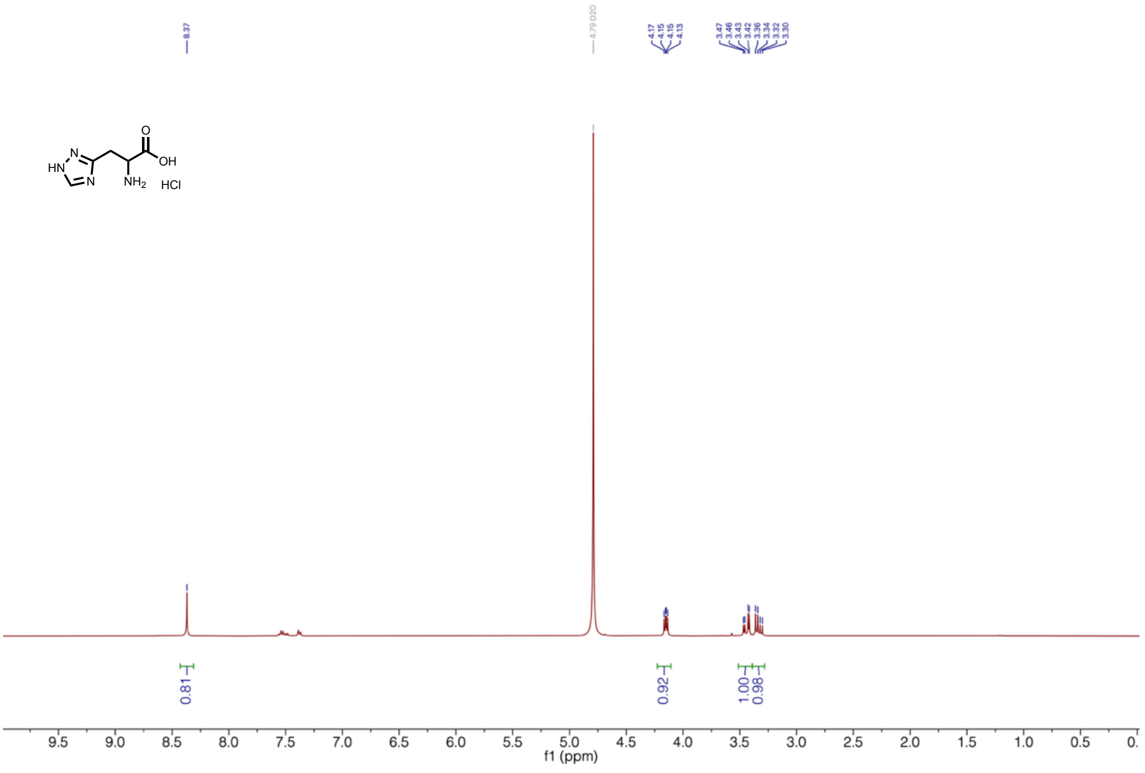

**Supplementary Figure 18 | $^{1}$H-NMR of 1,2,4-triazol-3-yl-alanine hydrochloride (124Trz-3A).**

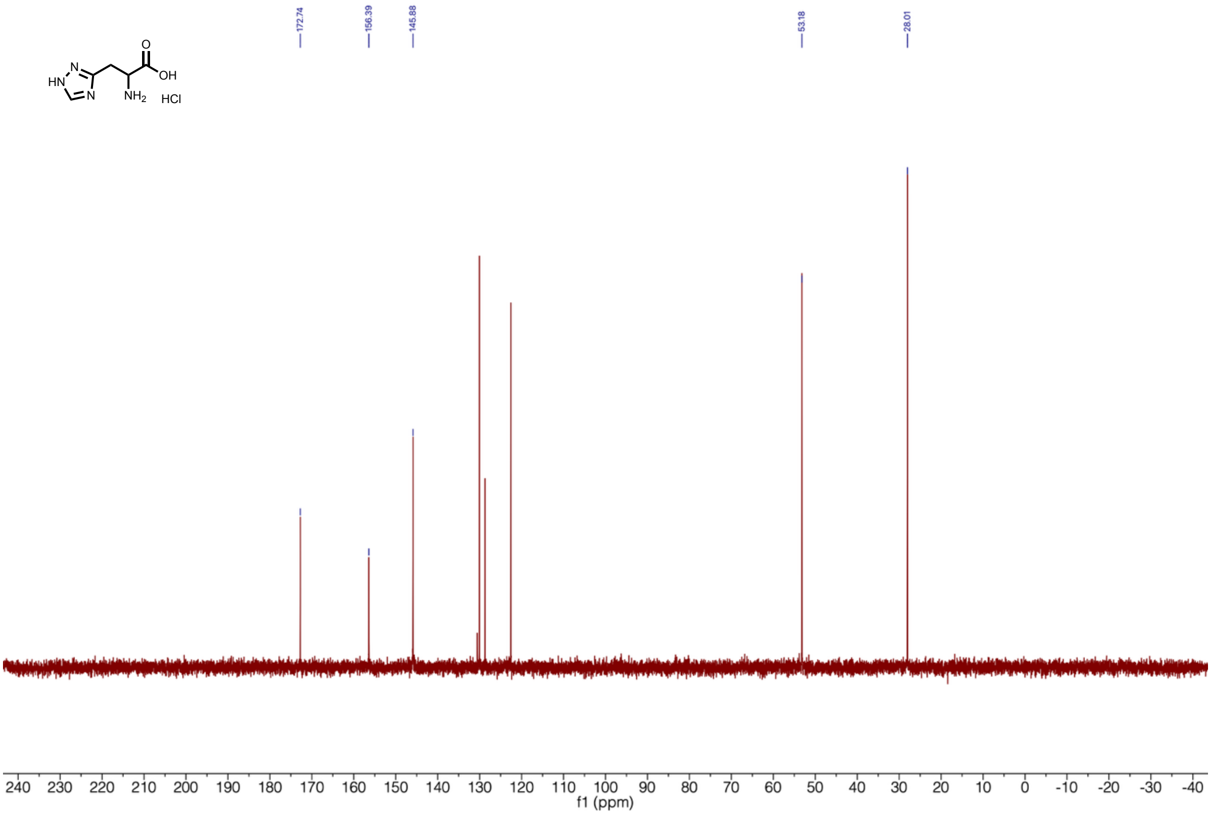


**Supplementary Figure 19 | $^{13}$C-NMR of 1,2,4-triazol-3-yl-alanine hydrochloride (124Trz-3A).**

**5-Nitro-L-histidine hydrochloride synthesis (5NO$_2$H):**

5-Nitro-L-histidine hydrochloride was synthesized as previously described[63].

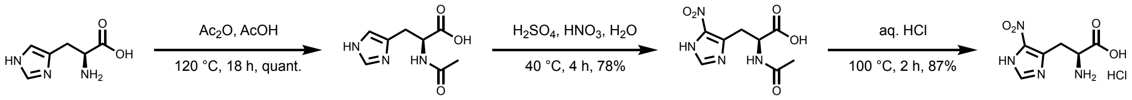


<u>*N*-acetyl-L-histidine:</u>

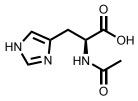

L-histidine (2.00 g, 12.8 mmol, 1.0 eq) was suspended in conc. acetic acid (10 mL). Acetic anhydride (1.2 mL, 13 mmol, 1.0 eq) was added and the solution was stirred at 120 °C for 18 h. The reaction mixture was allowed to reach room temperature and the solvent was removed *in vacuo*. The residue was resuspended in water (5 mL) and the solvent was removed *in* vacuo. This step was repeated two more times to yield *N*-acetyl-L-histidine (2.52 mg, 12.8 mmol, quant.) as a white powder.

<u>*N*-acetyl- 5-nitro-L-histidine:</u>

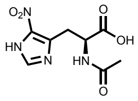

*N*-acetyl-L-histidine (2.52 g, 12.8 mol, 1.0 eq.) was suspended in conc. sulfuric acid (20 mL) and cooled to 0 °C. Aq. nitric acid (65%, 8 mL) was added dropwise and the reaction mixture was stirred at 40 °C for 4 h. The solution was allowed to reach room temperature and was poured into ice-cold water (100 mL). The pH was adjusted to pH 4 with aq. sat. $K_2CO_3$. The solvent was removed *in vacuo*. The crude product was resuspended in methanol, filtered, and the filtrate was concentrated *in vacuo* to yield *N*-acetyl-5-nitro-L-histidine (1.50 g, 7.49 mmol, 78%) as a colorless oil.

**5**<u>-Nitro-L-histidine hydrochloride (5NO2H):</u>

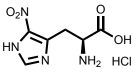


*N*-acetyl-5-nitro-L-histidine (1.00 g, 4.12 mmol, 1.0 eq.) was suspended in aq. HCl (2M, 10 mL) and stirred at 100 °C for 2h. The solvent was removed *in vacuo* and the crude product was purified by automated flash column chromatography (C18, 0 – 95% acetonitrile in water) to yield 5-nitro-L-histidine hydrochloride as an off white solid (850 mg, 3.59 mmol, 87%).

$^1$H-NMR (400 MHz, $D_2O$) δ 7.75 (s, 1H), 4.44 (dd, $^3J_{HH}$ = 7.2, 6.6 Hz, 1H), 3.73 (dd, $^2J_{HH}$ = 14.9 Hz, $^3J_{HH}$ = 7.2 Hz, 1H), 3.66 (dd, $^2J_{HH}$ = 14.9 Hz, $^3J_{HH}$ = 6.6 Hz, 1H).

$^{13}$C-NMR (101 MHz, $D_2O$) δ 170.5, 143.9, 135.1, 128.1, 51.7, 26.1.

MS (ESI): calc. for $[M+H]^+$: 201.05, obs.: 200.9.

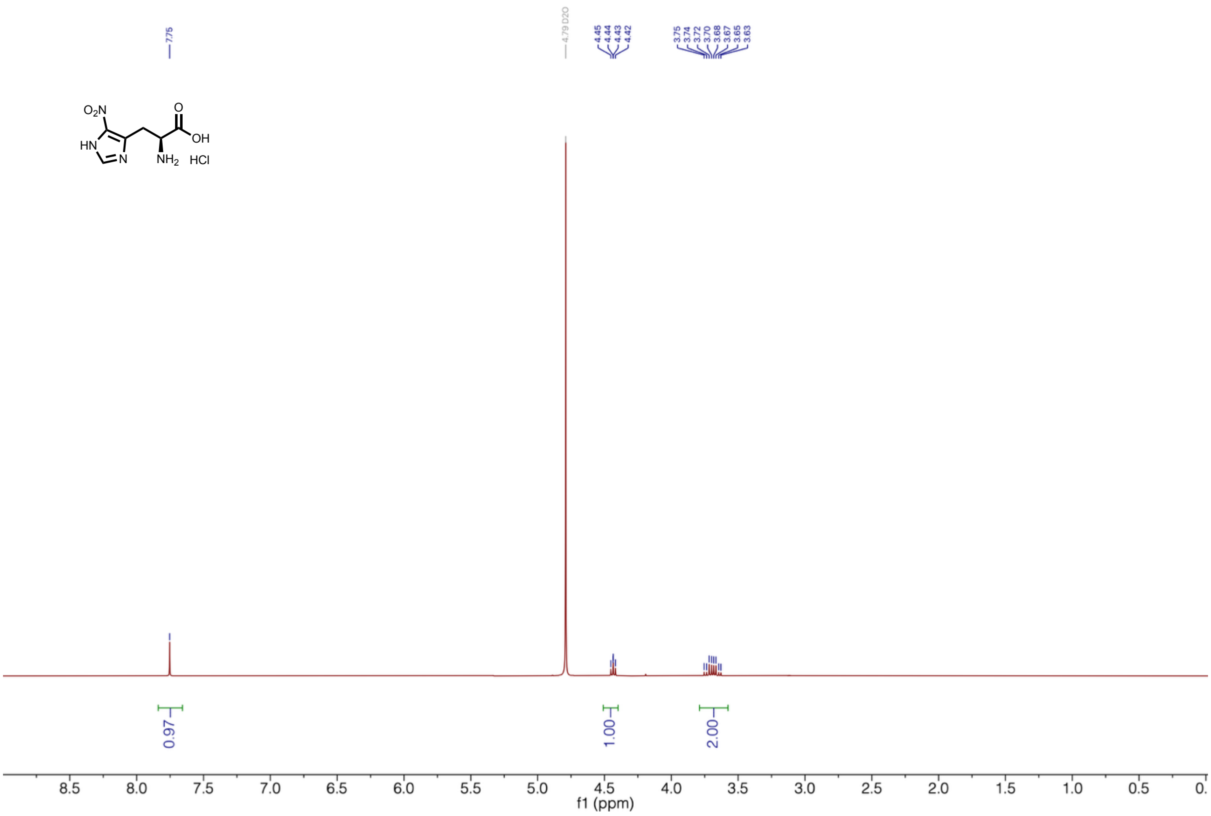


**Supplementary Figure 20 | $^1$H-NMR of 5-nitro-L-histidine hydrochloride ($5NO_2H$).**

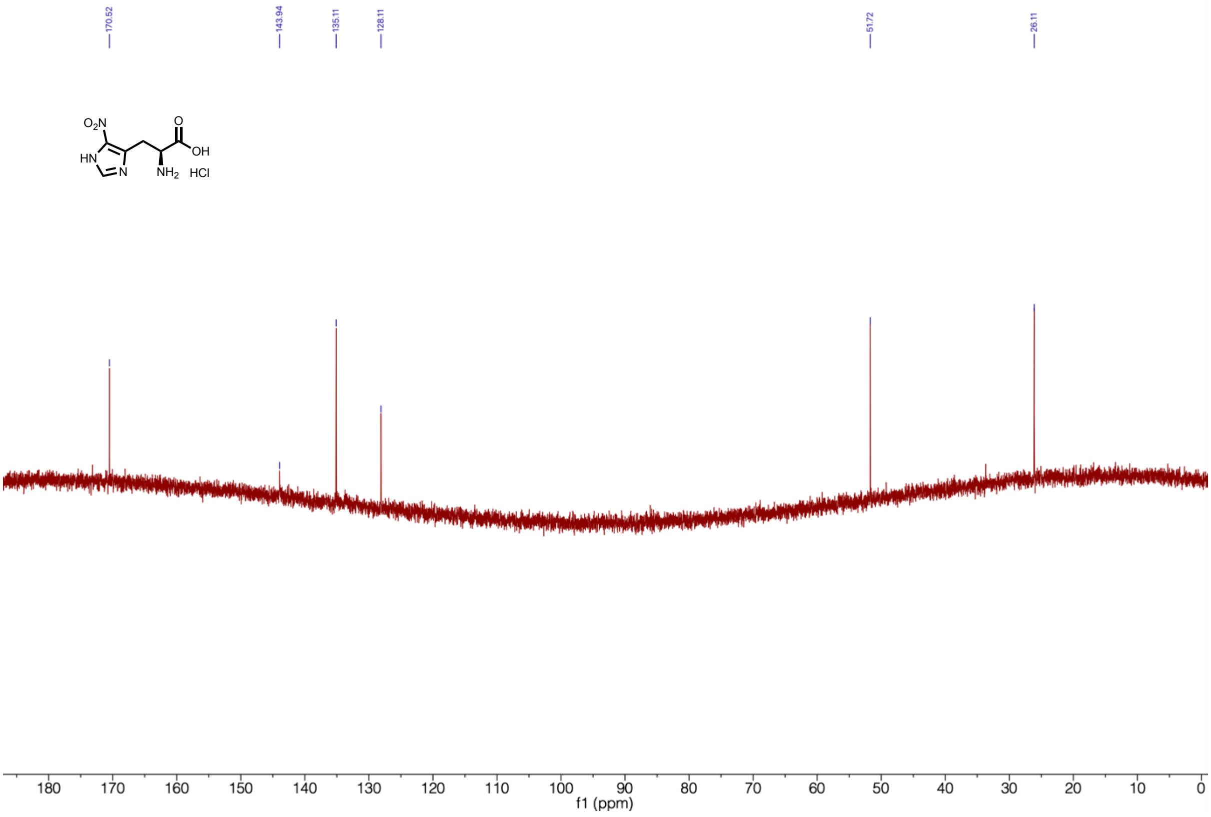

**Supplementary Figure 21 | $^{13}$C-NMR of 5-nitro-L-histidine hydrochloride (5NO$_2$H).**

**4-Pyridinyl-alanine hydrochloride synthesis (4PyA)**

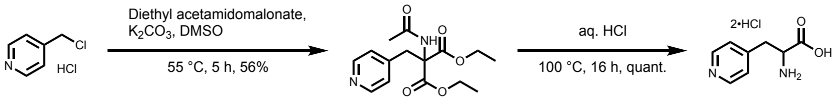


Diethyl-2-acetamido-2-(pyridin-4-ylmethyl) malonate:

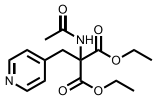

The following reaction was adapted from literature[64]. 4-Picolylchloride hydrochloride (2.18 g, 13.3 mmol, 1.0 eq.) was dissolved in DMSO (20 mL). Diethyl acetamidomalonate (2.91 g, 13.4 mmol, 1.0 eq) and $K_2CO_3$ (3.73 g, 27.0 mmol, 2.0 eq.) were added and the reaction was stirred at 55 °C for 5h. The reaction mixture was allowed to reach room temperature, was poured into ice-cold water (40 mL), and EtOAc (100 mL) was added. The mixture was centrifuged for 10 min at 4'000 g. The phases were separated and EtOAc (100 mL) was added to the aqueous phase. The mixture was centrifuged for 10 min at 4'000 g. The combined organic phase was washed with brine, dried over $MgSO_4$, and the solvent was removed *in vacuo* to yield diethyl-2-acetamido-2-(pyridin-4-ylmethyl)malonate (2.31 g, 7.49 mmol, 56%) as a brown solid.

$^1$H-NMR (400 MHz, DMSO-d$_6$) δ 8.49 – 8.44 (m, 2H), 7.03 – 6.96 (m, 2H), 4.21 – 4.10 (m, 4H), 3.44 (s, 2H), 2.54 (s, 3H), 1.17 (t, $^3J_{HH}$ = 7.1 Hz, 6H).

4-Pyridinyl-alanine hydrochloride:

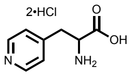


Diethyl-2-acetamido-2-(pyridin-4-ylmethl)malonate (1.45 g, 4.70 mmol, 1.0 eq.) was suspended in aq. HCl (4 M, 20 mL) and stirred at 100 °C for 16 h. The solvent was removed *in vacuo* and the crude product was purified by automated flash column chromatography (C18, 0-95% acetonitrile in water) to yield 4-pyridinyl alanine hydrochloride (953 mg, 4.70 mmol, quant.) as an off white solid.

$^1$H-NMR (400 MHz, $D_2O$) δ 8.76 (d, $^3J_{HH}$ = 6.8 Hz, 2H), 8.05 (d, $^3J_{HH}$ = 6.8 Hz, 2H), 4.47 – 4.41 (m, 1H), 3.66 – 3.52 (m, 2H).

$^{13}$C-NMR (101 MHz, $H_2O$+$D_2O$) δ 170.7, 156.8, 141.2, 128.2, 52.9, 35.8.

MS (ESI): calc. for $[M+H]^+$: 167.07, obs.: 167.2.

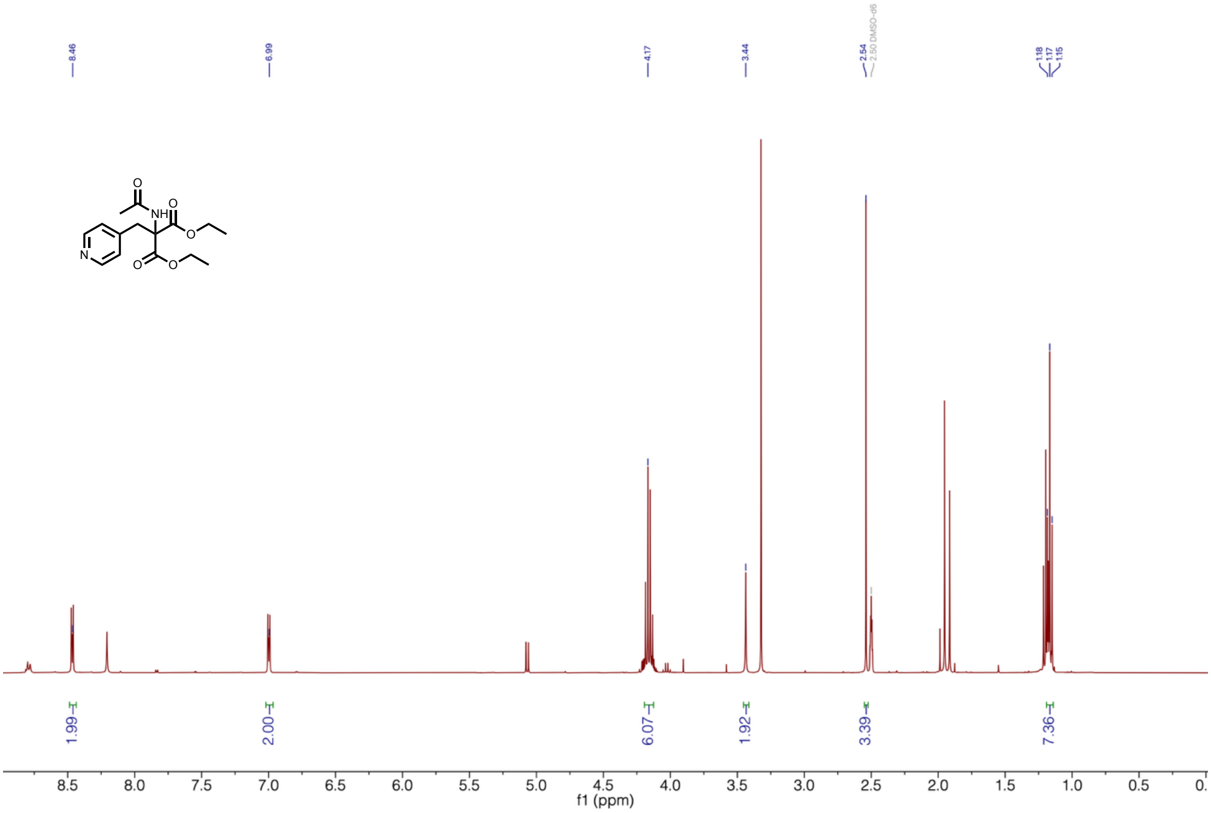


**Supplementary Figure 22 | $^1$H-NMR of diethyl-2-acetamido-2-(pyridin-4-ylmethyl) malonate.**

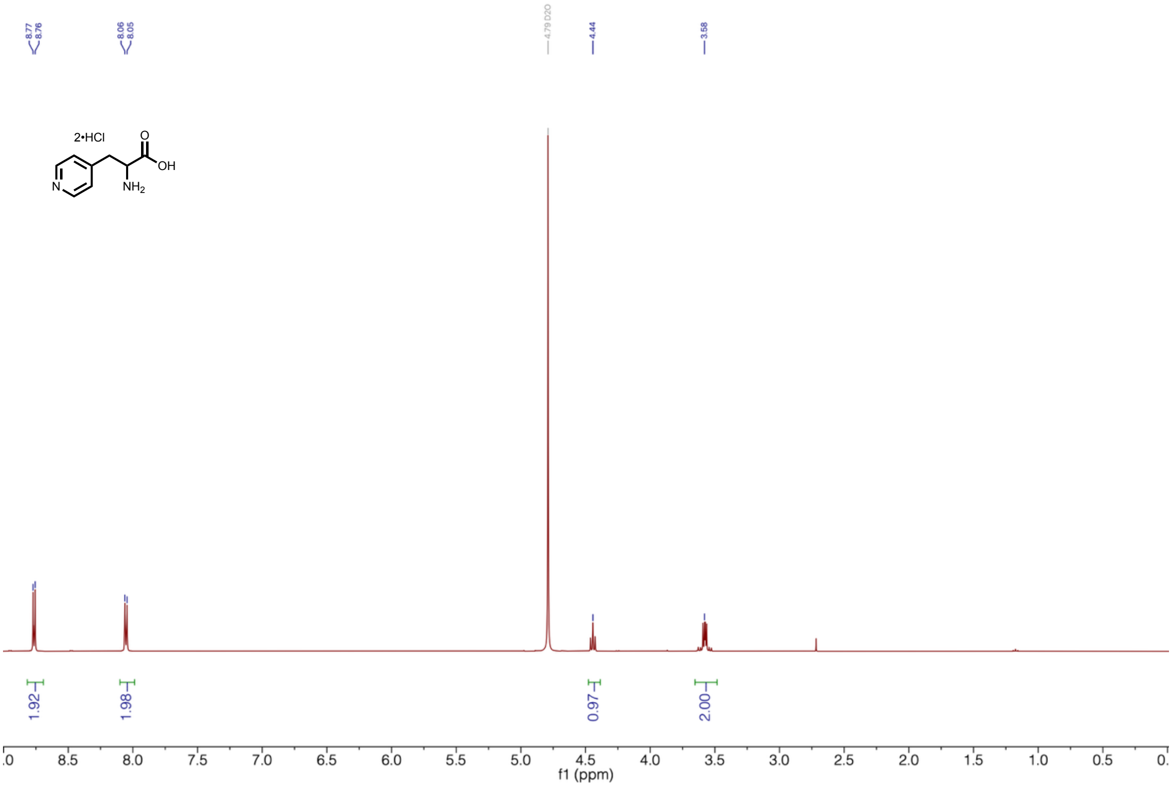


**Supplementary Figure 23 | $^{1}$H-NMR of 4-pyridinyl-alanine (4PyA).**

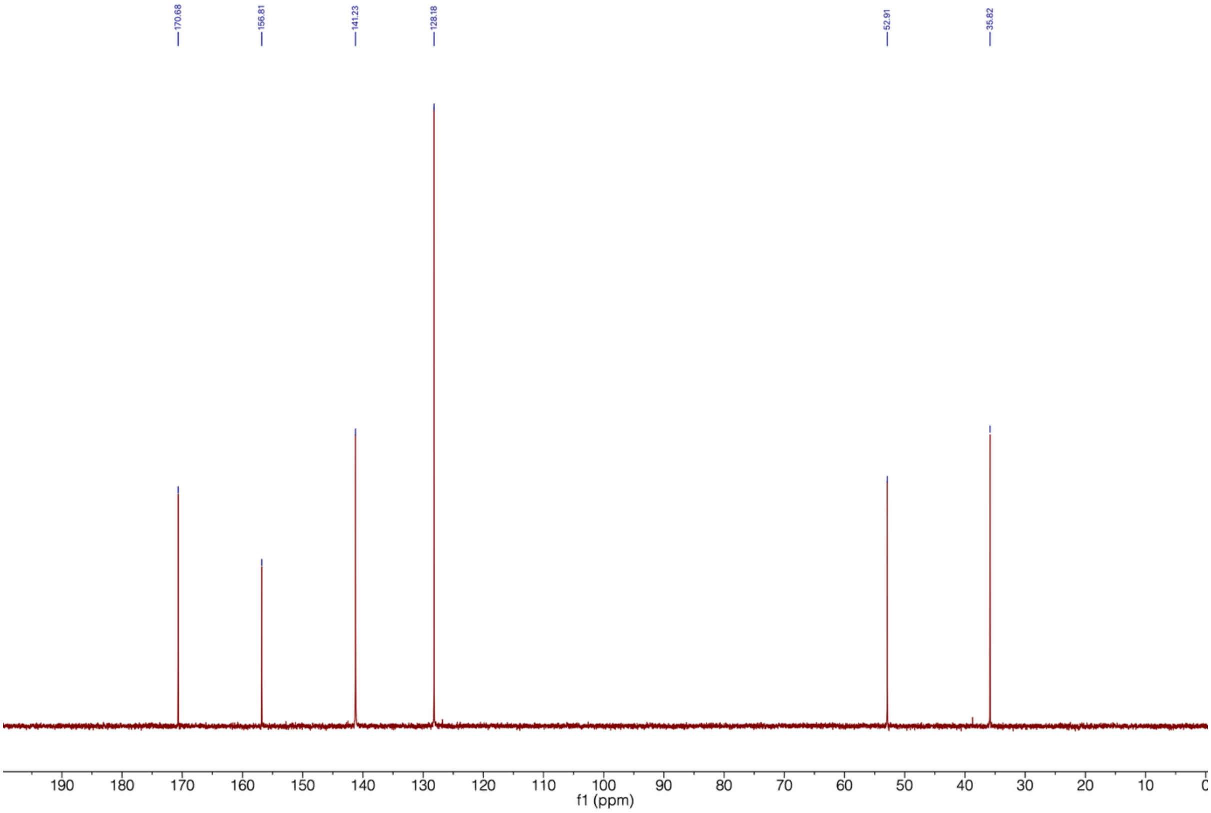


**Supplementary Figure 24 | $^{13}$C-NMR of 4-pyridinyl-alanine (4PyA).**

**1,2,3-Triazol-4-yl-L-alanine synthesis (123Trz-4A):**

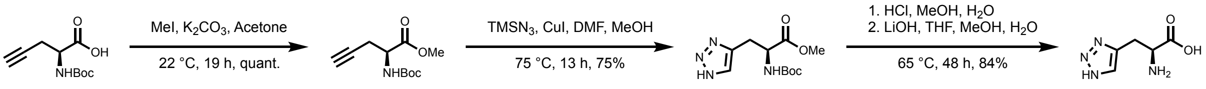


*N*-Boc-propargyl-L-glycine-OMe

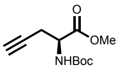

The following reaction was adapted from literature[65]. *N*-Boc-propargylglycine (5.00 g, 22.7 mmol, 1.0 eq.) was dissolved in acetone (100 mL). $K_2CO_3$ (6.37 g, 46.1 g, 2.0 eq.) and methyl iodide (2.9 mL, 46 mmol, 2.0 eq.) were added and the suspension was stirred at room temperature for 19 h. The solvent was removed *in vacuo* and the crude product was dissolved in ethyl acetate. The organic phase was washed with water and brine and dried over $Na_2SO_4$. The solvent was removed *in vacuo* to yield *N*-boc-propargyl-L-glycine-OMe (5.36 g, 22.7 mmol, quant.) as a yellow-orange oil.

$^1$H-NMR (400 MHz, $CDCl_3$) δ 5.34 (d, $^3J_{HH}$ = 8.6 Hz, 1H), 4.54 – 4.42 (m, 1H), 3.78 (s, 3H), 2.80 – 2.66 (m, 2H), 2.04 (t, $^4J_{HH}$ = 2.0 Hz, 1H), 1.45 (s, 9H).

*N*-Boc-1,2,3-triazol-4-yl-L-alanine-OMe

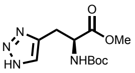

*N*-Boc-propargyl-L-glycine-OMe (1.10 g, 4.84 mmol, 1.0 eq.) was dissolved in dry dimethylformamide (5 mL) and dry methanol (1 mL) under an inert atmosphere. Trimethylsilylazide (0.97 mL, 7.3 mmol, 1.5 eq.) was added. Copper iodide (86 mg, 10 mol%) was added and the reaction mixture was stirred at 60 °C for 5 h and then at 75 °C for 13 h. The suspension was allowed to reach room temperature and was then further cooled to 4 °C. The reaction mixture was quenched by the addition of water and the solution was extracted with EtOAc (2x 50 mL). The combined organic phase was dried over $Na_2SO_4$ and the solvent was removed *in vacuo*. The crude product was purified by flash column chromatography ($SiO_2$, 50-100% ethyl acetate in cyclohexane) to yield *N*-Boc-1,2,3-triazol-4-yl-L-alanine-OMe (983 mg, 4.84 mmol, 75%) as a yellow sticky oil.

$R_f$ (cyclohexane:ethyl acetate, 1:1): 0.2, brown spot with ninhydrin stain.

$^1$H-NMR (400 MHz, $CDCl_3$) δ 7.52 (s, 1H), 5.43 (d, $^3J_{HH}$ = 8.2 Hz, 1H), 4.72 – 4.62 (m, 1H), 3.74 (s, 3H), 3.30 – 3.23 (m, 2H), 1.43 (s, 9H).

1,2,3-Triazol-4-yl-L-alanine (123Trz-4A)

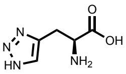

*N*-Boc-1,2,3-triazol-4-yl-alanine-OMe (2.95 g, 10.9 mmol, 1.0 eq.) was dissolved in methanol (70 mL) and aq. HCl (conc., 4.5 mL, 54 mmol, 5.0 eq.) was added. The solution was stirred at 40 °C for 17 h, followed by 50 °C for 4 h. The solvent was removed *in vacuo* to yield an orange-brown oil. The oil was dissolved in a mixture of THF, methanol, and water (15 mL each) and LiOH•$H_2O$ (1.15 g, 42.0 mmol,

2.5 eq.) was added. The suspension was stirred at 65 °C for 2 days. The solvents were removed *in vacuo* and the crude product was purified by preparative HPLC chromatography (C18, 50-90% acetonitrile in water (0.1% formic acid)) to yield 1,2,3-triazol-4-yl-L-alanine (1.43 g, 10.9 mmol, 84%) as a white solid.

$^1$H-NMR (400 MHz, $D_2O$) δ 7.82 (s, 1H), 4.10 (dd, $^3J_{HH}$ = 6.8, 5.3 Hz, 1H), 3.43 – 3.31 (m, 1H).

$^{13}$C-NMR (101 MHz, $D_2O$) δ 173.0, 140.2, 127.1, 54.2, 25.9.

MS (ESI): calc. for $[M+H]^+$: 157.06, obs.: 157.1.

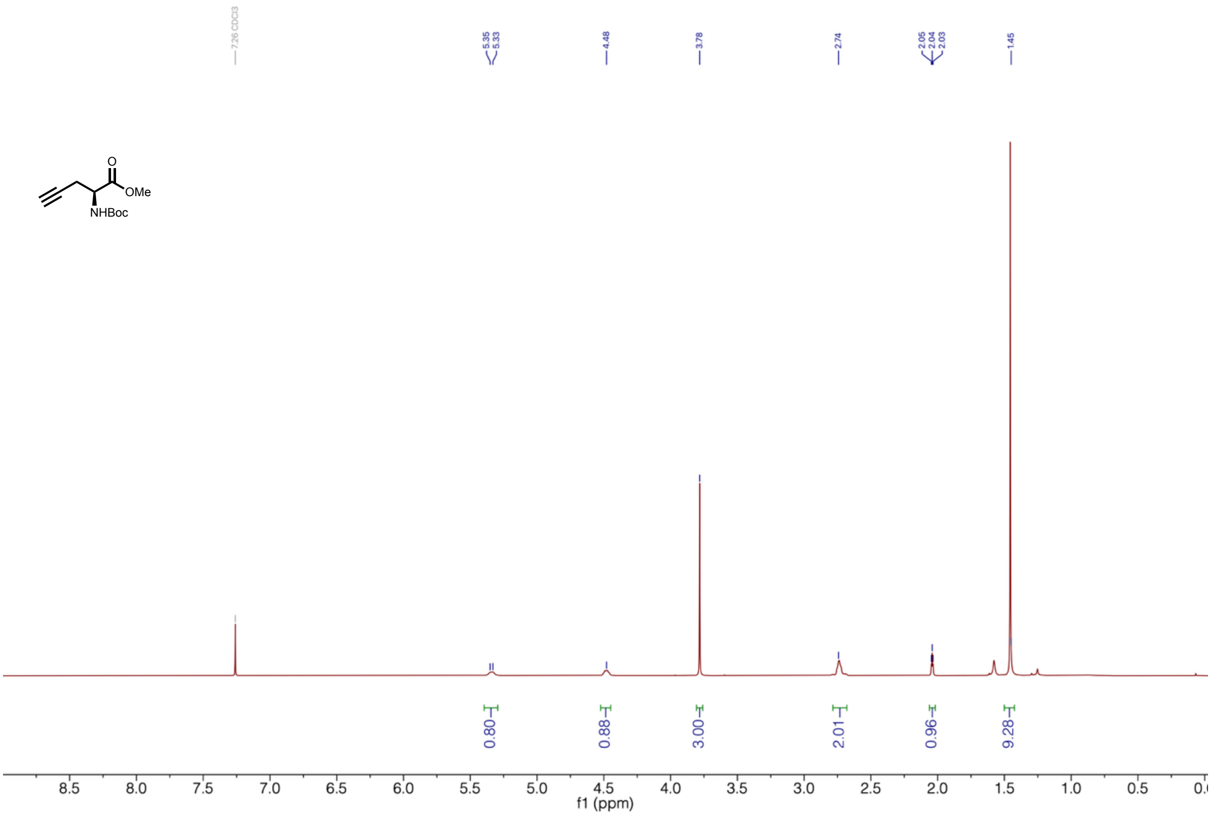


**Supplementary Figure 25 | $^1$H-NMR of *N*-Boc-propargyl-L-glycine-OMe.**

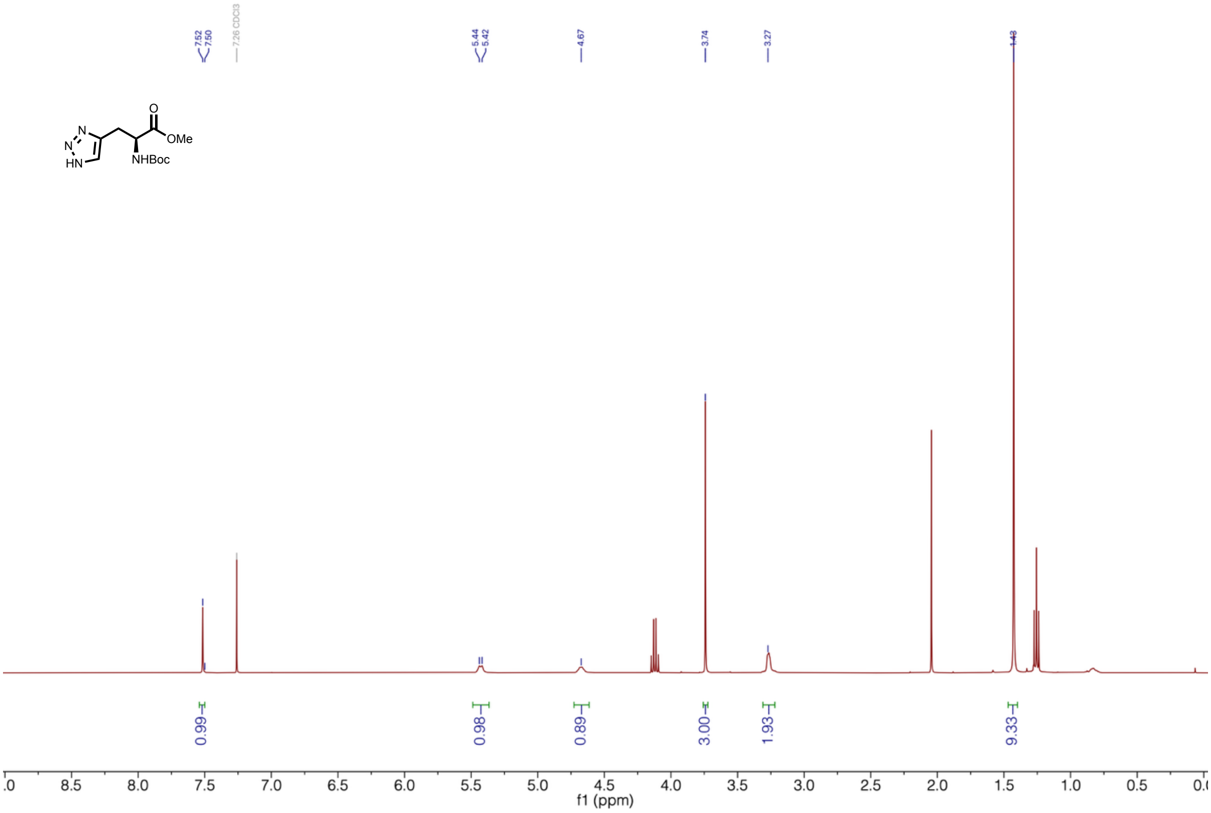


**Supplementary Figure 26 | $^{1}$H-NMR of *N*-Boc-1,2,3-triazol-4-yl-L-alanine-OMe.**

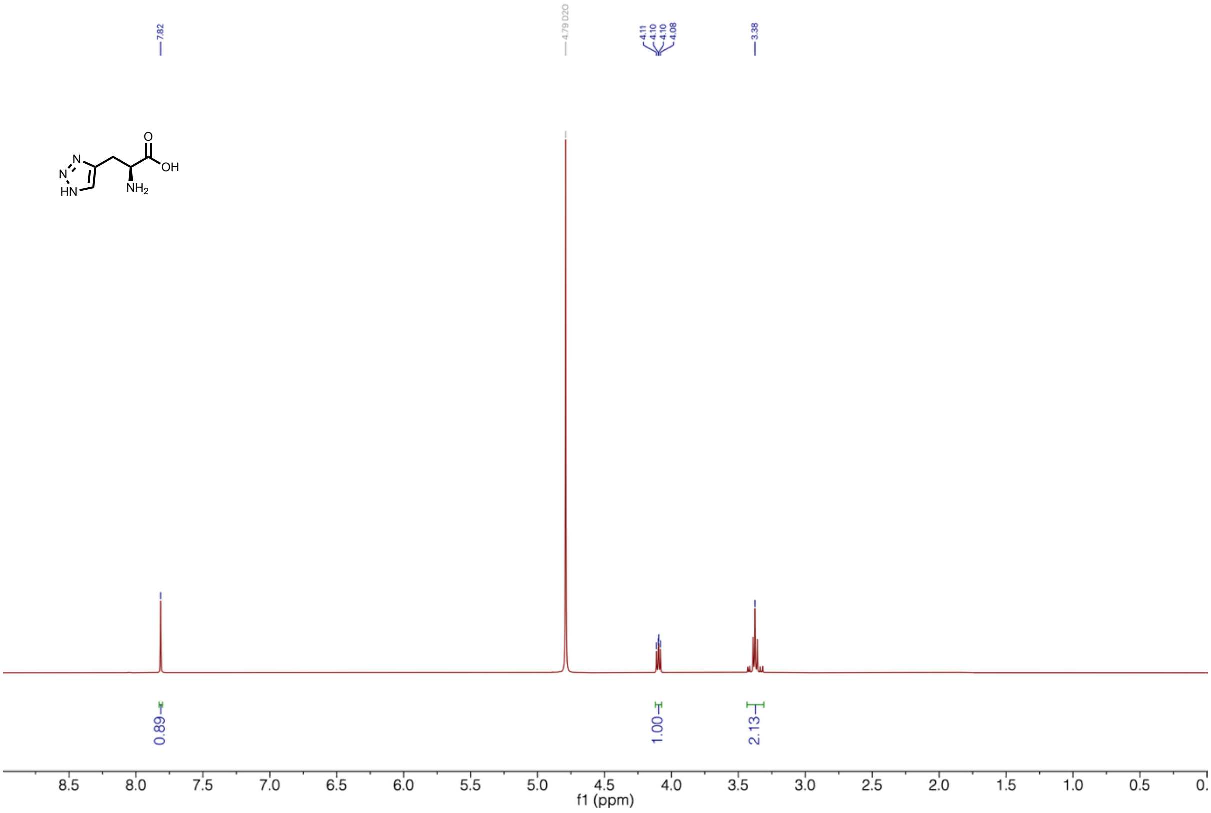


**Supplementary Figure 27 | $^{1}$H-NMR of 1,2,3-triazol-4-yl-L-alanine (123Trz-4A).**

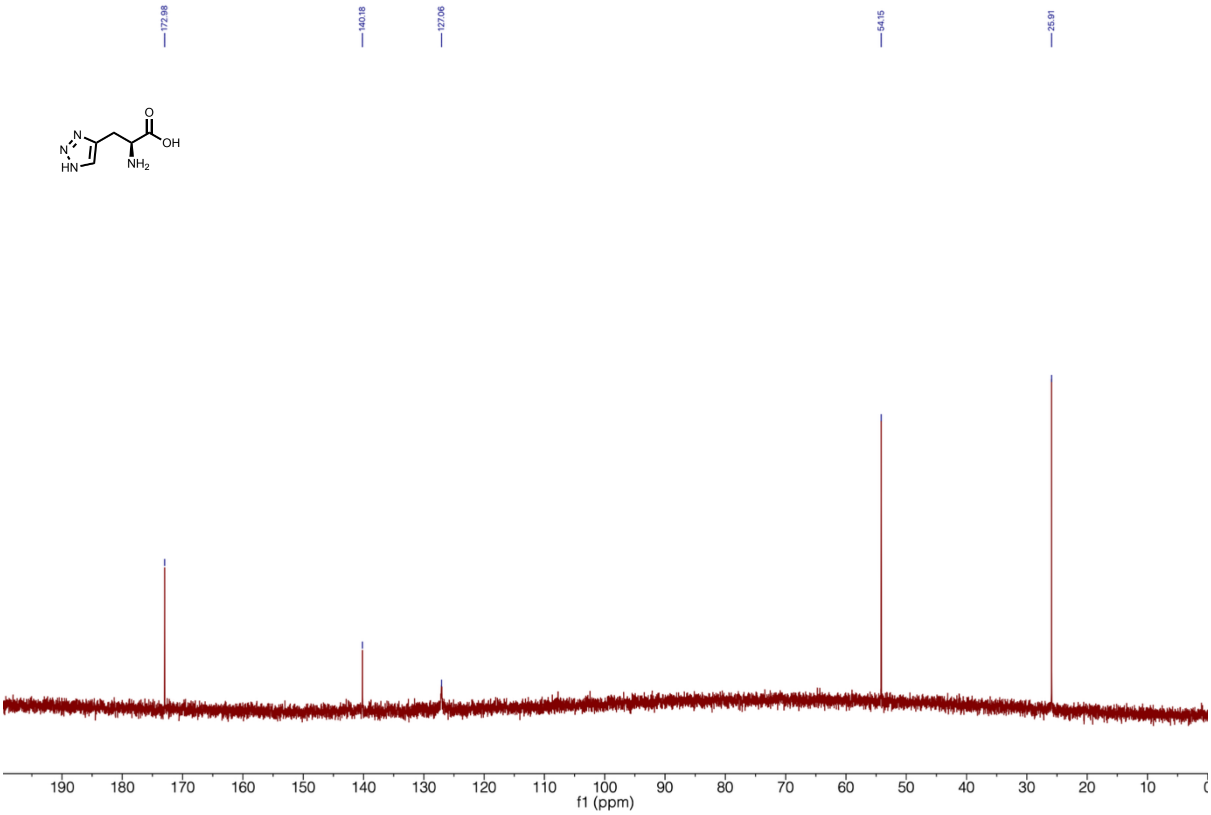


**Supplementary Figure 28 | $^{13}$C-NMR of 1,2,3-triazol-4-yl-L-alanine (123Trz-4A).**

**1-Benzyl-1,2,3-triazol-4-yl-L-alanine synthesis (1Bn123Trz-4A):**

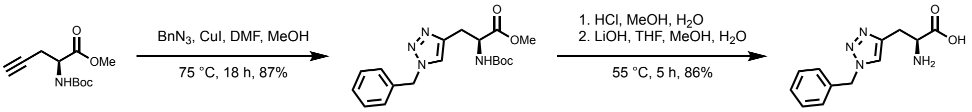


<u>*N*-Boc-1-benzyl-1,2,3-triazol-4-yl-L-alanine-OMe</u>

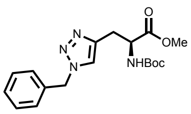

*N*-boc-propargyl-L-glycine-OMe (1.02 g, 4.48 mmol, 1.0 eq.) and benzylazide (913 mg, 6.86 mmol, 1.5 eq.) were added to a mixture of dry DMF (5 mL) and dry MeOH (1 mL) under a nitrogen atmosphere. CuI (82 mg, 430 µmol, 10 mol%) was added and a white precipitate occurred, that vanished after a short amount of time. The solution was stirred at 75 °C for 18 h. The dark green solution was cooled to 4 °C and quenched with water (25 mL). The mixture was extracted with EtOAc (2 x 125 mL), and the combined organic phases were wash with aq. LiCl (5%, 2 x 50 mL) and brine (1 x 100 mL). The organic phase was dried over $Na_2SO_4$ and the solvent was evaporated *in vacuo* to yield a green solid. The crude product was purified by automated flash column chromatography ($SiO_2$, 50-100% EtOAc in cyclohexane) to yield *N*-Boc-1-benzyl-1,2,3-triazol-4-yl-L-alanine-OMe as a green solid (1.40 g, 3.88 mmol, 87%).

Rf (EtOAc): 0.75, brown spot with ninhydrin stain.

$^1$H-NMR (400 MHz, $CDCl_3$) δ 7.41 – 7.29 (m, 3H), 7.21 (d, $^3J_{HH}$ = 7.1 Hz, 2H), 5.69 – 5.45 (m, 3H), 4.66 (s, 1H), 3.70 (s, 3H), 3.14 (br. s, 2H), 1.40 (s, 9H).

1-Benzyl-1,2,3-triazol-4-yl-L-alanine (1Bn123Trz-4A)

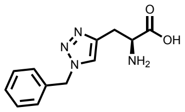


*N*-Boc-1-benzyl-1,2,3-triazol-4-yl-L-alanine-OMe (1.40 g, 3.88 mmol, 1.0 eq.) was dissolved in methanol (24 mL) and aq. HCl (conc., 1.6 mL, 19 mmol, 5.0 eq.) was added. The orange-green solution was stirred at room temperature for 5 h. The solvent was removed *in vacuo* to yield a green glue. The crude product was purified by automated flash column chromatography (C18, 5-90% acetonitrile in water (0.1% formic acid) to yield 1-benzyl-1,2,3-triazol-4-yl-L-alanine-OMe that was already partially deprotected and directly used for the next step. Crude 1-benzyl-1,2,3-triazol-4-yl-L-alanine-OMe was dissolved in THF, MeOH, and water (6 mL each). LiOH monohydrate (488 mg, 11.6 mmol, 3.0 eq.) was added and the reaction mixture was stirred at 55 °C for 30 min. The solvents were removed *in vacuo* to yield 1-benzyl-1,2,3-triazol-4-L-alanine (821 mg, 3.33 mmol, 86%) as a pale blue solid.

$^1$H-NMR (400 MHz, MeOD) δ 8.05 (s, 1H), 7.43 – 7.31 (m, 5H), 5.65 (s, 2H), 4.37 (dd, $^3J_{HH}$ = 5.7 Hz, 1H), 3.43 (dd, $^2J_{HH}$ = 15.7, $^3J_{HH}$ = 5.7 Hz, 1H), 3.40 – 3.30 (m, 1H).

$^{13}$C-NMR (101 MHz, MeOD) δ 170.5, 136.2, 130.1, 129.8, 129.4, 126.0, 55.5, 53.3, 26.9.

MS (ESI): calc. for $[M+H]^+$: 247.11, obs.: 247.1.

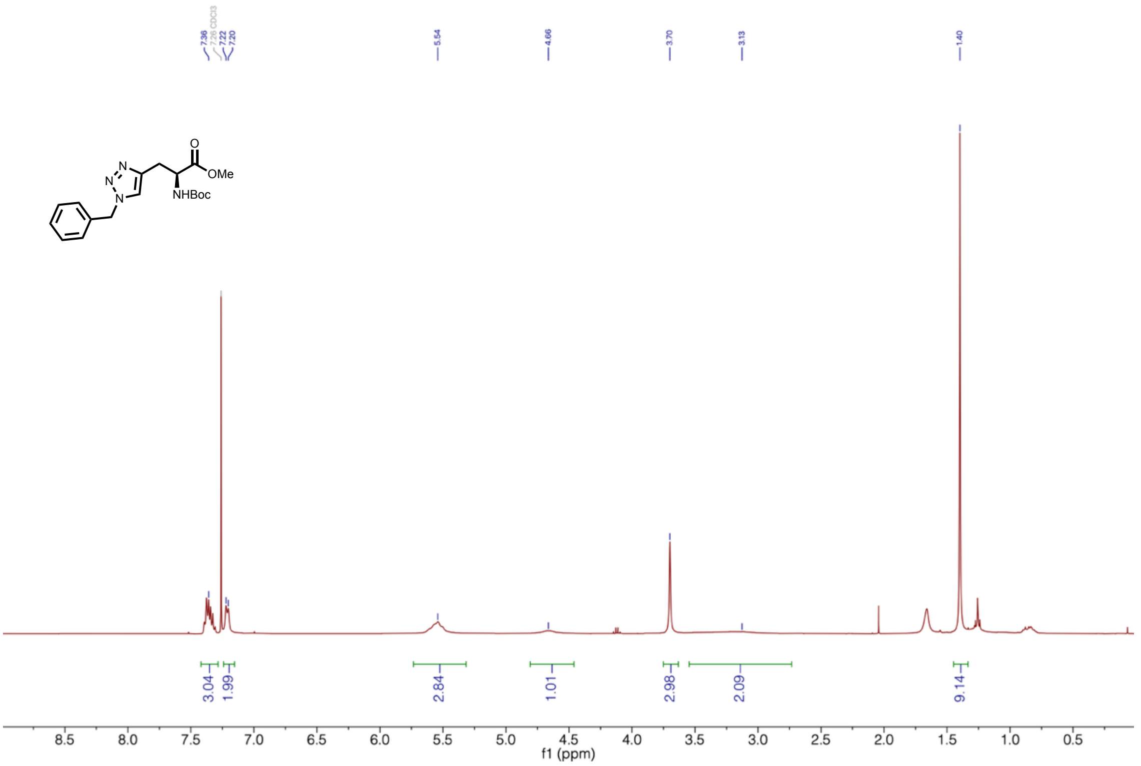


**Supplementary Figure 29 | $^{1}$H-NMR of *N*-Boc-1-benzyl-1,2,3-triazol-4-yl-L-alanine-OMe.**

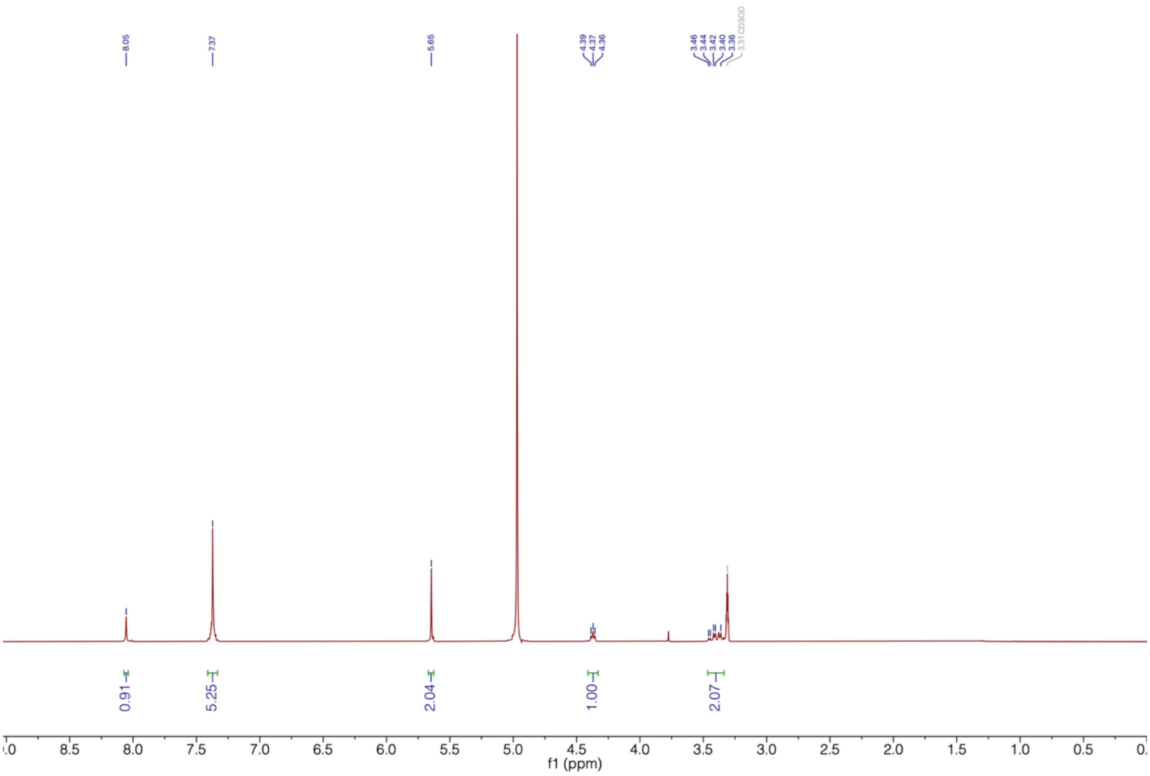


**Supplementary Figure 30 | $^{1}$H-NMR of 1-benzyl-1,2,3-triazol-4-yl-L-alanine (1Bn123Trz-4A).**

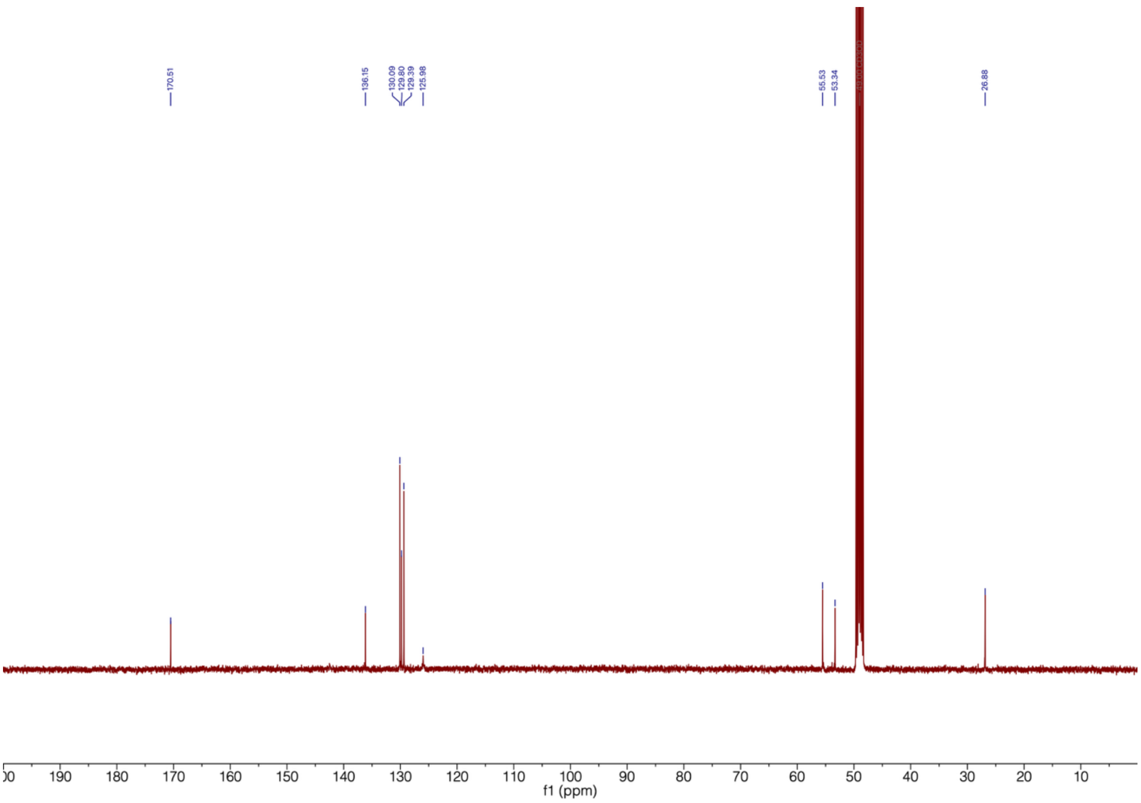


**Supplementary Figure 31 | $^{13}$C-NMR of 1-benzyl-1,2,3-triazol-4-yl-L-alanine (1Bn123Trz-4A).**

**Pyrazol-3-yl-alanine hydrochloride synthesis (3PyzA):**

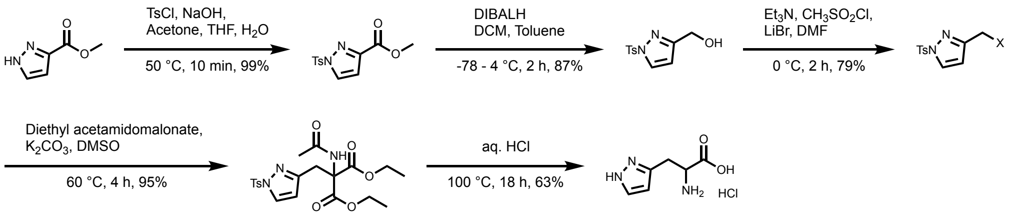


Methyl 1-tosyl-pyrazole-3-carboxylate

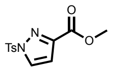

The synthesis of methyl 1-tosyl-pyrazole-3-carboxylate was adapted from literature[66]. *p*-Toluenesulfonylchloride (831 mg, 4.36 mmol, 1.1 eq.) was dissolved in acetone (6 mL). Aq. NaOH (2.5 M, 1.75 mL, 4.36 mmol, 1.1 eq.), tetrahydrofuran (1.5 mL) and methyl pyrazole-3-carboxylate (500 mg, 3.96 mmol, 1.0 eq.) were added. The reaction was stirred at 50 °C for 10 min in a microwave reactor. The suspension was diluted with dichloromethane (40 mL). The mixture was washed with water, aq. sat. $NaHCO_3$, aq. HCl (0.5 M), and brine. The organic phase was dried over $MgSO_4$ and the solvent was removed *in vacuo* to yield methyl 1-tosyl-pyrazole-3-carboxylate (1.10 g, 3.92 mmol, 99%) as a white solid.

$^1$H NMR (400 MHz, $CDCl_3$) δ 8.13 (d, $^3J_{HH}$ = 2.8 Hz, 1H), 7.95 (d, $^3J_{HH}$ = 8.2 Hz, 2H), 7.35 (d, $^3J_{HH}$ = 8.2 Hz, 2H), 6.87 (d, $^3J_{HH}$ = 2.8 Hz, 1H), 3.90 (s, 3H), 2.43 (s, 3H).

<u>(1-tosyl-pyrazol-3-yl)methanol</u>

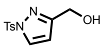


Methyl 1-tosyl-pyrazole-3-carboxylate (7.94 g, 28.3 mmol, 1.0 eq.) was dissolved in dry dichloromethane under a nitrogen atmosphere and the solution was cooled to -78 °C. DIBALH (25% in toluene, 30.1 mL, 65.0 mmol, 2.3 eq.) was added dropwise. The reaction stirred for 2 h while allowing to reach 4 °C. The reaction was quenched with water (2.7 mL), aq. NaOH (10%, 4 mL), and water (6.6 mL). The suspension was stirred for 15 min and filtered over celite. The filter cake was washed extensively with dichloromethane and the filtrate was washed with brine. The organic phase was dried over $MgSO_4$ and the solvent was removed *in vacuo* to yield (1-tosyl-pyrazol-3-yl)methanol (6.24 g, 24.7 mmol, 87%) as a white solid.

$^1$H NMR (400 MHz, $CDCl_3$) δ 8.06 (d, $^3J_{HH}$ = 2.8 Hz, 1H), 7.87 (d, $^3J_{HH}$ = 8.2 Hz, 2H), 7.32 (d, $^3J_{HH}$ = 8.2 Hz, 2H), 6.41 (d, $^3J_{HH}$ = 2.8 Hz, 1H), 4.67 (s, 2H), 2.42 (s, 3H).

<u>(1-tosyl-pyrazol-3-yl)methylchloride/bromide</u>

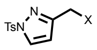


(1-tosyl-pyrazol-3-yl)methanol (12.5 g, 49.4 mmol, 1.0 eq.) was dissolved in dry DMF (100 mL) under a nitrogen atmosphere and triethylamine (9.0 mL, 65 mmol, 1.3 eq.) was added. The solution was cooled to 4 °C and methane sulfonyl chloride (5.0 mL, 64 mmol, 1.3 eq.) was added dropwise. The reaction was stirred for 45 min and lithium bromide (10.0 g, 115 mmol, 2.3 eq.) was added. The suspension was stirred for 2 h and the reaction was quenched with water (200 mL). The suspension was extracted with EtOAc (2 x 125 mL) and the combined organic phase was washed with water, aq. sat. $NaHCO_3$, aq. LiCl (5%), and brine. The organic phase was dried over $MgSO_4$ to yield a mixture of (1-tosyl-pyrazol-3-yl)methylchloride and (1-tosyl-pyrazol-3-yl)methylbromide (10 mol% (1-tosyl-pyrazol-3-yl)methylbromide, 10.8 g, 39.1 mmol, 79%) as an off-white solid.

$^1$H NMR (400 MHz, $CDCl_3$) δ 8.06 (d, $^3J_{HH}$ = 2.8 Hz, 1H, Cl), 8.05 (d, $^3J_{HH}$ = 2.8 Hz, 1H, Br), 7.89 (d, $^3J_{HH}$ = 8.2 Hz, 2H), 7.34 (d, $^3J_{HH}$ = 8.2 Hz, 1H), 6.48 (d, $^3J_{HH}$ = 2.8 Hz, 1H, Cl), 6.47 (d, $^3J_{HH}$ = 2.8 Hz, 1H, Br), 4.54 (s, 2H, Cl), 4.39 (s, 2H, Br), 2.43 (s, 3zH).

<u>Diethyl-2-acetamido-2-((1-tosyl-pyrazol-3-yl)methyl)malonate</u>

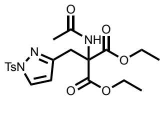


The following reaction was adeapted from literature[64]. (1-Tosyl-pyrazol-3-yl)methylchloride/bromide (39.1 mmol, 1.0 eq.) was dissolved in DMSO (75 mL). Diethyl acetamidomalonate (8.61 g, 39.6 mmol, 1.0 eq.) and $K_2CO_3$ (9.42 g, 68.2 mmol, 1.7 eq.) were added and the suspension was stirred at 60 °C for 4 h. The reaction mixture was allowed to reach room temperature and ice-cold water (500 mL) was added. The mixture was extracted with EtOAc (2 x 250 mL) and the combined organic phases were washed with Brine (250 mL). The organic phase was dried over $MgSO_4$ and the solvent was removed *in vacuo* to yield diethyl-2-acetamido-2-((1-tosyl-pyrazol-3-yl)methyl)malonate (16.8 g, 37.2 mmol, 95%) as a yellow solid.

$^{1}$H NMR (400 MHz, $CDCl_3$) δ 7.97 (d, $^{3}J_{HH}$ = 2.6 Hz, 1H), 7.82 (d, $^{3}J_{HH}$ = 8.2 Hz, 2H), 7.33 (d, $^{3}J_{HH}$ = 8.2 Hz, 2H), 6.55 (s, 1H), 6.13 (d, $^{3}J_{HH}$ = 2.6 Hz, 1H), 4.19 (q, $^{3}J_{HH}$ = 7.1 Hz, 4H), 3.66 (s, 2H), 2.42 (s, 3H), 1.89 (s, 3H), 1.23 (t, $^{3}J_{HH}$ = 7.1 Hz, 6H).

<u>Pyrazol-3-yl-alanine hydrochloride (3PyzA):</u>

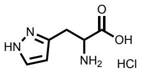


Diethyl-2-acetamido-2-((1-tosyl-pyrazol-3-yl)methyl)malonate (16.8 g, 37.2 mmol, 1.0 eq.) was dissolved in aq. HCl (4 M, 250 mL) and stirred at 100 °C for 16 h. The solvent was removed *in vacuo* and the crude product was purified by automated flash column chromatography (C18, 1% MeCN in water (0.1% formic acid)) to yield pyrazol-3-yl-alanine hydrochloride (4.50 g. 23.5 mmol, 63%) as a white solid.

$^{1}$H-NMR (400 MHz, $D_2O$) δ 7.98 (d, $^{3}J_{HH}$ = 2.6 Hz, 1H), 6.61 (d, $^{3}J_{HH}$ = 2.6 Hz, 1H), 4.42 (t, $^{3}J_{HH}$ = 6.4 Hz, 1H), 3.54 – 3.39 (m, 2H).

$^{13}$C-NMR (101 MHz, $D_2O$) δ 170.4, 142.9, 134.0, 107.1, 52.0, 26.4.

MS (ESI): calc. for $[M+H]^+$: 156.07, obs.: 156.1.

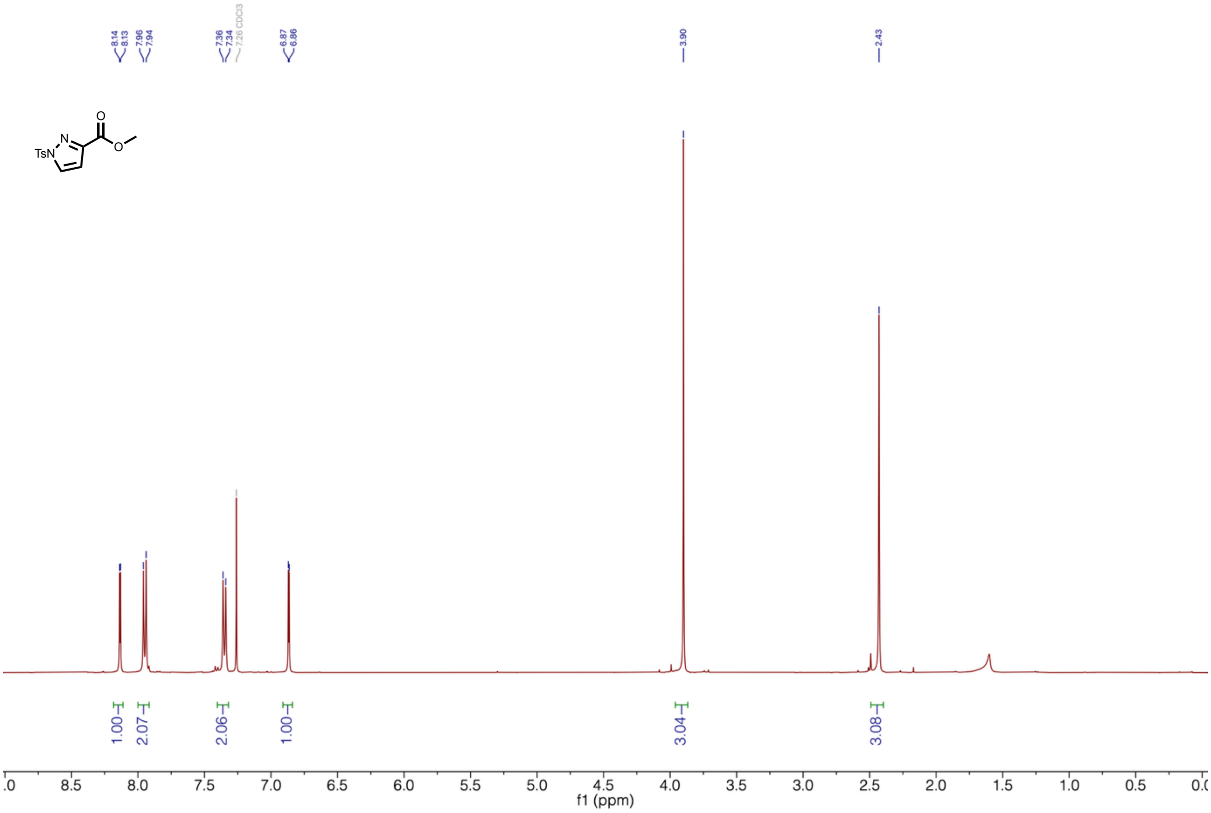

**Supplementary Figure 32 | $^{1}$H-NMR of Methyl 1-tosyl-pyrazole-3-carboxylate.**

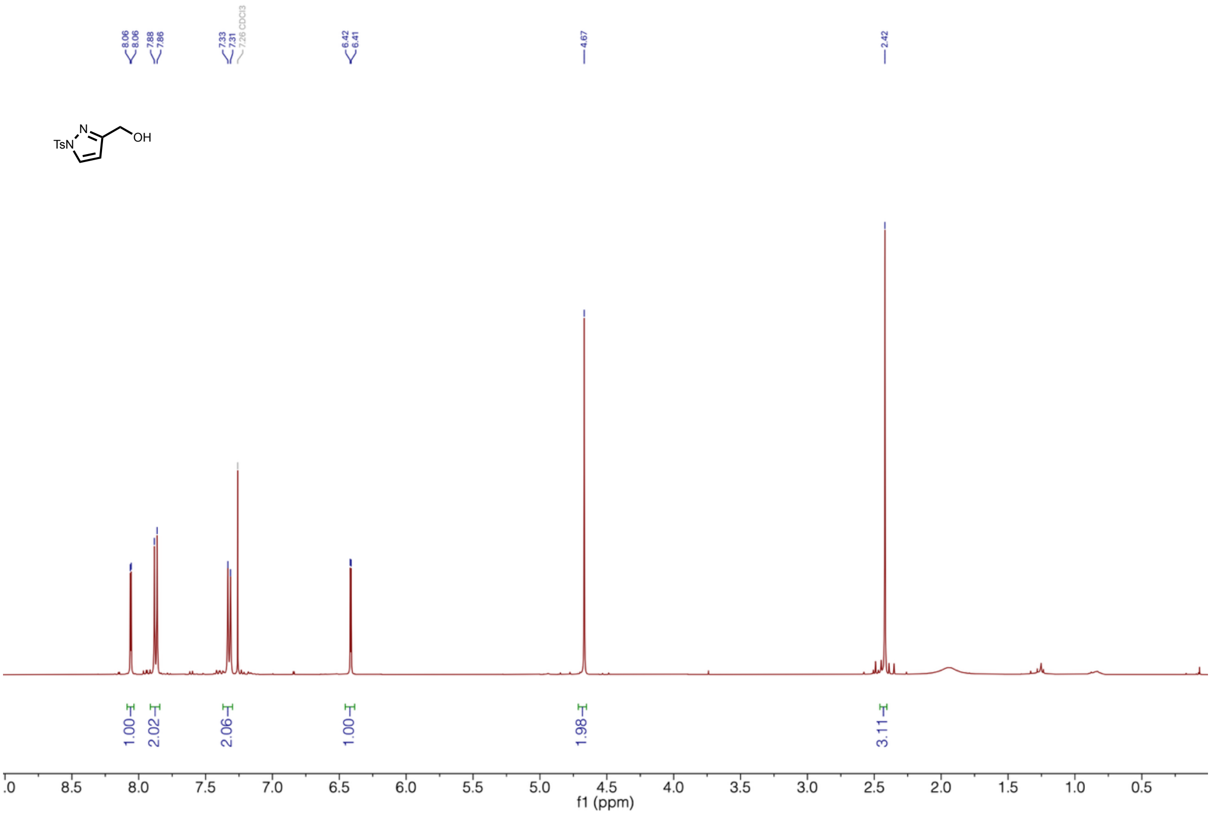


**Supplementary Figure 33 | $^{1}$H-NMR of (1-tosyl-pyrazol-3-yl)methanol.**

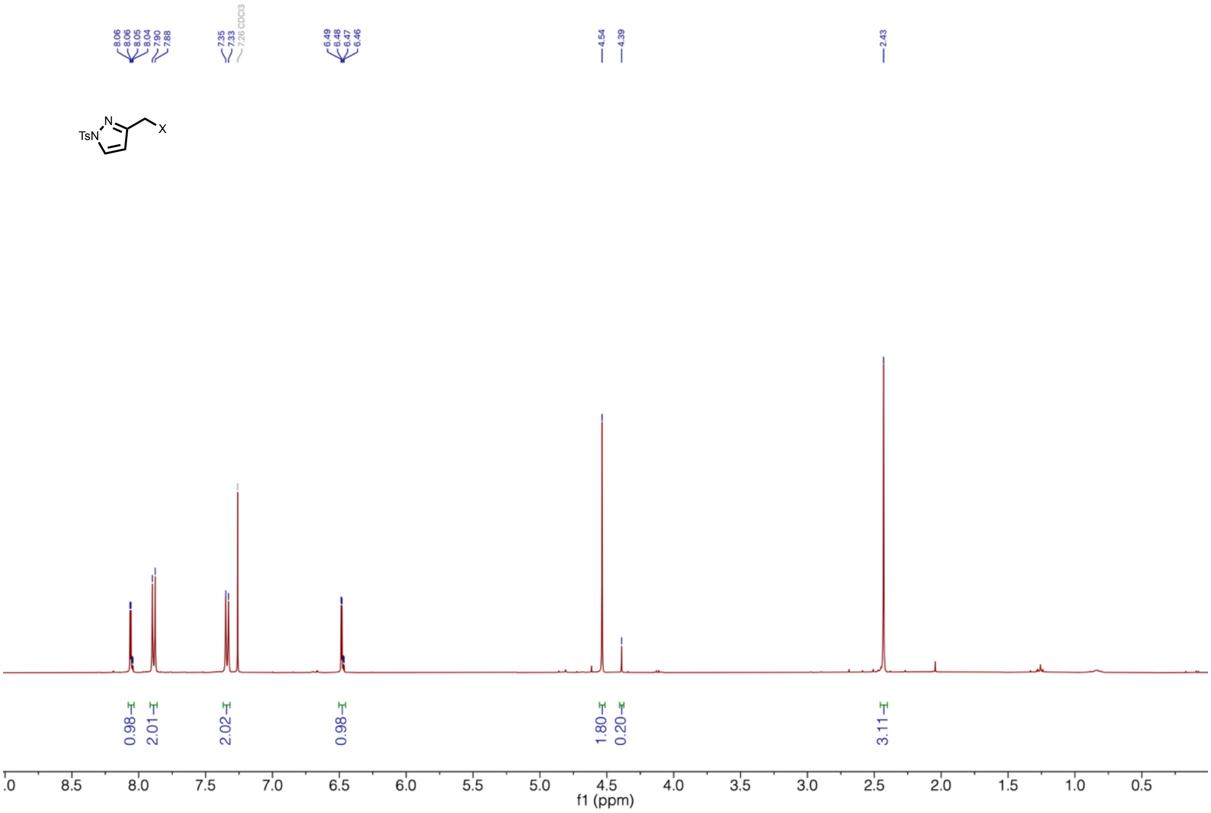

**Supplementary Figure 34 | ¹H-NMR of (1-tosyl-pyrazol-3-yl)methylchloride/bromide.**

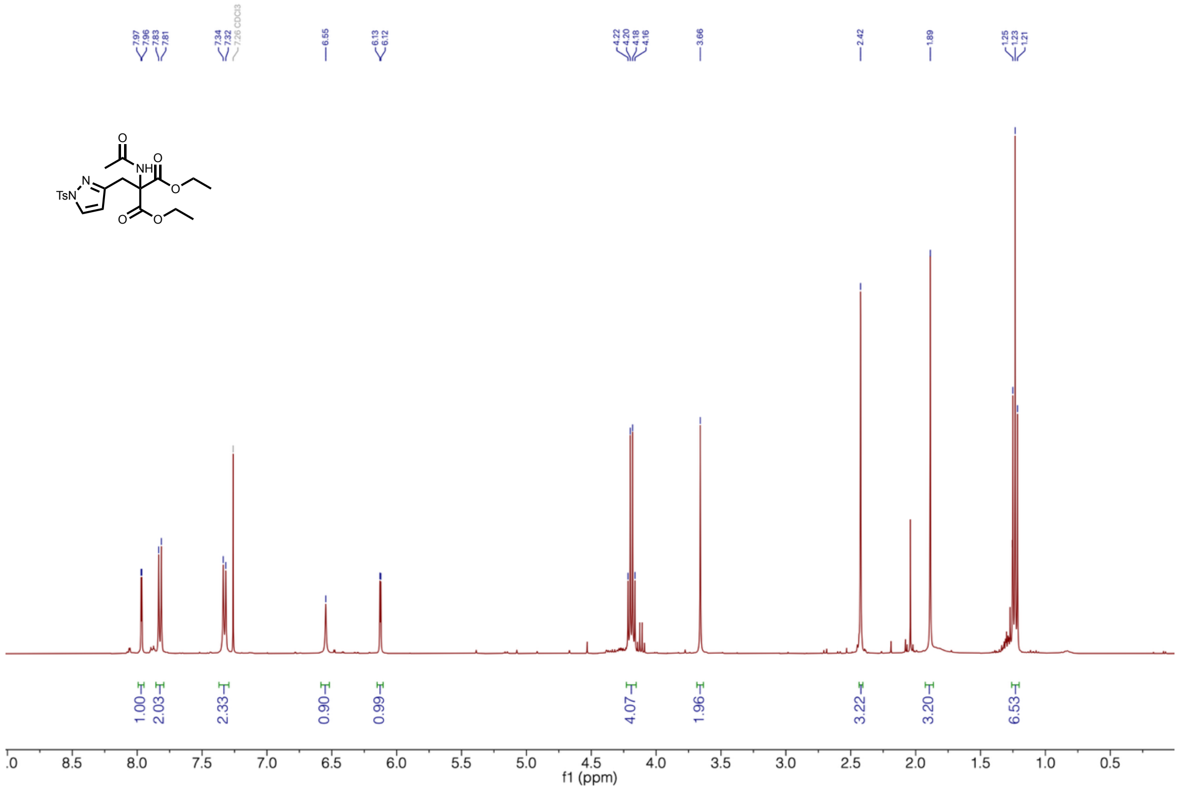


**Supplementary Figure 35 | ¹H-NMR of Diethyl-2-acetamido-2-(*N*-boc-pyrazol-3-yl-methyl)malonate.**

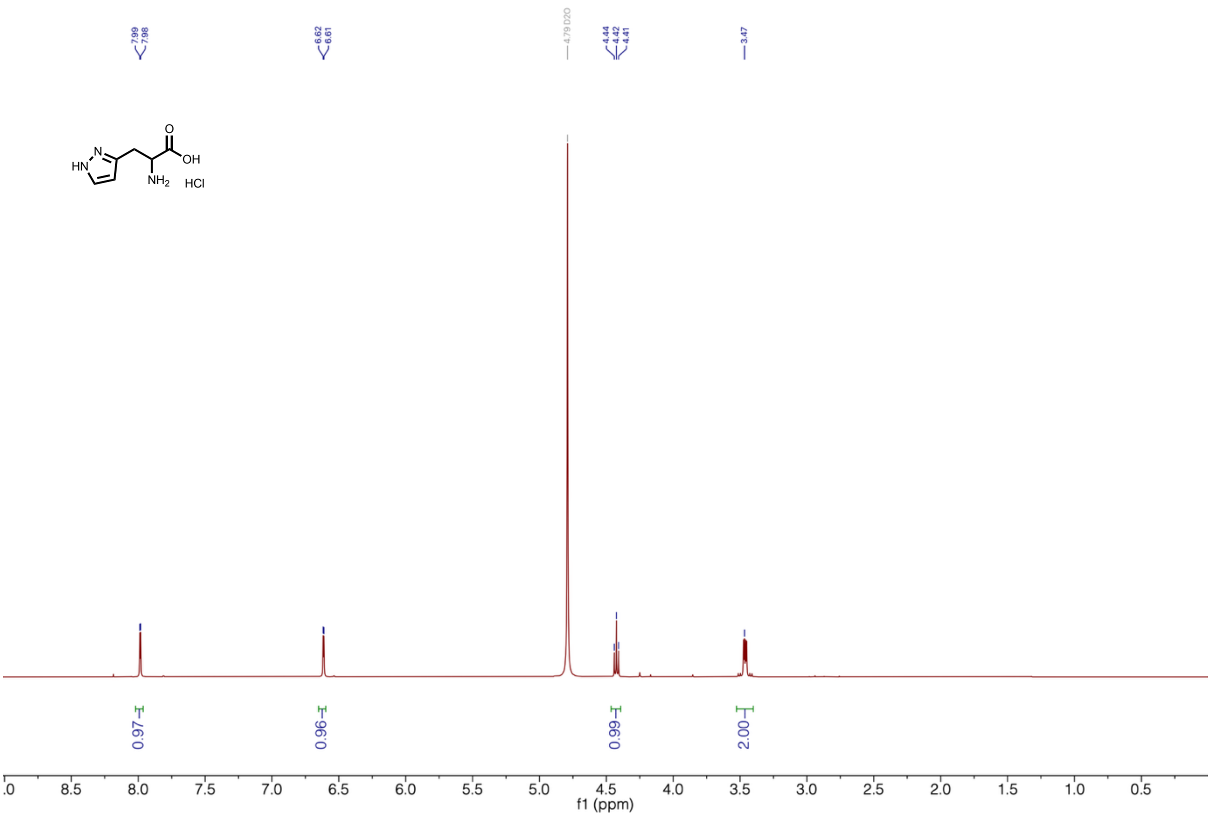

**Supplementary Figure 36 | ¹H-NMR of pyrazol-3-yl-alanine hydrochloride (3PyzA).**

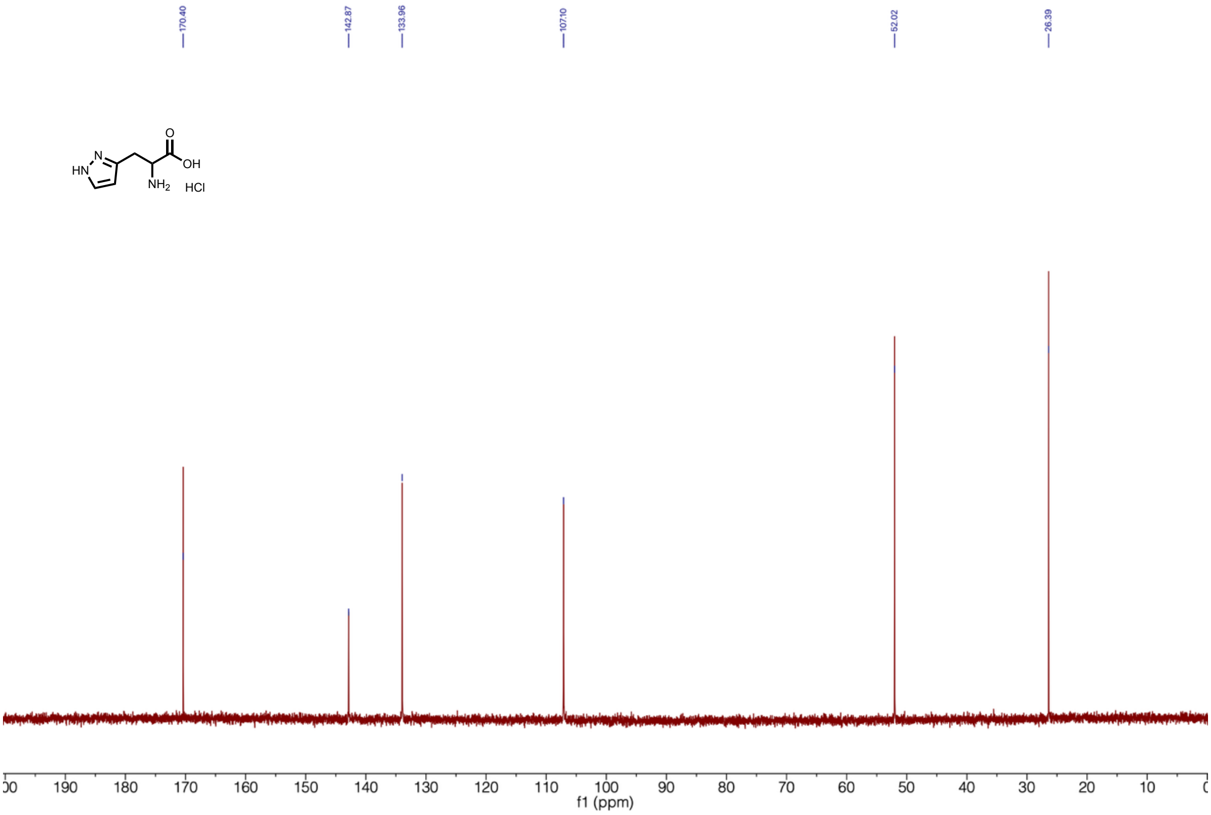


**Supplementary Figure 37 | ¹³C-NMR of pyrazol-3-yl-alanine hydrochloride (3PyzA).**

**Oxazol-4-yl-L-alanine synthesis (4OxzA):**

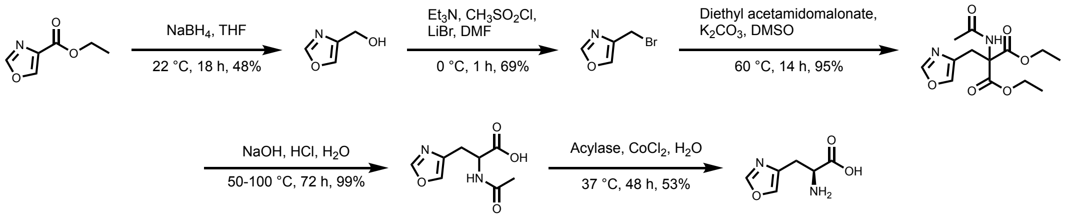


<u>(Oxazol-4-yl)methanol</u>

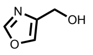

Oxazole-4-carboxylic acid ethyl ester (12.5 g, 88.6 mmol, 1.0 eq.) was dissolved in THF and water (9:1, 125 mL) and cooled to 0 °C. $NaBH_4$ (6.70 g, 177 mmol, 2.0 eq.) was added portion wise, the suspension was allowed to reach room temperature and stirred for 18 h. The suspension was cooled to 0 °C and water (10 mL) was added. The mixture was stirred for 10 min, $MgSO_4$ was added, and stirred for another 10 min. The suspension was filtered through Celite and the filter cake was washed with a lot of EtOAc. The solvent was removed *in vacuo*. The crude product was purified by automated flash column chromatography ($SiO_2$, 0 – 10% MeOH in EtOAc) to yield (oxazol-4-yl)methanol (4.18 g. 42.1 mmol, 48%) as a colorless oil.

$^1$H-NMR (400 MHz, $CDCl_3$) δ 7.89 (s, 1H), 7.64 (s, 1H), 4.64 (s, 2H).

(Oxazol-4-yl)methylbromide

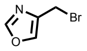


The following reaction was adapted from literature[67]. (Oxazol-4-yl)methanol (4.15 g, 41.9 mmol, 1.0 eq.) and $Et_3N$ (7.6 mL, 55 mmol, 1.3 eq.) were dissolved in dry DMF (100 mL) under a nitrogen atmosphere. The solution was cooled to 0 °C and methanesulfonyl chloride (4.2 mL, 54 mmol, 1.3 eq.) was added. The yellow suspension was stirred for 10 min and LiBr (8.37 g, 96.3 mmol, 2.3 eq.) was added. The suspension was allowed to reach room temperature and stirred for 1 h. The reaction mixture was quenched with water and extracted with EtOAc (3 x). The combined organic phase was washed with water, sat. aq. $NaHCO_3$, and brine. The organic phase was dried over $Na_2SO_4$ and the solvent was removed *in vacuo* to yield (oxazol-4-yl)methylbromide (4.69 g, 29.0 mmol, 69%) as a light-brown oil that still contained DMF and EtOAc.

$^1$H-NMR (400 MHz, $CDCl_3$) δ 7.88 (s, 1H), 7.69 (s, 1H), 4.53 (s, 2H).

Diethyl-2-acetamido-2-(oxazol-4-yl-methyl)malonate

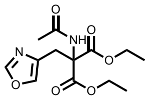


The following reaction was adeapted from literature [64]. (Oxazol-4-yl)methylbromide (7.75 g, 28.7 mmol, 1.1 eq.) and diethyl-2-acetamidomalonate (5.94 g, 27.3 mmol, 1.0 eq.) were dissolved in DMSO (40 mL) and $K_2CO_3$ (5.67, 41.0 mmol, 1.5 eq.) was added. The suspension was stirred at 60 °C for 14 h and allowed to reach room temperature. Ice-cold water (200 mL) was added and the mixture was extracted with EtOAc (3 x 100 mL). The combined orgainc phase was washed with Brine (2 x 100 mL) and dried over $Na_2SO_4$ to yield diethyl-2-acetamido-2-(oxazol-4-yl-methyl)malonate (7.78 g, 26.1 mmol, 95%) as a beige solid.

$^1$H-NMR (400 MHz, $CDCl_3$) δ 7.80 (s, 1H), 7.44 (s, 1H), 6.73 (s, 1H), 4.28 (q, $^3J_{HH}$ = 7.1 Hz, 4H), 3.62 (s, 2H), 2.00 (s, 3H), 1.28 (t, $^3J_{HH}$ = 7.1 Hz, 6H).

*N*-Acetyl-oxazol-4-yl-alanine

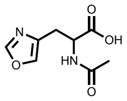


The following reaction was adapted from literature[68]. Diethyl-2-acetamido-2-(oxazol-4-yl-methyl)malonate (500 mg, 1.68 mmol, 1.0 eq.) was dissolved in water (7 mL) and NaOH (154 mg, 3.86 mmol, 2.3 eq.) was added. The solution was stirred at 50 °C for 26 h. The reaction mixture was adjusted to pH 4 with aq. HCl (6 M) and stirred at 100 °C for 39 h while adjusyting the pH every few hours to pH 4. The solvent was removed *in vacuo* to yield *N*-acetyl-oxazol-4-yl-alanine (330 mg, 1.67 mg, 99%) as a brown soldi. The crude product was used for the next step without further purification.

$^{1}$H-NMR (400 MHz, $D_2O$) δ 8.12 (s, 1H), 7.72 (s, 1H), 4.48 (dd, $^{3}J_{HH}$ = 8.7, 4.7 Hz, 1H), 3.09 (dd, $^{3}J_{HH}$ = 15.1Hz, $^{3}J_{HH}$ = 4.7 Hz, 1H), 2.94 (dd, $^{2}J_{HH}$ = 15.1Hz, $^{3}J_{HH}$ = 8.7 Hz, 1H), 2.00 (s, 3H).

Oxazol-4-yl-L-alanine (4OxzA)

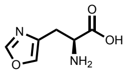


The following reaction was adapted from literature[68]. *N*-Acetyl-oxazole-alanine (7.29 g, 36.8 mmol, 1.0 eq.) was dissolved in water (100 mL) and $CoCl_2$ (13 mg, 0.27 mol%) was added. The mixture was adjusted to pH 7-8 and acylase from *Aspergillus genus* (75 mg) was added. The reaction mixture was stirred at 37 °C for 48 h and the solvent was removed *in vacuo* to yield an orange glue. The crude product was purified by preparative HPLC chromatography (HILIC, 50-90% water in acetonitrile) to yield oxazol-4-yl-L-alanine (1.51 g, 9.67 mmol, 53%) as an off-white solid.

$^{1}$H-NMR (400 MHz, $D_2O$) δ 8.15 (s, 1H), 7.80 (s, 1H), 4.03 (dd, $^{3}J_{HH}$ = 7.6, 4.9 Hz, 1H), 3.25 – 3.06 (m, 2H).

$^{13}$C-NMR (101 MHz, $D_2O$) δ 173.3, 152.9, 137.3, 133.4, 54.0, 26.6.

MS (ESI): calc. for $[M+H]^+$: 157.05, obs.: 157.1.

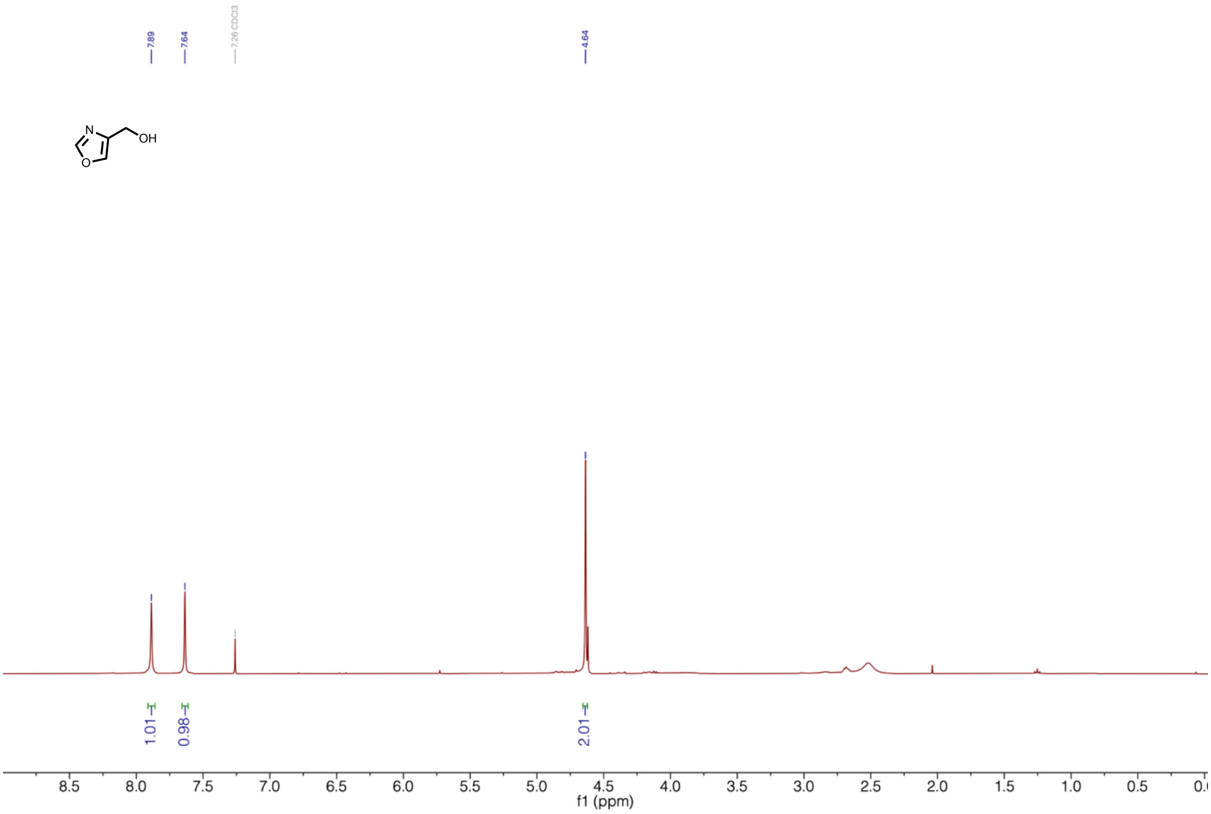


**Supplementary Figure 38 | $^{1}$H-NMR of (oxazol-4-yl)methanol.**

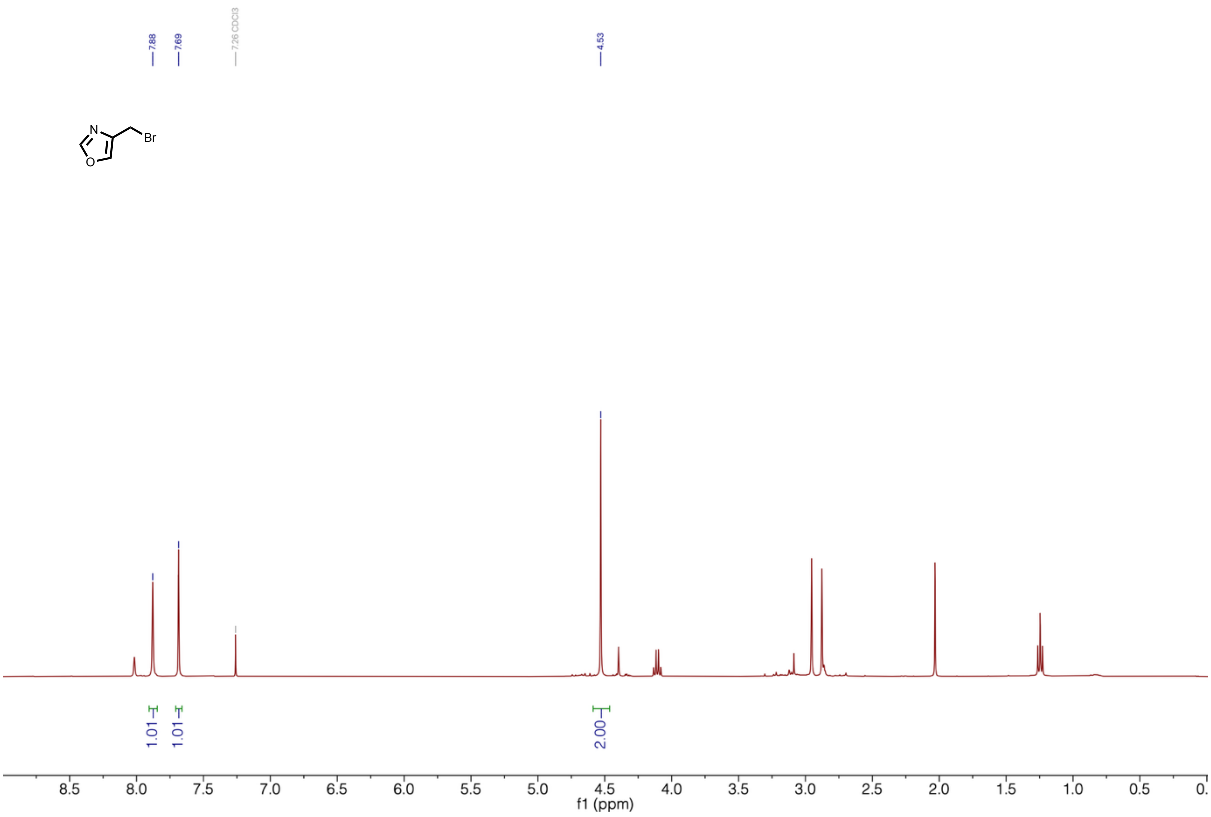

**Supplementary Figure 39 | $^{1}$H-NMR of (oxazol-4-yl)methylbromide.**

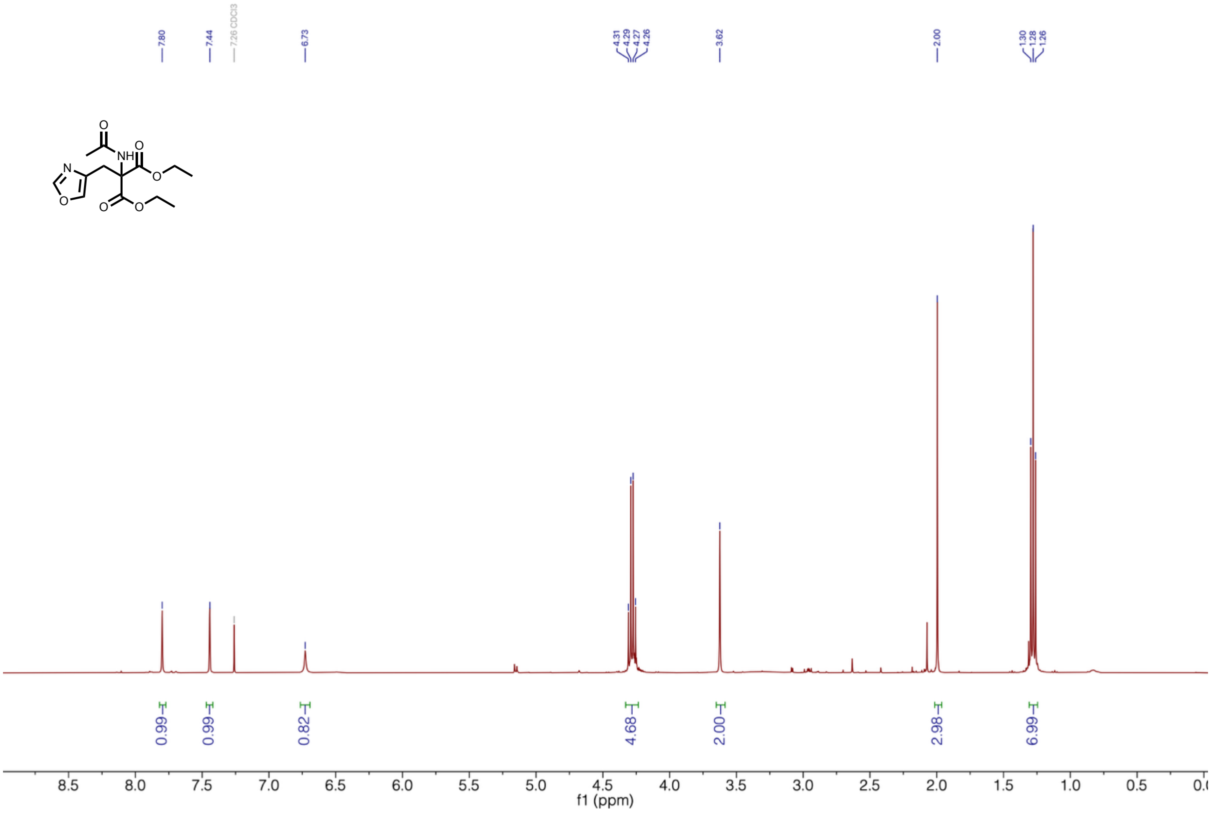

**Supplementary Figure 40 | $^{1}$H-NMR of diethyl-2-acetamido-2-(oxazol-4-yl-methyl)malonate**

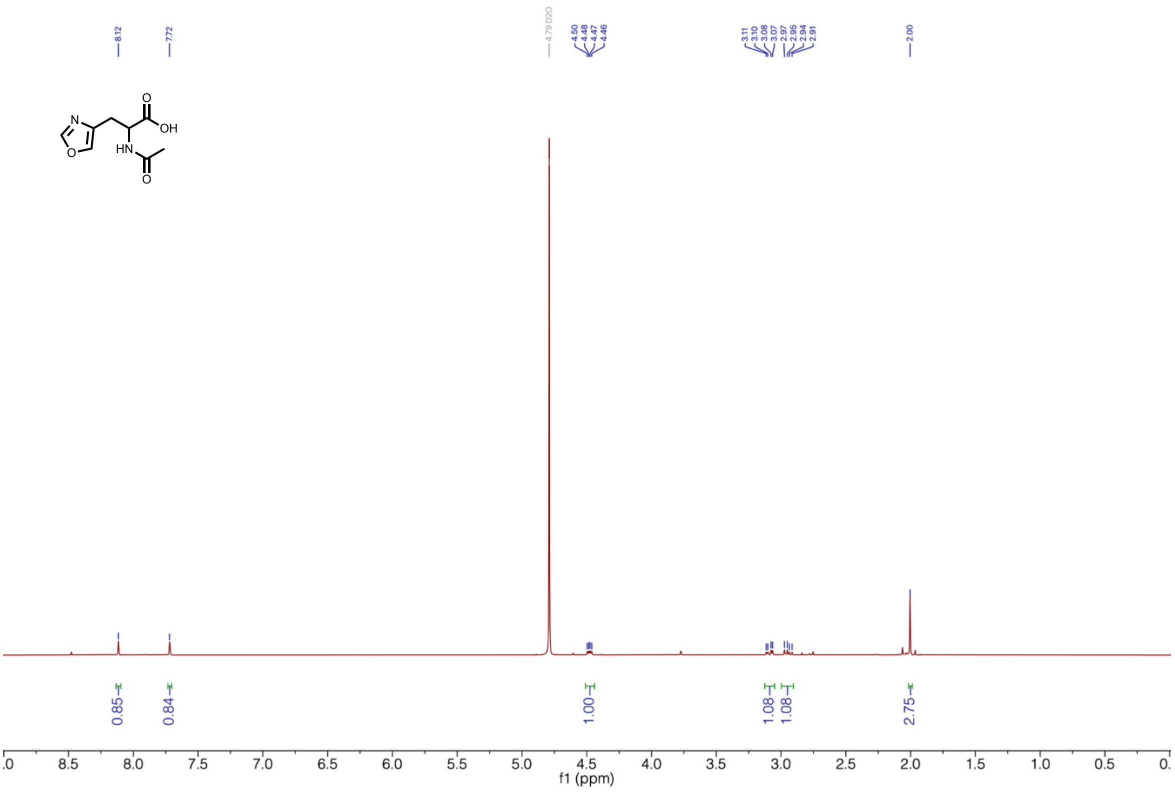


**Supplementary Figure 41 | $^{1}$H-NMR of *N*-acetyl-oxazol-4-yl-alanine.**

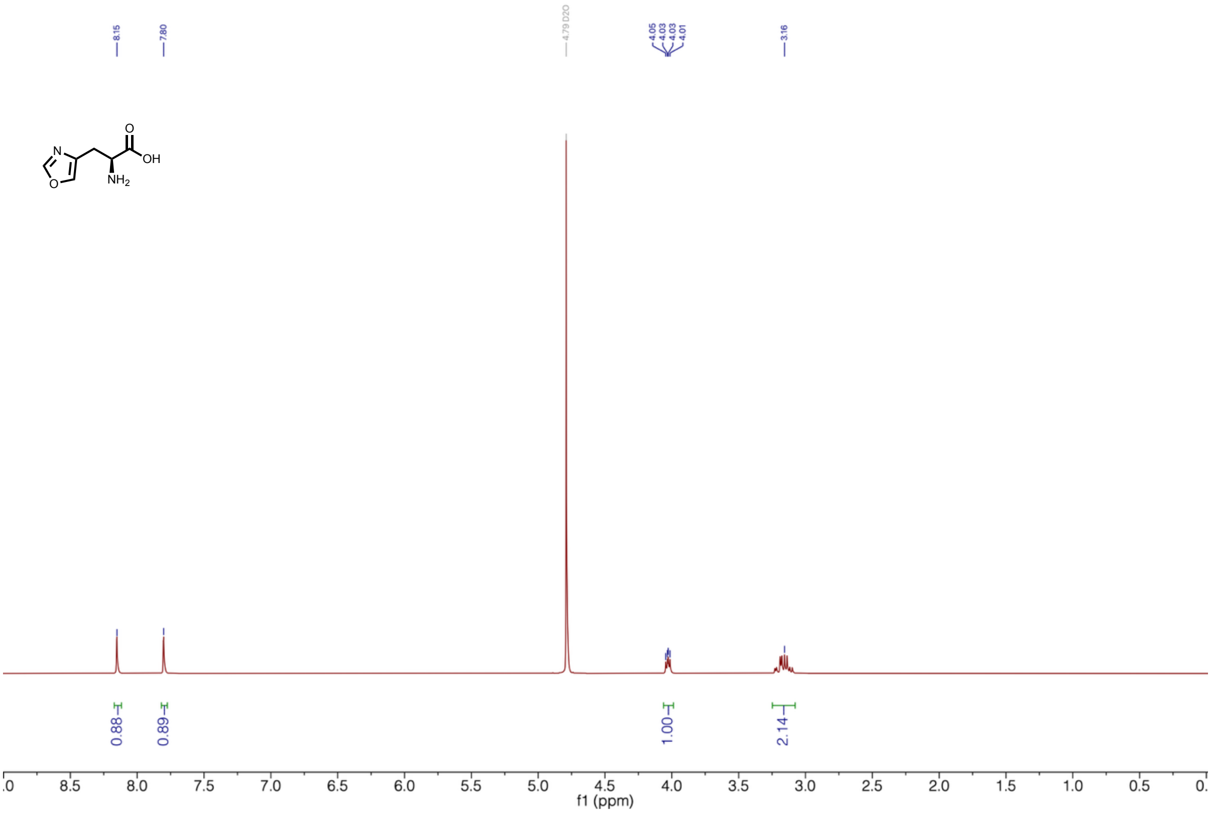


**Supplementary Figure 42 | $^{1}$H-NMR of oxazol-4-yl-L-alanine (4OxzA).**

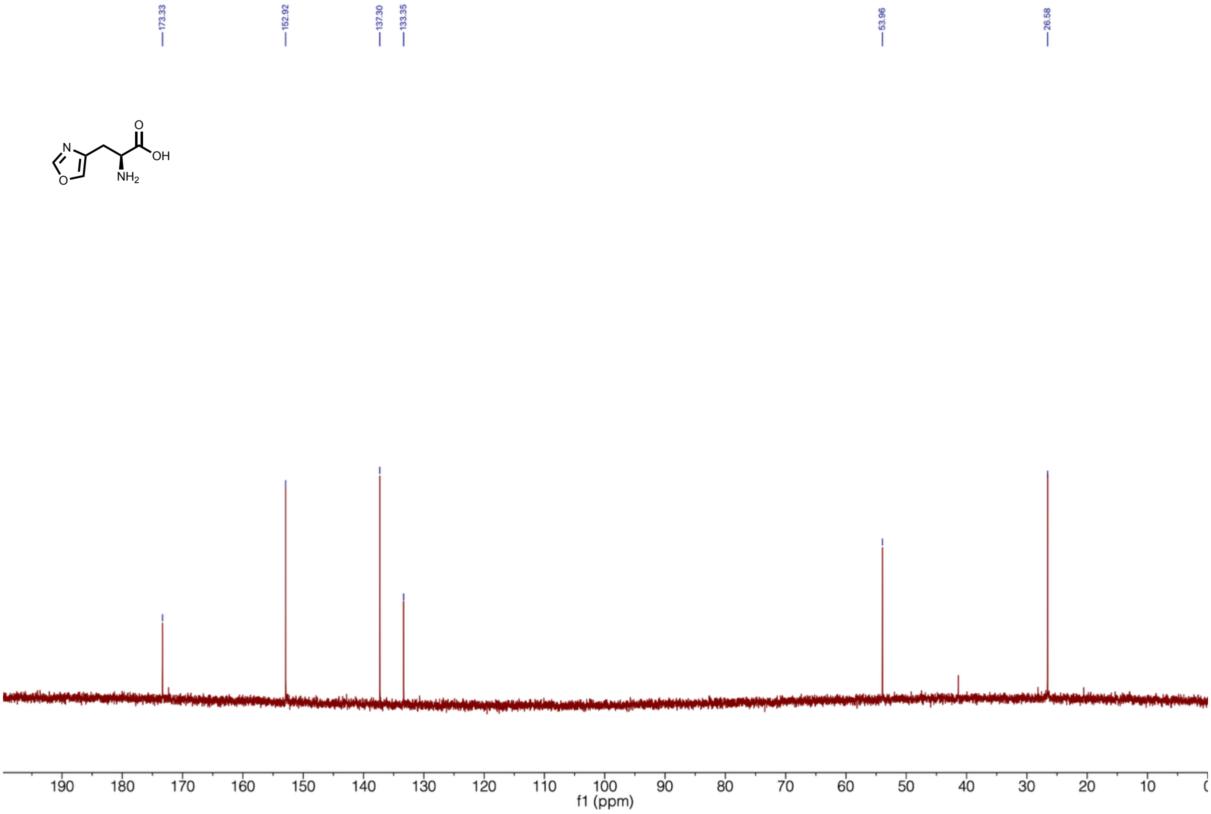


**Supplementary Figure 43 | $^{13}$C-NMR of oxazol-4-yl-L-alanine (4OxzA).**

<u>2-Ethyl-5-methyl-histidine hydrochloride synthesis (2E5MH):</u>

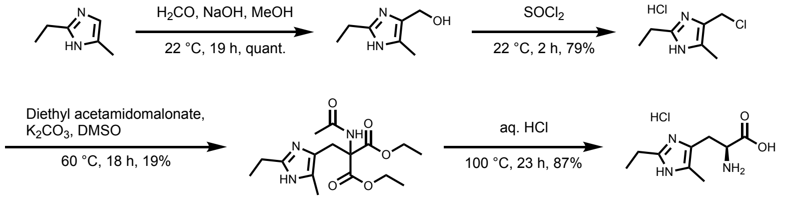


<u>2-Ethyl-4-hydroxymethyl-5-methylimidazole</u>

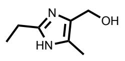

2-Ethyl-4-methylimidazole (5.00 g, 45.4 mmol, 1.0 eq.) was dissolved in 100 mL EtOH. Aq. NaOH (2.5 M, 16 mL) and paraformaldehyde (37%, 116 mmol, 2.5 eq.) was added mixture was stirred at room temperature for 19 h. The reaction was cooled to 0 °C and the pH was adjusted to pH 7 using aq. HCl (4 M). The solvent was removed *in vacuo* and the crude product was purified by automated flash column chromatography ($SiO_2$, 20 – 80% MeOH in EtOAc) to yield 2-ethyl-4-hydroxymethyl-5-methylimidazole (6363 mg, 45.4 mmol, quant.) as a yellow glue.

$^1$H-NMR (400 MHz, $D_2O$) δ 4.52 (s, 1H), 2.76 (q, $^3J_{HH}$ = 7.7 Hz, 1H), 2.19 (s, 1H), 1.25 (t, $^3J_{HH}$ = 7.7 Hz, 2H).

4-Chloromethyl-2-ethyl-5-methylimidazole hydrochloride

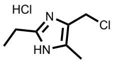


2-ethyl-4-hydroxymethyl-5-methylimidazole (5.20 g, 37.1 mmol, 1.0 eq.) was dissolved in thionyl chloride (13.6 mL, 185 mmol, 5.0 eq.) and the reaction mixture was stirred at room temperature for 2 h. Chloroform was added and the solvent was removed *in vacuo*. Chloroform was added and the solvent was removed two more times to yield 4-chloromethyl-2-ethyl-5-methylimidazole hydrochloride (5.69 g, 29.2 mmol, 79%) as a beige solid.

$^1$H-NMR (400 MHz, DMSO-d$_6$) δ 4.86 (s, 2H), 2.88 (q, $^3J_{HH}$ = 7.6 Hz, 2H), 2.27 (s, 3H), 1.29 (t, $^3J_{HH}$ = 7.6 Hz, 3H).

Diethyl-2-acetamido-2-(2-ethyl-4-methyl-imidaz-5-yl-methyl)malonate

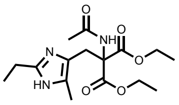


The following reaction was adapted from literature[64]. 4-Chloromethyl-2-ethyl-5-methyl-imidazole hydrochloride (5.69 g, 29.2 mmol, 1.1 eq.) was dissolved in DMSO (40 mL) and diethyl-2-acetamidomalonate (5.76 g, 26.5 mmol, 1.0 eq.) and $K_2CO_3$ (9.16 g, 66.3 mmol, 2.5 eq.) were added. The suspension was stirred at 60 °C for 18 h and the reaction mixture was allowed to reach room temperature. Ice-cold water (200 mL) was added and the solution was adjusted to pH 9-10 with aq. NaOH. The solution was extracted with EtOAc (4 x 100 mL) and the combined organic phase was washed with brine (200 mL). The organic phase was dried over $Na_2SO_4$ and the sovent was removed *in vacuo*. The crude product was purified by automated flash column chromatography ($SiO_2$, 5 – 15% MeOH in EtOAc) to yield diethyl-2-acetamido-2-(2-ethyl-4-methyl-imidaz-5-yl-methyl)malonate (1.75 g, 5.16 mmol, 19%) as a yellow glue. The product was directly used for the next step.

2-Ethyl-5-methyl-histidine hydrochloride (2E5MH)

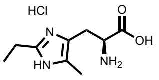


Diethyl-2-acetamido-2-(2-ethyl-4-methyl-imidaz-5-yl-methyl)malonate (1.75 g, 5.16 mmol, 1.0 eq.) was suspended in aq. HCl (4 M, 35 mL) and stirred at 100 °C for 23 h. The solvent was removed *in vacuo* and the crude product was purified by automated flash column chromatography (C18, 1% acetonitrile in water) to yield 2-ethyl-5-methyl-histidine hydrochloride (1.05 g, 4.49 mmol, 87%) as a white solid.

$^1$H-NMR (400 MHz, D$_2$O) δ 4.22 (t, $^3J_{HH}$ = 7.1 Hz, 1H), 3.37 – 3.21 (m, 1H), 2.91 (q, $^3J_{HH}$ = 7.6 Hz, 2H), 2.23 (s, 3H), 1.32 (t, $^3J_{HH}$ = 7.6 Hz, 3H).

$^{13}$C-NMR (101 MHz, D$_2$O) δ 170.8, 148.2, 127.2, 120.4, 52.3, 24.4, 18.9, 10.3, 7.9.

MS (ESI): calc. for $[M+H]^+$: 198.12, obs.: 198.2.

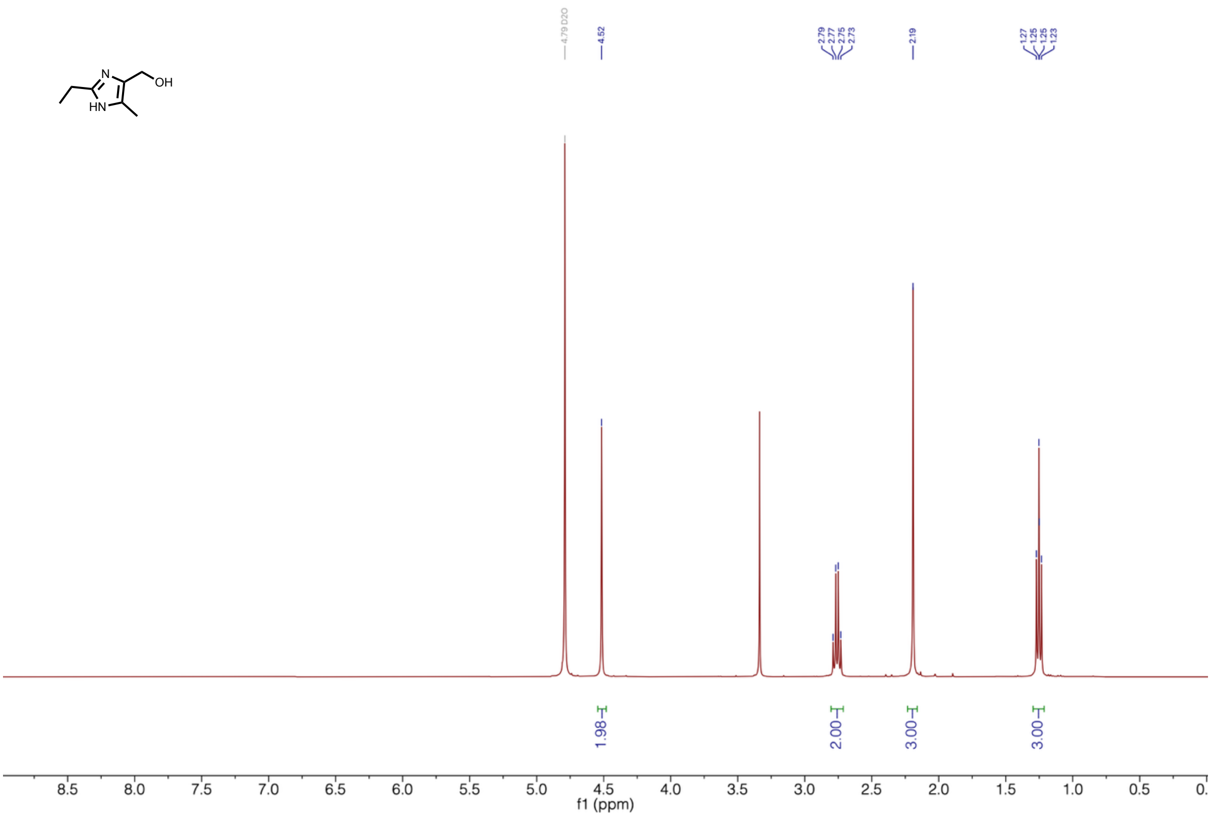


**Supplementary Figure 44 | $^{1}$H-NMR of 2-ethyl-4-hydroxymethyl-5-methylimidazole.**

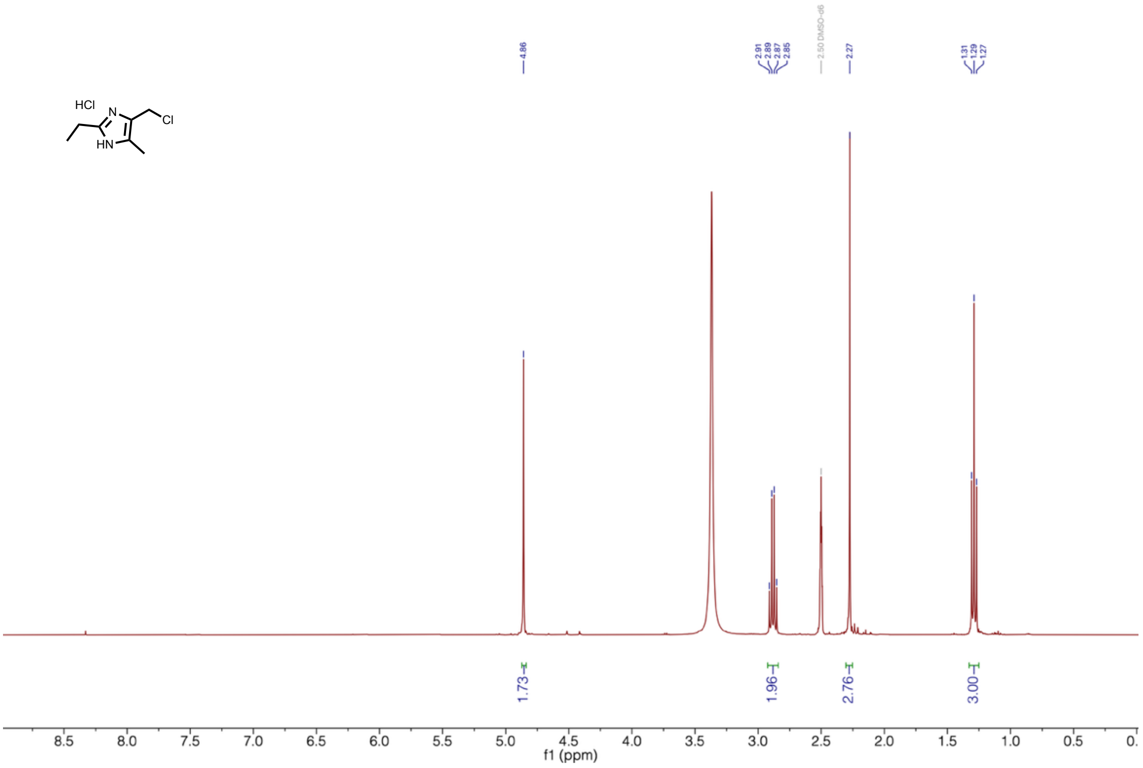


**Supplementary Figure 45 | $^{1}$H-NMR of 4-chloromethyl-2-ethyl-5-methylimidazole hydrochloride.**

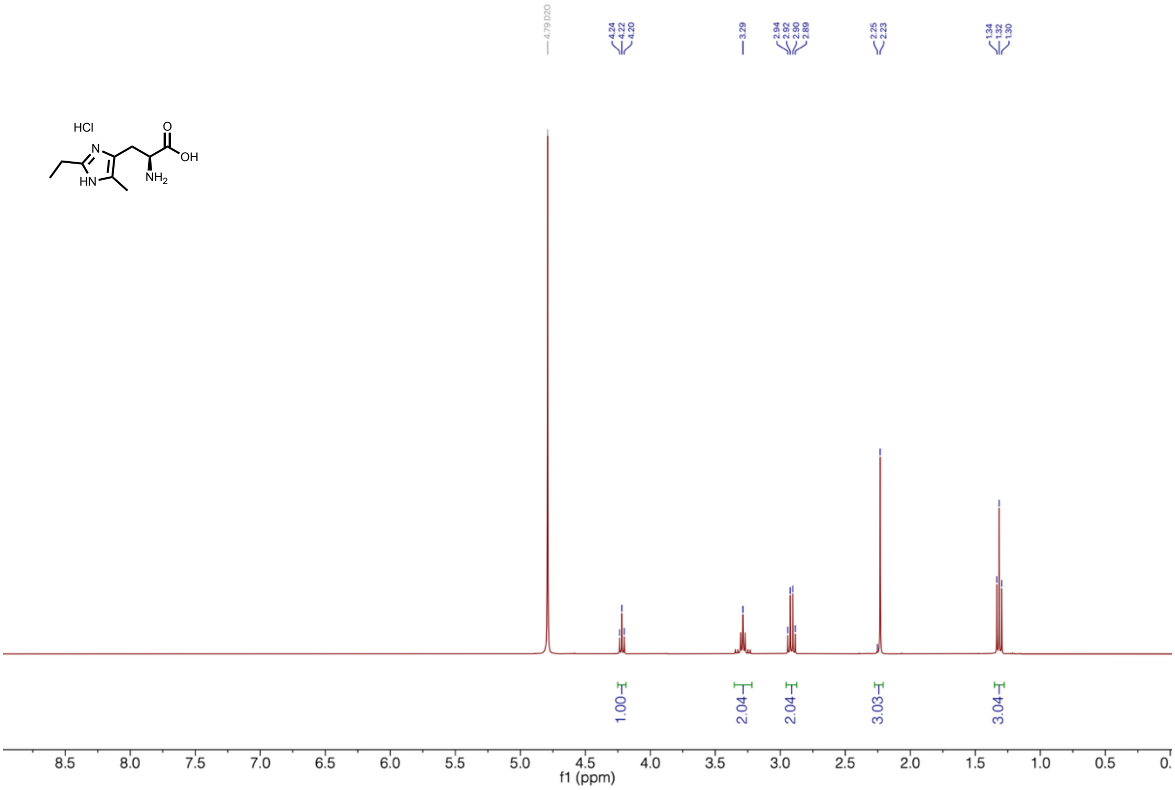


**Supplementary Figure 46 | $^{1}$H-NMR of 2-ethyl-5-methyl-histidine hydrochloride (2E5MH).**

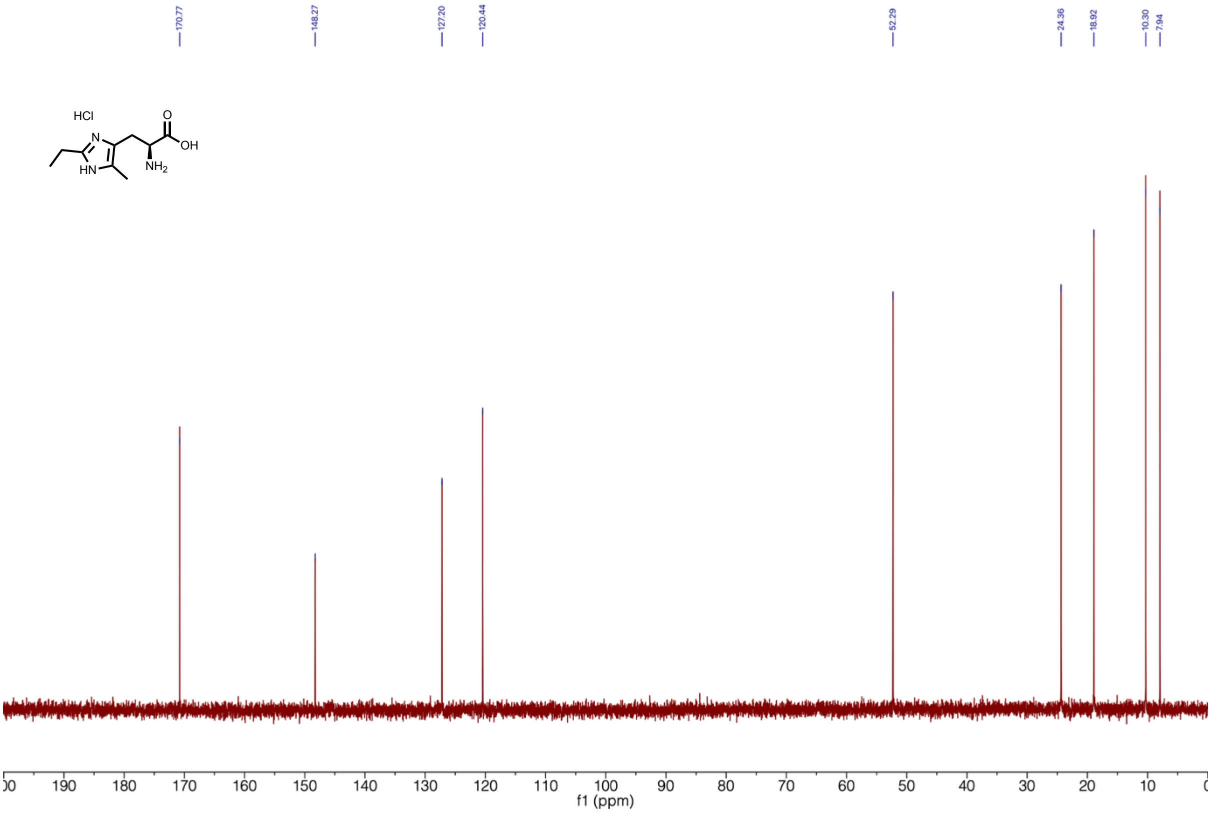


**Supplementary Figure 47 | $^{13}$C-NMR of 2-ethyl-4-methyl-histidine hydrochloride (2E5MH).**

**1,2,4-Triazol-1-yl-L-alanine synthesis (124Trz-1A):**

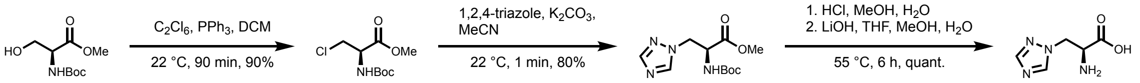


*N*-Boc-chloro-L-alanine-OMe

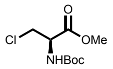

The following reaction was adapted from literature[69]. *N*-Boc-L-serine-OMe (2.00 g, 8.67 mmol, 1.0 eq.) was dissolved in dry dichloromethane (32 mL) under a nitrogen atmosphere. Triphenylphosphine (2.55 g. 9.71 mmol, 1.1 eq.) in dry dichloromethane (4 mL) and hexachloroethane (2.30 g, 9.62 mmol, 1.1 eq.) in dry dichloromethane (4 mL) were added in one portion and the reaction was stirred at room temperature for 90 min. The reaction was quenched with aq. sat. $NaHCO_3$ (20 mL) and the phases were separated. The aqueous phase was extracted with dichloromethane, the combined organic phase was washed with brine (2 x), and dried over $Na_2SO_4$. The solvent was removed *in vacuo* and the crude product was purified by flash column chromatography ($SiO_2$, dichloromethane) to yield *N*-boc-chloro-L-alanine-OMe (1.85 g, 7.78 mmol, 90%) as a white solid.

$^1$H-NMR (400 MHz, $CDCl_3$) δ 5.42 (d, $^3J_{HH}$ = 8.1 Hz, 1H), 4.76 – 4.67 (m, 1H), 3.97 (dd, $^2J_{HH}$ = 11.2 Hz, $^3J_{HH}$ = 3.1 Hz, 1H), 3.85 (dd, $^2J_{HH}$ = 11.2 Hz, $^3J_{HH}$ = 3.5 Hz, 1H), 3.81 (s, 3H), 1.46 (s, 9H).

*N*-Boc-1,2,4-triazol-1-yl-L-alanine-OMe

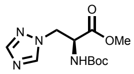

1,2,4-Triazole (1.56 g, 22.6 mmol, 3.1 eq.) was dissolved in acetonitrile and $K_2CO_3$ (1.91 g, 13.8 mmol, 1.9 eq.) was added. The suspension was stirred at room temperature for 15 min and *N*-boc-chloro-L-alanine-OMe (1.75 g, 7.36 mmol, 1.0 eq.) in acetonitrile (17 mL) was added dropwise. The suspension was stirred at room temperature for 1 min and filtered through Celite. The solvent was removed *in vacuo* and the crude product was purified by automated flash column chromatography (5 – 20% MeOH in dichloromethane) to yield *N*-boc-1,2,4-triazol-1-yl-L-alanine-OMe (1.60 g, 5.92 mmol, 80%) as a colorless oil.

$^1$H-NMR (400 MHz, $CDCl_3$) δ 8.05 (s, 1H), 7.93 (s, 1H), 5.39 (s, 1H), 4.72 – 4.59 (m, 3H), 3.79 (s, 3H), 1.44 (s, 9H).

1,2,4-Triazol-1-yl-L-alanine (124Trz-1A)

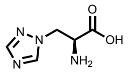

*N*-Boc-1,2,4-triazol-1-yl-L-alanine-OMe (1.60 g, 5.92 mmol, 1.0 eq.) was dissolved in MeOH and conc. aq. HCl (2.5 mL, 30 mmol, 5.0 eq.) was added. The solution was stirred at room temperature for 1 h, the solvent was removed *in vacuo* and the solid was dissolved in a mixture of THF, MeOH, and water (9 mL each). LiOH monohyderate (621 mg, 14.8 mmol, 2.5 eq.) was added and the solution was stirred at 55 °C for 6 h. The solvent was removed *in vacuo to* yield 1,2,4-trialzol-1-yl-L-alanine (924 mg, 5.92 mmol, quant.) as a white solid.

$^1$H-NMR (400 MHz, $D_2O$) δ 8.39 (s, 1H), 8.04 (s, 1H), 4.47 (dd, $^2J_{HH}$ = 14.2 Hz, $^3J_{HH}$ = 5.0 Hz, 1H), 4.41 (dd, $^2J_{HH}$ = 14.2 Hz, $^3J_{HH}$ = 6.8 Hz, 1H), 3.72 (dd, $^3J_{HH}$ = 6.8, 5.0 Hz, 1H).

$^{13}$C-NMR (101 MHz, $D_2O$) δ 168.7, 147.9, 144.0, 52.3, 49.1.

MS (ESI): calc. for $[M+H]^+$: 157.07, obs.: 157.1.

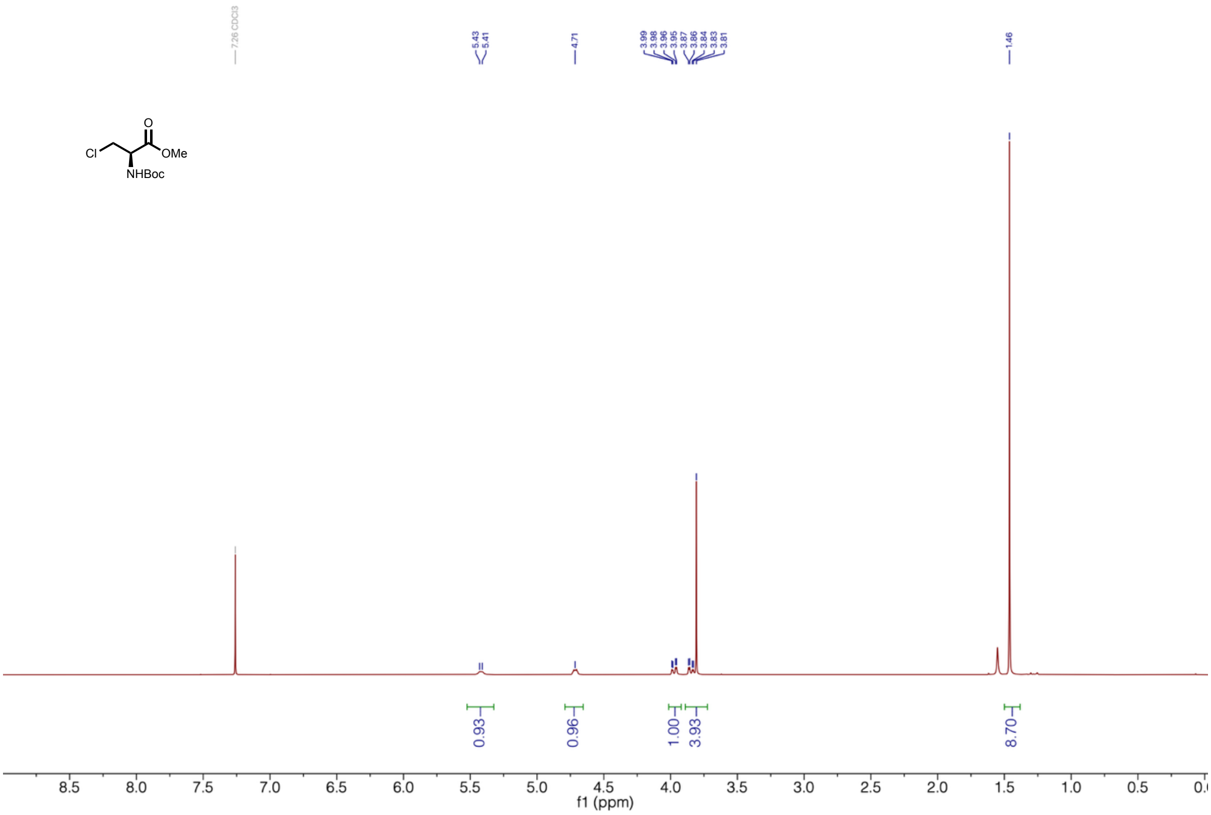


**Supplementary Figure 48 | $^1$H-NMR of *N*-Boc-chloro-L-alanine-OMe.**

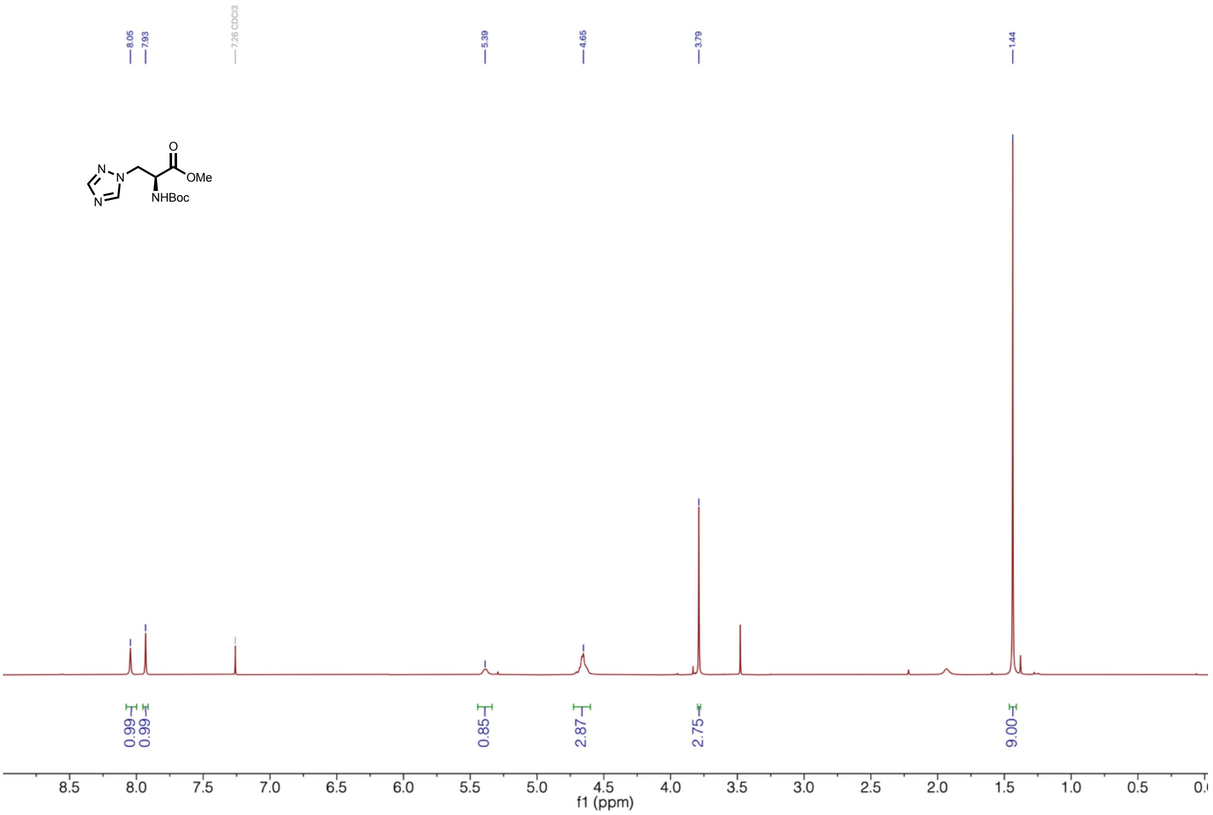


**Supplementary Figure 49 | $^{1}$H-NMR of *N*-Boc-1,2,4-triazol-1-yl-L-alanine-OMe.**

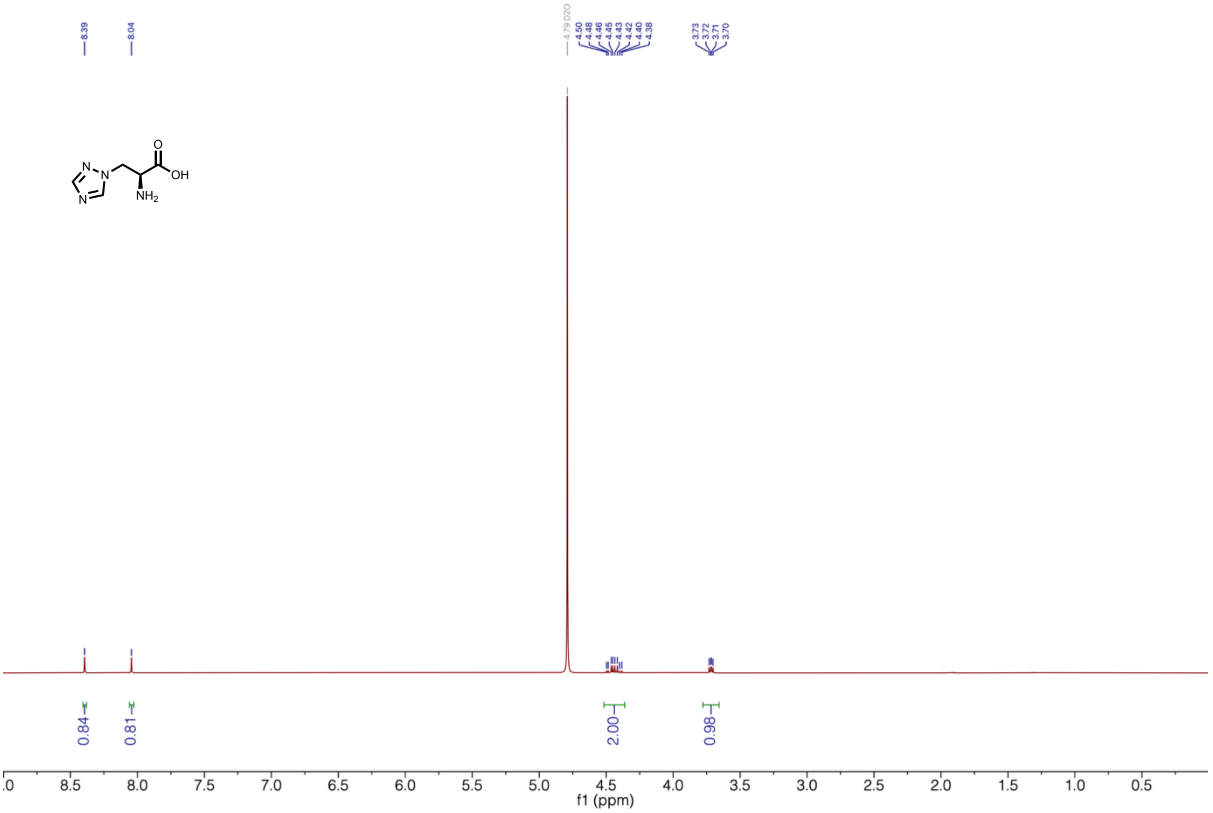


**Supplementary Figure 50 | $^{1}$H-NMR of 1,2,4-triazol-1-yl-L-alanine (124Trz-1A).**

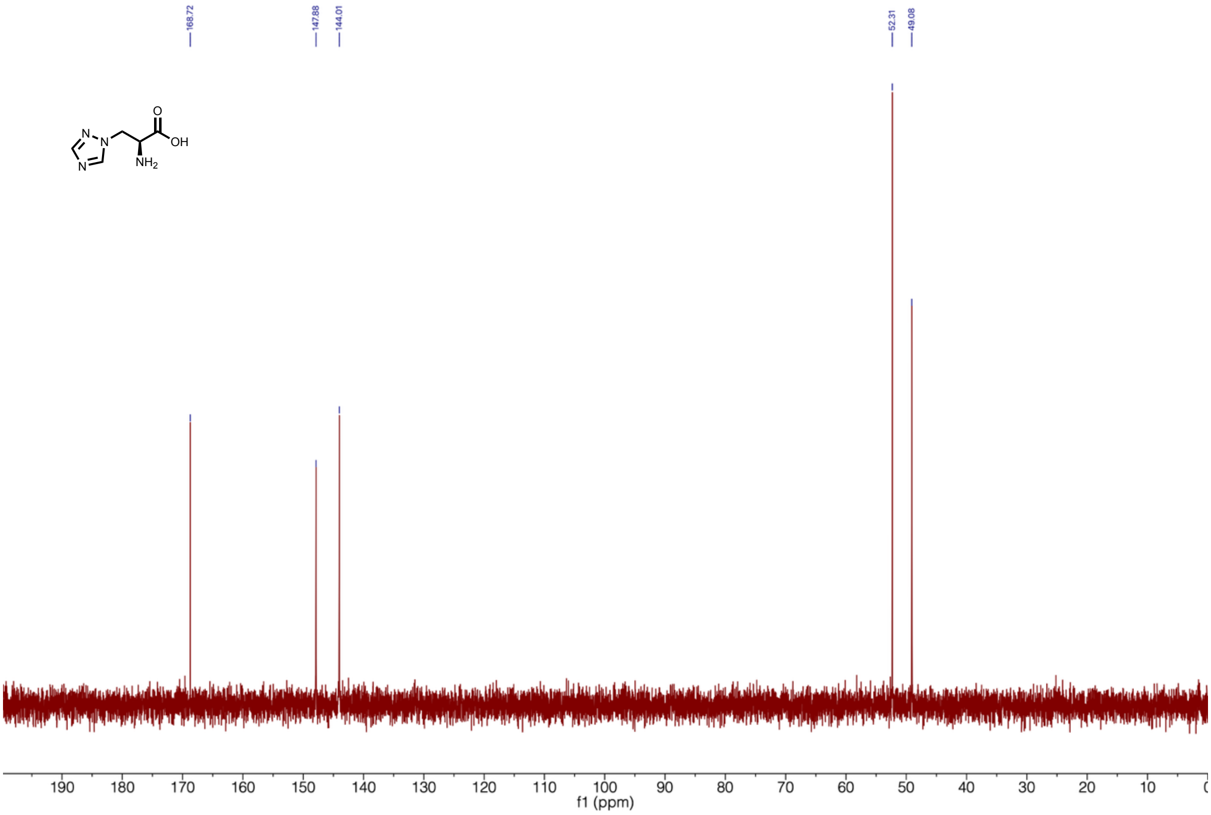


**Supplementary Figure 51 | $^{13}$C-NMR of 1,2,4-triazol-1-yl-L-alanine (124Trz-1A).**

#### **$N^{\tau}$-methyl-L-histidine synthesis (τMH)**

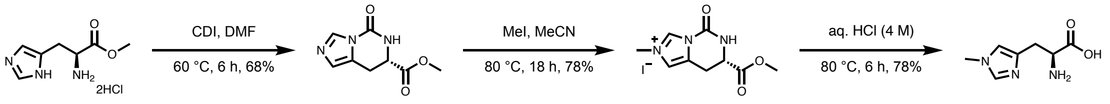


Methyl-(*S*)-5-oxo-5,6,7,8-tetrahydroimidazo[1,5-*c*]pyrimidine-7-carboxylate

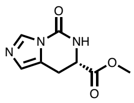

The procedure was adapted from literature[70]. L-histidine-OMe dihydrochloride (10.0 g, 41.3 mmol, 1.0 eq.) and *N*,*N*'-carbonyldiimidazole (7.37 g, 45.4 mmol, 1.1 eq.) were dissolved in DMF (200 mL). The reaction mixture was stirred at 60 ºC for 6 h. The solvent was removed *in vacuo* and the crude product was purified by automated flash column chromatography ($SiO_2$, 5% MeOH in DCM) to yield methyl-(*S*)-5-oxo-5,6,7,8-tetrahydroimidazo[1,5-c]pyrimidine-7-carboxylate (5.50 g, 28.2 mmol, 68%) as an off-white solid.

$^{1}$H-NMR (400 MHz, DMSO-$d_6$) δ 8.56-8.55 (m, 1H), 8.10 (s, 1H), 6.82 (s, 1H), 4.46-4.42 (m, 1H), 3.62 (s, 3H), 3.23-3.21 (m, 2H).

(*S*)-7-(methoxycarbonyl)-2-methyl-5-oxo-5,6,7,8-tetrahydroimidazo[1,5-*c*]pyrimidin-2-ium iodide

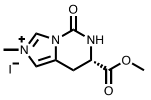

Methyl iodide (4.3 mL, 69 mmol, 6.7 eq.) was added to methyl (*S*)-5-oxo-5,6,7,8-tetrahydroimidazo[1,5-*c*]pyrimidine-7-carboxylate (2.00 g, 10.3 mmol, 1.0 eq.) in acetonitrile (60 mL). The reaction was stirred at 80 °C for 18 h. The solvent was removed *in vacuo* and the crude residue was recrystallized from 20% MeOH in DCM to yield (*S*)-7-(methoxycarbonyl)-2-methyl-5-oxo-5,6,7,8-tetrahydroimidazo[1,5-c]pyrimidin-2-ium iodide (2.70 g, 8.01 mmol, 78%) as an off-white solid.

$^{1}$H-NMR (400 MHz, $D_2O$) δ 9.38 (s, 1H), 7.42 (s, 1H), 4.74 (t, $^{3}J_{HH}$ = 5.6 Hz, 1H), 3.97 (s, 3H), 3.79 (s, 3H), 3.51-3.49 (m, 2H).

$N^{\tau}$-methyl-L-histidine

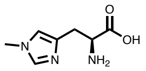

(*S*)-7-(methoxycarbonyl)-2-methyl-5-oxo-5,6,7,8-tetrahydroimidazo[1,5-c]pyrimidin-2-ium iodide (17.0 g, 50.4 mmol, 1.0 eq.) was suspended in aq. HCl (4 M, 500 mL) and stirred at 80 °C for 6 h. The solvent was removed *in vacuo* and the crude product was purified by automated flash column chromatography (C18, 0-95% acetonitrile in water (0.1% formic acid)) to yield $N^{\tau}$-methyl-L-histidine (6.40 g, 50.4 mmol, 75%) as an off-white solid.

$^{1}$H-NMR (400 MHz, $D_2O$) δ 8.64 (s, 1H), 7.38 (s, 1H), 4.19 (t, $^{3}J_{HH}$ = 6.6 Hz, 1H), 3.86 (s, 3H), 3.36-3.33 (m, 2H).

$^{13}$C-NMR (101 MHz, $D_2O$) δ 171.2, 135.5, 127.3, 121.9, 52.6, 35.6, 25.4.

MS (ESI): calc. for $[M+H]^+$: 170.09, obs.: 170.1.

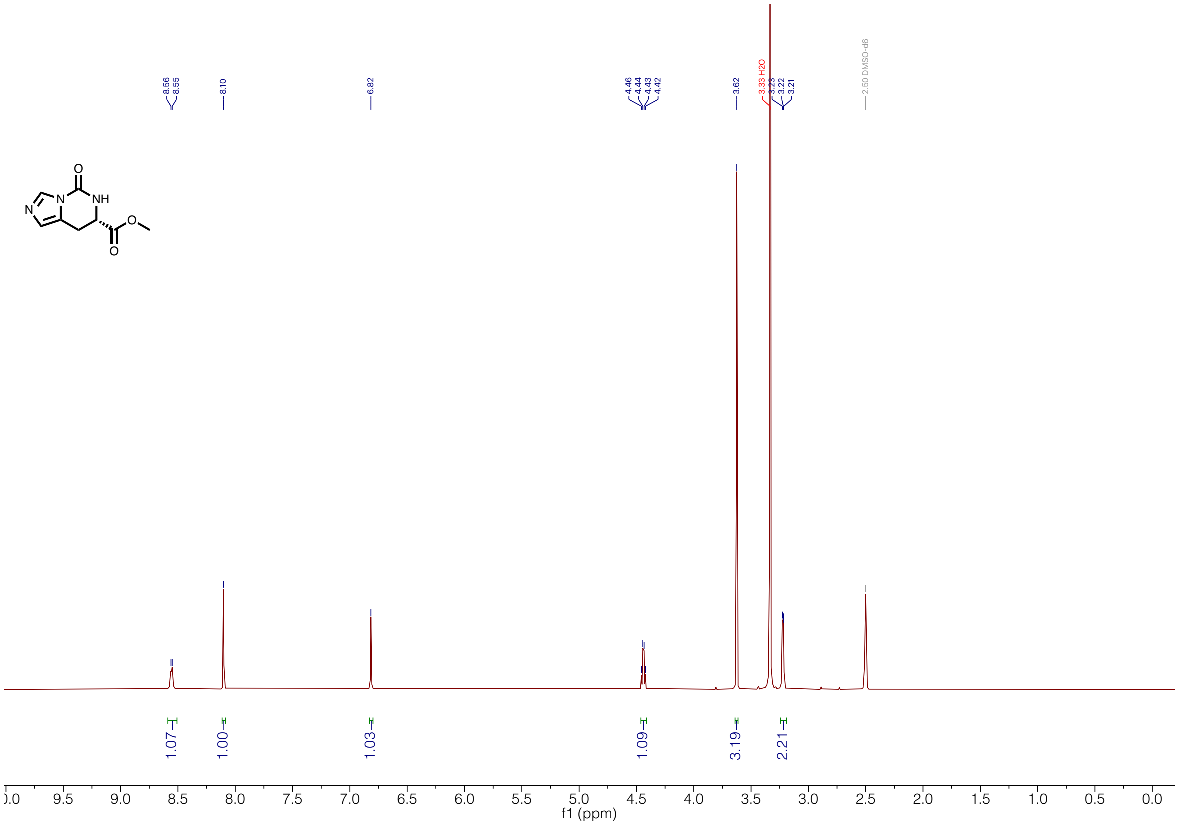


**Supplementary Figure 52 | $^{1}$H-NMR of methyl (*S*)-5-oxo-5,6,7,8-tetrahydroimidazo[1,5-*c*]pyrimidine-7-carboxylate.**

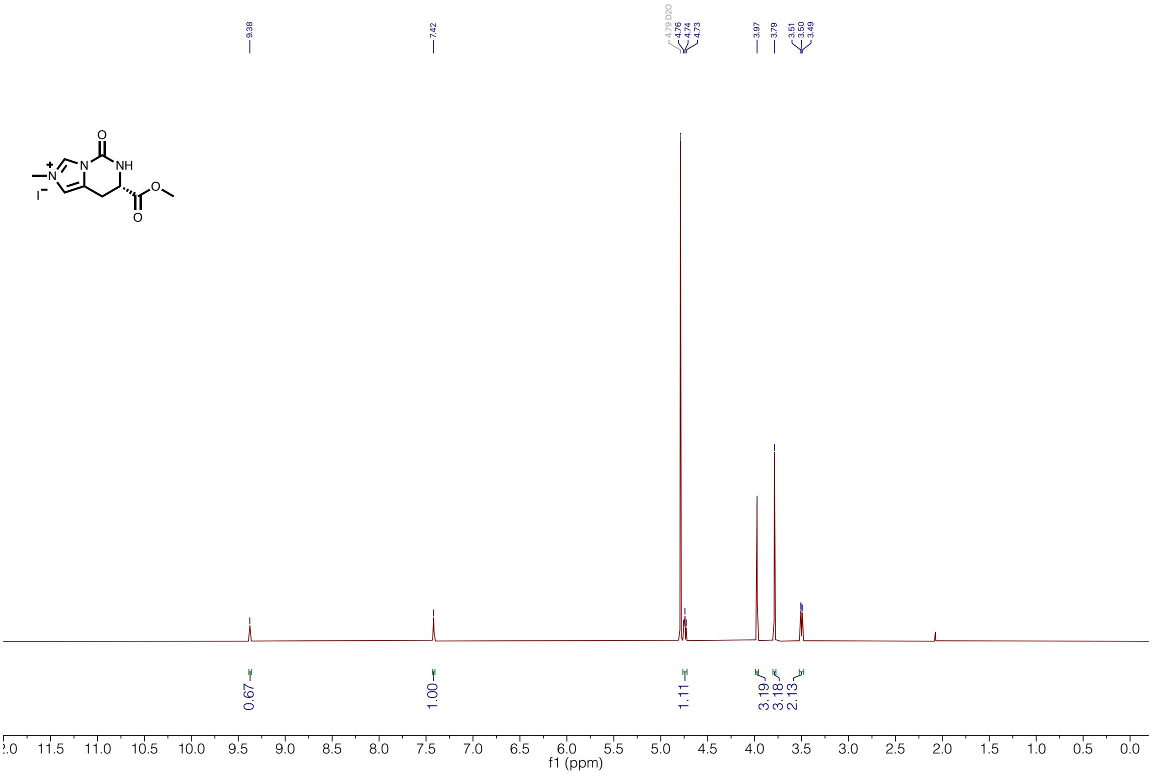


**Supplementary Figure 53 | 1H-NMR of (S)-7-(methoxycarbonyl)-2-methyl-5-oxo-5,6,7,8-tetrahydroimidazo[1,5-c]pyrimidin-2-ium iodide.**

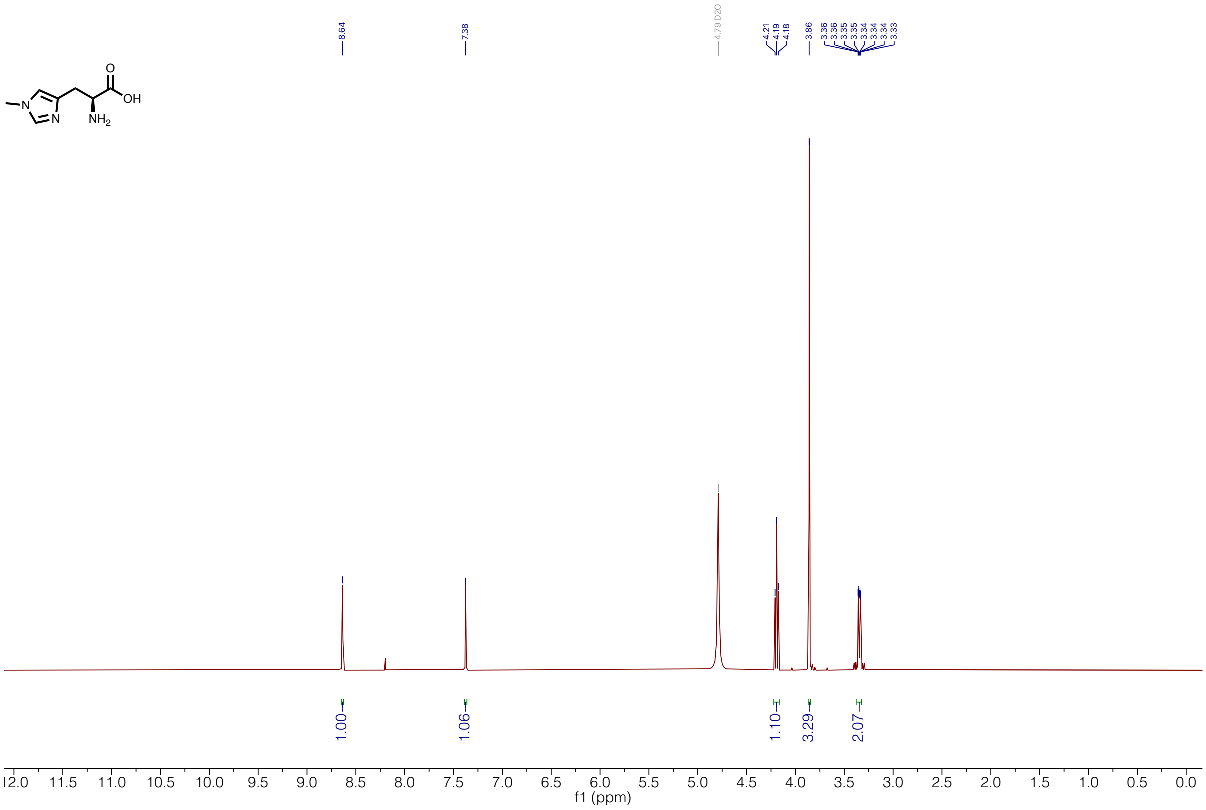


**Supplementary Figure 54 | $^{1}$H-NMR of *N*$^{\tau}$-methyl-L-histidine (τMH).**

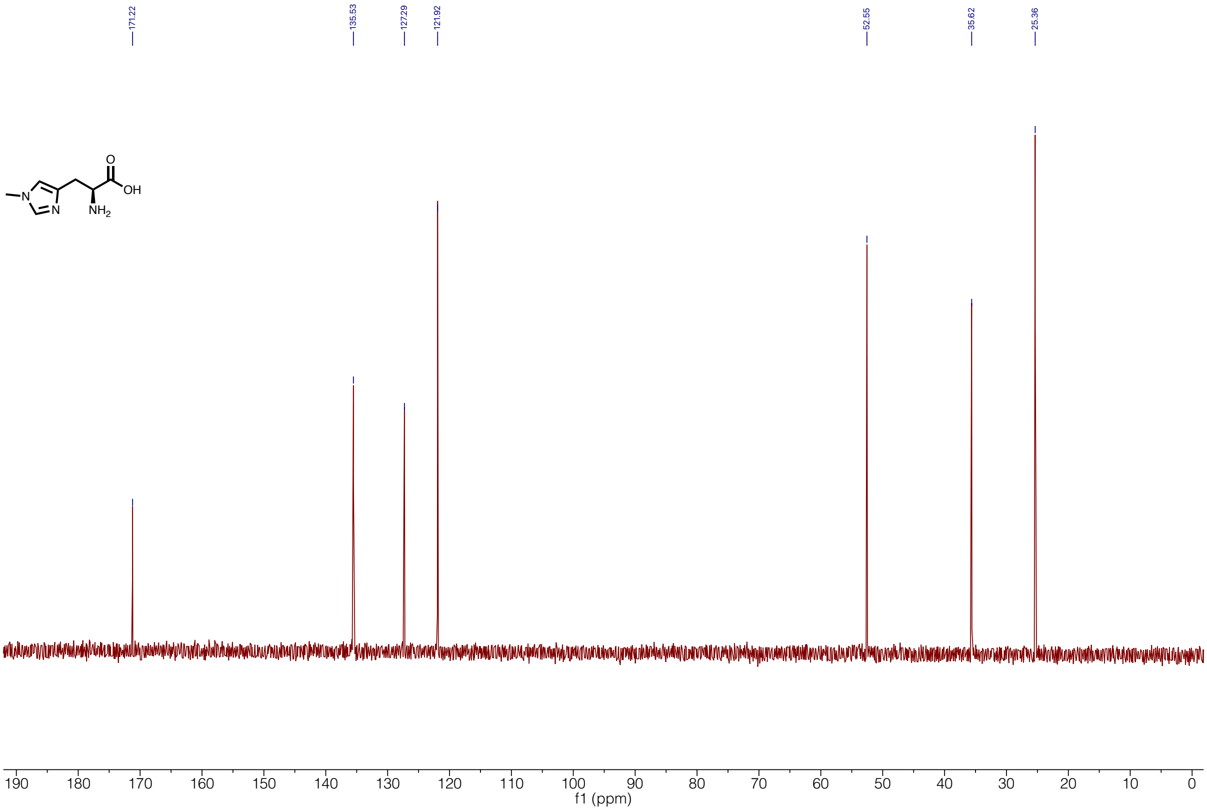


**Supplementary Figure 55 | $^{13}$C-NMR of *N*$^{\tau}$-methyl-L-histidine (τMH).**

**Oxazol-5-yl-L-alanine synthesis (5OxzA)**

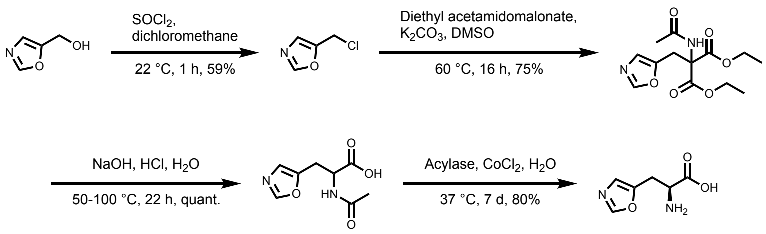


<u>(Oxazol-5-yl)methylchloride:</u>

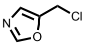

The reaction was performed under a nitrogen atmosphere. (Oxazol-5-yl)methanol (100 mg, 1.01 mmol, 1.0 eq.) was dissolved in dry dichloromethane (5 mL) and thienyl chloride (150 µL, 2.05 mmol, 2.0 eq.) was added. The solution was stirred for 1 h and the reaction was cooled to 4 °C. The reaction was quenched with aq. sat. $NaHCO_3$ (10 mL), the phases were separated, and the aqueous layer was extracted with EtOAc (3x). The combined organic phases were dried over $Na_2SO_4$ and the solvent was removed *in vacuo* to yield (oxaxol-5-yl)methanol (70 mg, 1.0 mmol, 59%) as a colorless oil.

$^1$H-NMR (400 MHz, $CDCl_3$) δ 7.90 (s, 1H), 7.11 (s, 1H), 4.63 (s, 2H).

<u>Diethyl-2-acetamido-2-(oxazol-5-ylmethyl)malonate:</u>

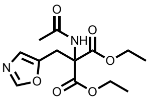

The following reaction was adapted from literature[64]. Diethyl acetamidomalonate (114 mg, 526 µmol, 1.0 eq.) was dissolved in DMSO (1.2 mL). (Oxazol-5-yl)methylchloride (68 mg, 579 µmol, 1.1 eq.) and $K_2CO_3$ (109 mg, 789 µmol, 1.5 eq.) were added and the suspension was stirred at 60 °C for 16 h. The reaction mixture was allowed to reach room temperature and ice-cold water (8 mL) was added. The mixture was extracted with EtOAc (2 x 10 mL), and the combined organic phase was washed with brine (10 mL). The organic phase was dried over $Na_2SO_4$ and the solvent was removed *in vacuo* to yield diethyl-2-acetamido-2-(oxazol-5-ylmethyl)malonate (118 mg, 526 mmol, 75%) as a white solid.

$^1$H-NMR (400 MHz, $CDCl_3$) δ 7.84 (s, 1H), 6.85 (s, 1H), 6.68 (s, 1H), 4.29 (q, $^3J_{HH}$ = 7.1 Hz, 4H), 3.82 (s, 2H), 2.03 (s, 3H), 1.29 (t, $^3J_{HH}$ = 7.1 Hz, 6H).

<u>*N*-Acetyl-oxazol-5-yl-alanine:</u>

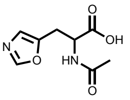

The following reaction was adapted from literature[68]. Diethyl-2-acetamido-2-(oxazol-5-yl-methyl)malonate (130 mg, 436 µmol, 1.0 eq.) was dissolved in water (7 mL) and NaOH (52 mg, 1.3 mmol, 3 eq.) was added. The solution was stirred at 50 °C for 16 h. The reaction mixture was adjusted to pH 4 with aq. HCl (4 M) and stirred at 100 °C for 6 h while adjusting the pH every few hours to pH 4. The solvent was removed *in vacuo* to yield *N*-acetyl-oxazol-5-yl-alanine (86 mg, 436 µmol, quant.) as a brown solid. The crude product was used for the next step without further purification.

Oxazol-5-yl-L-alanine (5OxzA):

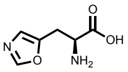

The following reaction was adapted from literature[68]. *N*-Acetyl-oxazol-5-yl-alanine (68 mg, 436 µmol, 1.0 eq.) was dissolved in water (1.2 mL) and $CoCl_2$ (0.27 mol%) was added. The mixture was adjusted to pH 7-8 and acylase from *Aspergillus genus* (1 mg) was added. The reaction mixture was stirred at 37 °C for 7 d and the solvent was removed *in vacuo* to yield an orange glue. The crude product was purified by automated flash column chromatography (C18, 1% acetonitrile in water (0.1% formic acid)) to yield oxazol-5-yl-L-alanine (27 mg, 174 µmol, 80%) as an off-white solid with formic acid as an impurity.

$^1$H-NMR (400 MHz, $D_2O$) δ 8.17 (s, 1H), 7.10 (s, 1H), 4.10 (dd, $^3J_{HH}$ = 6.4 Hz, $^3J_{HH}$ = 5.3 Hz, 1H), 3.47 – 3.35 (m, 2H).

$^{13}$C-NMR (101 MHz, $D_2O$) δ 173.0, 152.8, 147.2, 124.1, 53.2, 26.4.

MS (ESI): calc. for $[M+H]^+$: 157.05, obs.: 157.1.

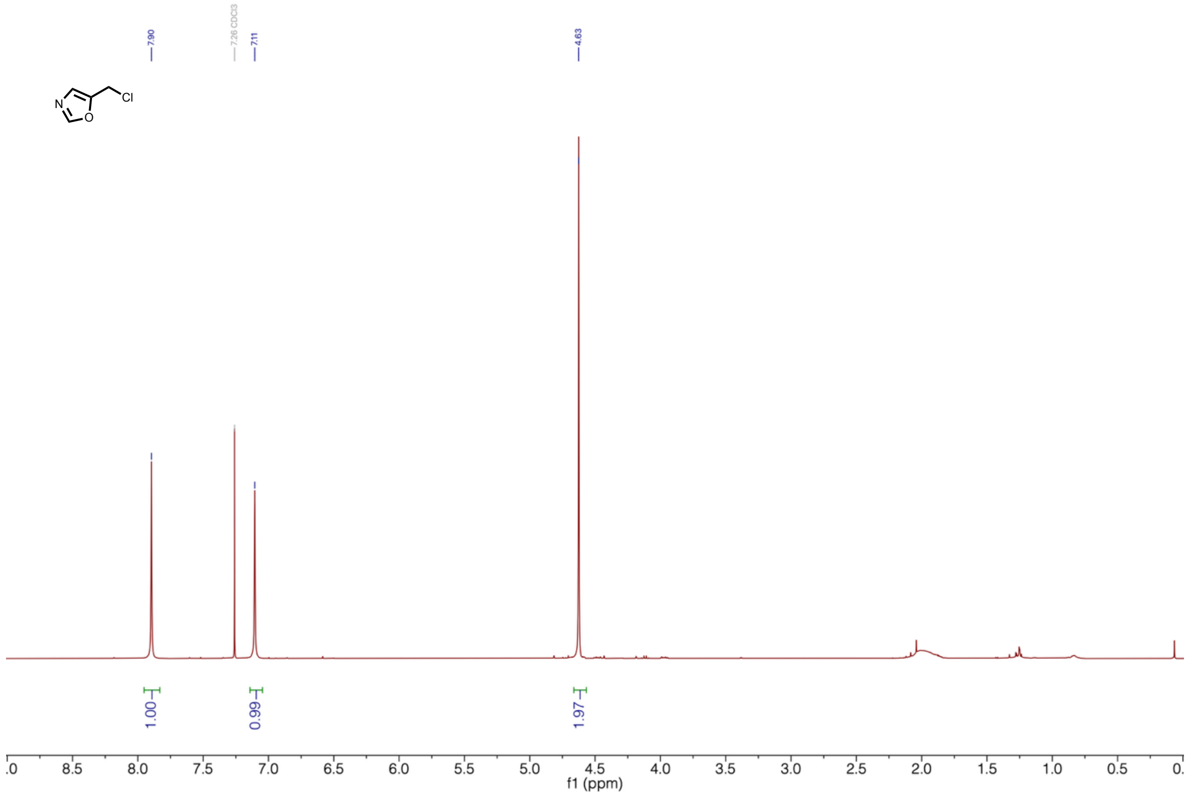


**Supplementary Figure 56 | $^1$H-NMR of (oxazol-5-yl)methylchloride.**

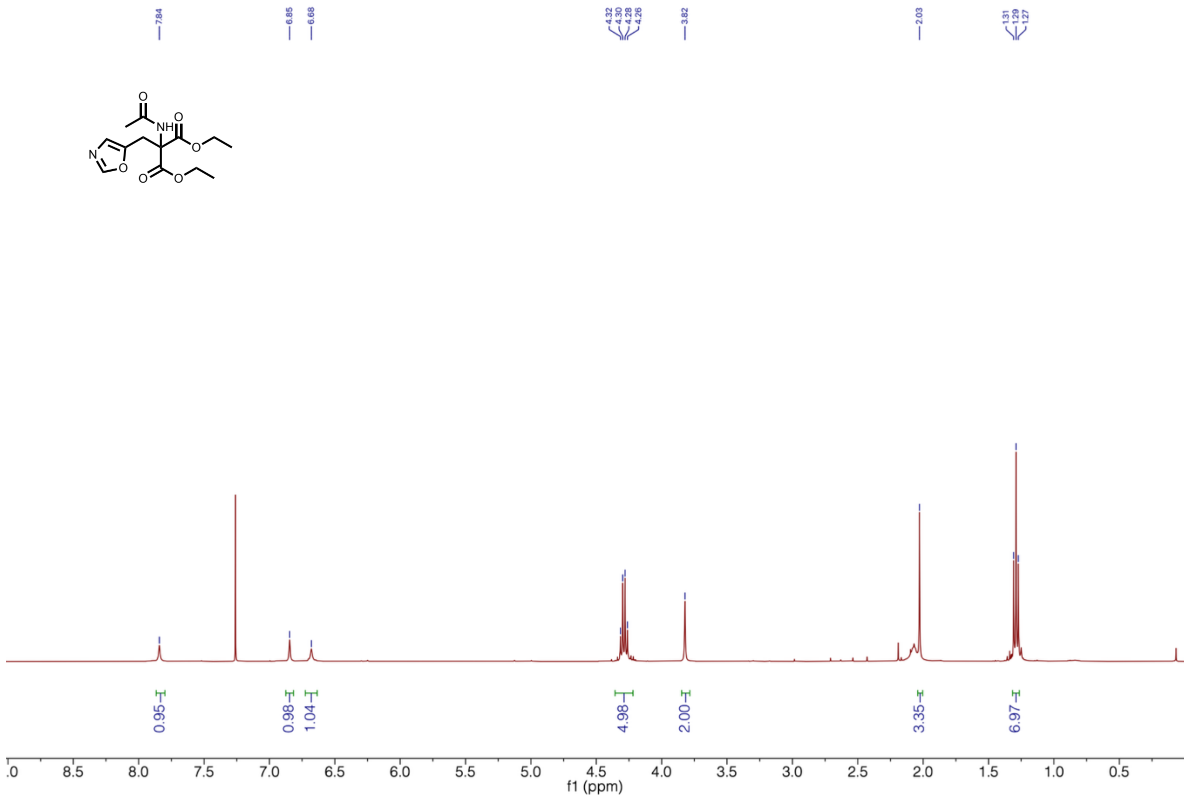


**Supplementary Figure 57 | $^1$H-NMR of diethyl-2-acetamido-2-(oxazol-5-ylmethyl)malonate.**

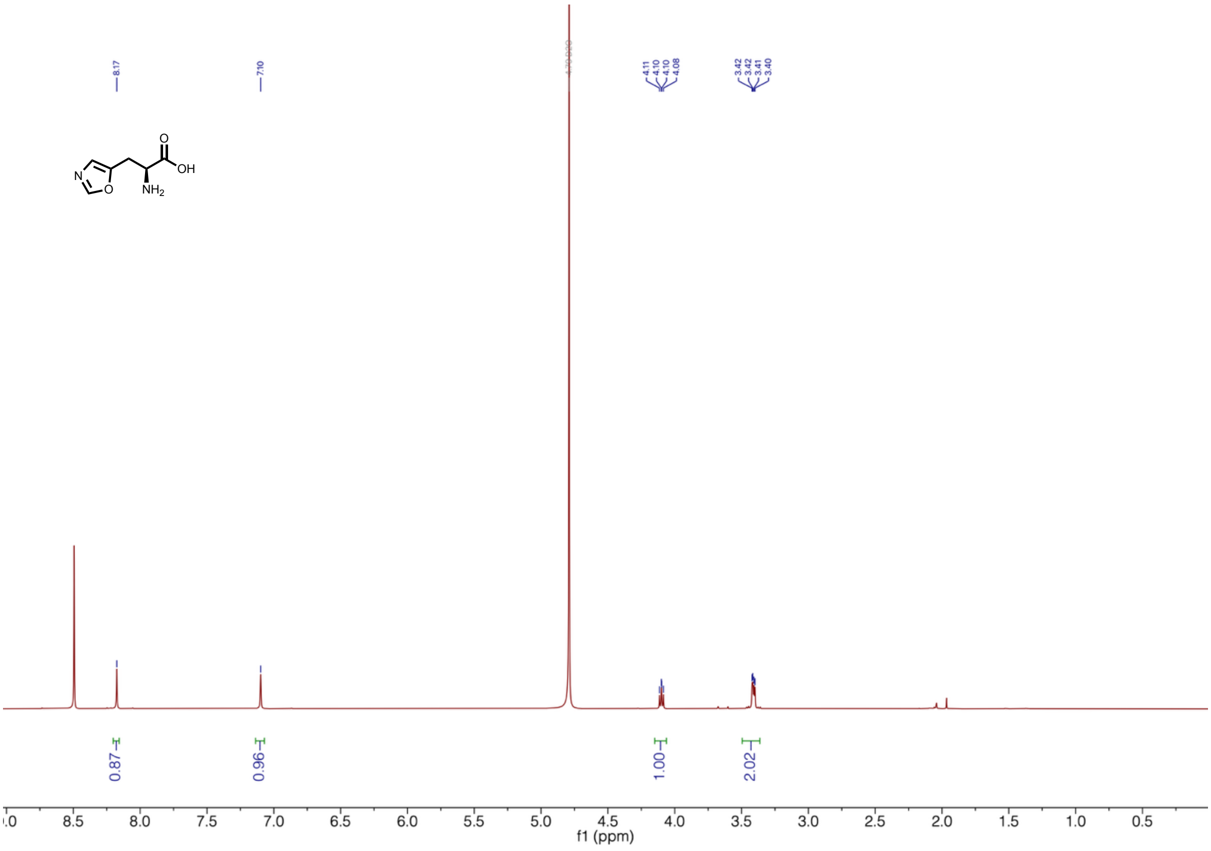


**Supplementary Figure 58 | $^1$H-NMR of oxazol-5-yl-alanaine (5OxzA)**

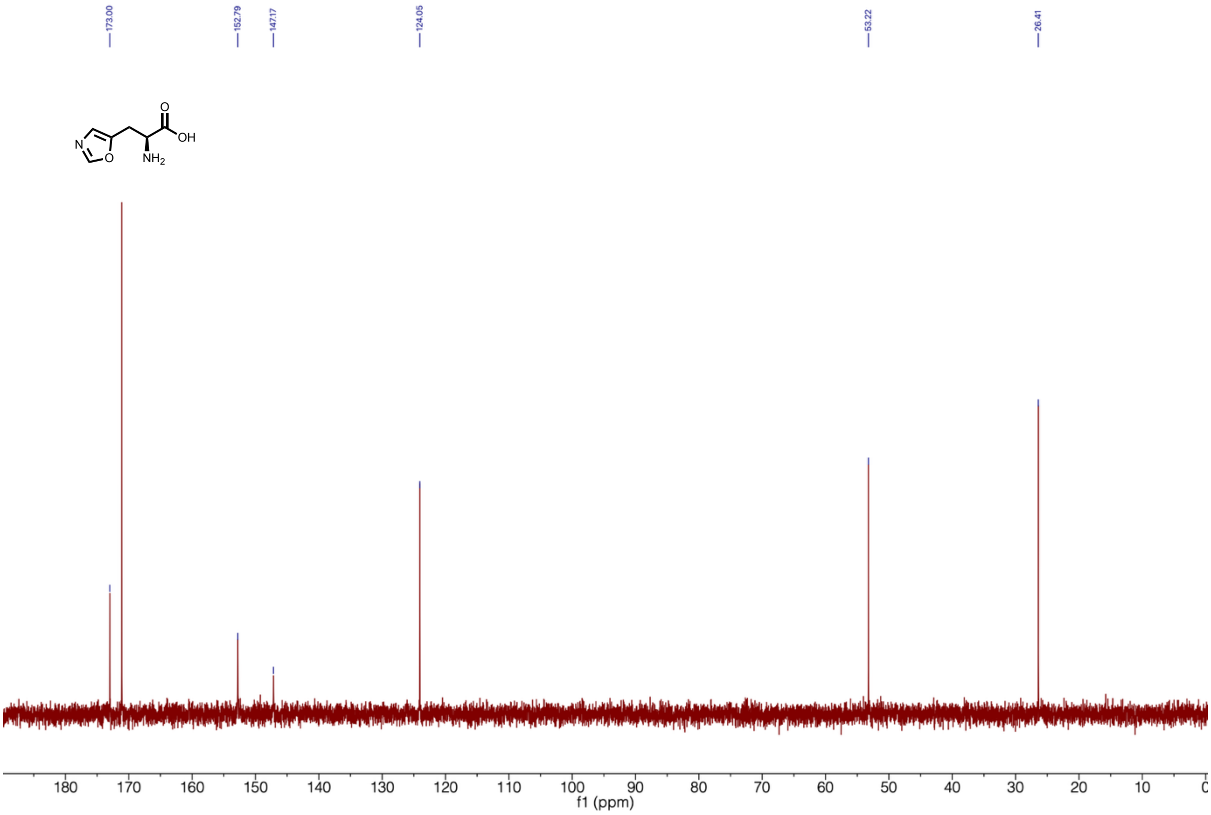


**Supplementary Figure 59 | $^{13}C$-NMR of oxazol-5-yl-alanaine (5OxzA).**
